# Adolescent stress recruits a latent amygdala-dopamine circuit to drive punishment-resistant reward-seeking

**DOI:** 10.64898/2026.07.29.741478

**Authors:** Jacob A. Nadel, Evan S. Swanson, Manhua Zhu, Baran Demir, Michelle H. Kwon, Sachin Patel, Talia N. Lerner

**Author notes:** Lead Contact: Talia N. Lerner, 303 E Chicago Ave, Ward 5-120, Chicago, IL 60611.

## Abstract

Adolescent stress is a lifelong risk factor for addiction, but the underlying neural circuit changes remain unknown. Here, we show that chronic unpredictable stress in adolescent mice causes a prominent increase in punishment-resistant reward-seeking – behavior tightly linked to the diagnostic criteria for addiction – establishing a model for mechanistic investigation. Using this model, we find a “gain-of-function” in reward processing wherein persistent hyperexcitability in a subset of central amygdala (CeA) neurons projecting to the substantia nigra pars lateralis (SNL) disinhibits dopamine release in the tail of the striatum (TS), newly recruiting TS to participate in punishment-resistant reward-seeking. Normalizing circuit-specific CeA hyperexcitability or optogenetically counteracting excessive TS dopamine release prevented the increase in punishment resistance. These results identify a novel circuit mediator for the lifelong effects of adolescent stress on a core feature of addictive disorders, opening a new avenue for interventions targeted to at-risk populations with specific formative experiences.

## Main Text

Adolescence is an evolutionarily-conserved developmental epoch marked by pronounced changes in physiology and behavior, including sexual maturation, increased exploratory and risk-taking behaviors, and widespread neurological remodeling^1–3^, with the scale of neurodevelopmental changes rivaling that of infancy^4^. The malleability of the adolescent brain provides unparalleled opportunities for adaptive learning as animals develop independence from caregivers; however, it also makes adolescence a period of vulnerability to perturbations such as stress^5,6^. Adolescents are physiologically more responsive to stress^7–11^, and under chronic stress, the adolescent brain is subjected to repeated, exaggerated activation of stress pathways, which can lead to dysregulation of stress responses and neural adaptations.^12,13^

One of the major consequences of chronic adolescent stress in humans is a predisposition towards addictive disorders later in life^14–17^. Addictive disorders are multifaceted, marked by deficits in impulse control^18^ and increases in punishment-resistant reward-seeking (the pursuit of the addictive substance or experience despite adverse consequences)^19,20^. While there has been extensive work on neural circuit alterations due to stress occurring during early life or adulthood^5^, significant gaps exist in understanding how chronic stress affects adolescent brain development. One important known effect is a transition towards habitual responding following social isolation in adolescence^21–23^. However, habitual responding and punishment-resistant reward-seeking are dissociable and may be subserved by distinct subcircuits^24–26^. Whether adolescent stress increases punishment resistant reward-seeking—the feature most directly tied to the diagnostic criteria for addictive disorders^20^—has not been directly tested.

An understanding of the specific circuits mediating the link between adolescent stress and lifelong addiction susceptibility is necessary to optimize treatments and patient outcomes. As psychiatry seeks to identify how social and environmental triggers for addiction and mental health disorders should influence diagnosis and treatment, animal models will be critical to pinpoint mechanisms underlying interactions between environmental triggers and individual neurobiology. These models offer the opportunity to precisely control the type of stress (e.g. social isolation, chronic unpredictable stress) and developmental timing. The identification of specific neural circuit changes following adolescent stress exposure will offer crucial insight for the treatment of at-risk populations based on known exposures to precipitating events.

Here, in mice, we show that chronic unpredictable stress in adolescence (but not adulthood) leads to a prominent, long-lasting increase in the propensity to develop punishment-resistant reward-seeking. We further demonstrate that this behavioral change is accompanied by and dependent upon a gain-of-function change in the central amygdala (CeA), a brain nucleus at the intersection of stress and addiction^27–30^. Hyperexcitability in a subset of CeA neurons projecting to the substantia nigra pars lateralis (SNL) is linked to potentiated dopamine release in the tail of the striatum (TS). The TS is a striatal subregion primarily studied in the context of threat processing in rodents^31^ but implicated in outcome-insensitive behavior in primates^32^. The result of potentiated TS dopamine release is a remarkable shift in the dopamine dependence of punishment-resistant reward-seeking from the dorsomedial striatum (DMS)^26^ to the TS.

### Chronic stress specifically in adolescence promotes punishment-resistant reward-seeking in mice

To model chronic adolescent stress in male and female mice, we applied a 12-day adolescent chronic unpredictable stress paradigm during early adolescence (P28 to P40)^5,33^ (Figure 1A). Mice received three distinct stressors (or combinations of stressors) in the morning, afternoon, and overnight (Supplementary Table 1). This paradigm led to sex-dependent changes in physiology; stressed male and female mice weighed less than unstressed littermate controls at P40, but females fully recovered by adulthood, while males remained slightly underweight (Figure S1A). Stressed mice had normal blood glucose levels in adulthood (Figure S1B), suggesting that any differences in sucrose-seeking during operant training are not due to changes in blood sugar levels. Adolescent stress blunted circulating corticosterone levels, particularly immediately after stress (Figure S1C), consistent with observations after adolescent social isolation^22^. Finally, consistent with previous reports^33^, adolescent stress increased anxiety-like behavior (Figure S1D-I).

**Figure 1.**
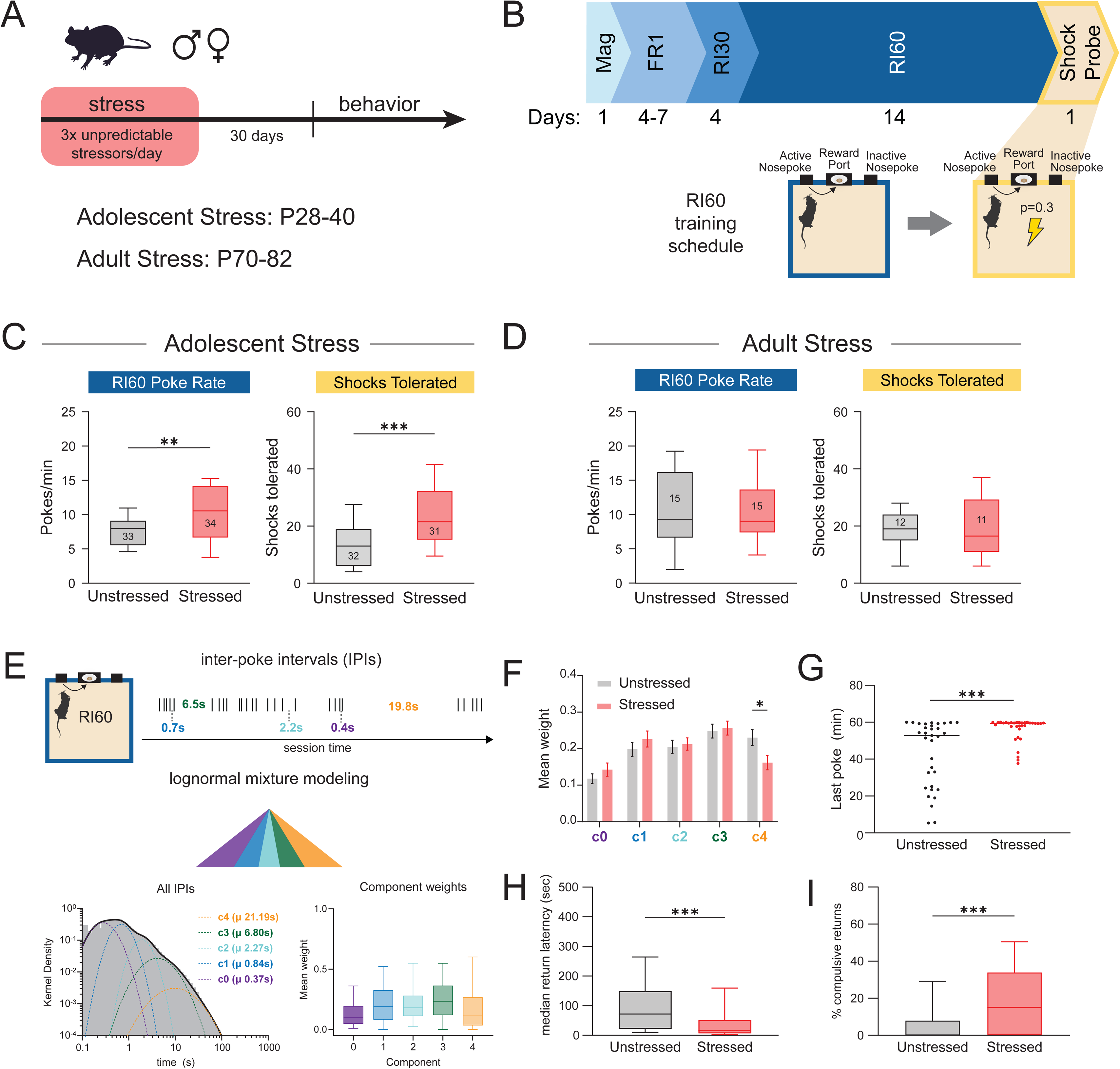
Chronic stress during adolescence but not adulthood increases punishment-resistant reward seeking. (A) Schematic of the chronic stress timeline. Adolescents were stressed from P28-40 and experiments began at P70, adults were stressed from P70-82 and experiments began at P112. (B) Top, schematic of RI60 behavioral paradigm: one day of magazine training (30 minute session with rewards delivered every 1 minute on average); 4-7 days of FR1 (every nosepoke on the active port was rewarded, mice reached criterion and moved to the next stage after 3 days with more than 30 rewards); 4 days of RI30 (random interval schedule, 30 seconds on average); 14 days of RI60 (random interval schedule, 60 seconds on average); one day of shock probe (FR1 reward delivery, RR3 shock delivery, time-out after 10 min without poke). Bottom, schematics of operant boxes during training and shock probes. (C) Left: Mice with a history of adolescent stress showed increased nosepoke rates during RI60 training (Welch’s t-test, **p = 0.0018. n = 33-34/group). Right: Mice with a history of adolescent stress tolerated an increased number of shocks in the shock probe (Mann-Whitney exact test, ****p < 0.0001, n = 31-32/group) (D) The same stress paradigm applied in adulthood did not produce changes in RI60 nosepoke rate (left; Mann-Whitney exact test, p = 0.8702, n = 15/group) or punishment-resistant reward-seeking (right; Unpaired t-test, p = 0.5119, n = 11-12/group) (E) Schematic of lognormal mixture modeling of inter-poke intervals (IPIs). A 5 component model was fit on IPI data and extracted 5 distinct modes of responding (from fastest to slowest): fast-motor (c0), engaged-fast (c1), engaged-slow (c2), between-bout (c3), and disengaged (c4). (F) Relative weights of each component in unstressed and stressed mice. Stressed mice spent less time in the disengaged state compared to unstressed controls (Bayesian Dirichlet regression, *p-analog = 0.022, n = 35/group) (G) Time of last nosepoke during the shock probe. Unstressed mice renounced nosepoking during the shock probe more frequently and faster than stressed mice (Mann-Whitney exact test, ***p = 0.0004, n = 33/group) (H) Median latency to return to nosepoking during the shock probe after receiving a shock. Stressed mice returned to nosepoking faster than unstressed controls (Mann-Whitney exact test, ***p = 0.0005, n = 34-36/group). (I) Percent compulsive returns (nosepokes performed within 5s of being shocked). Stressed mice performed compulsive returns after a larger percentage of shocks than unstressed mice (Mann-Whitney exact test, ***p = 0.0001, n = 34-36/group) C-D, H-I are presented as box-and-whisker plots with a line at median, box at 25-75 percentile, whiskers at most extreme point within IQR x 1.5 (Tukey method). Outliers not shown. F is presented as mean ± SEM. G is presented with a line at median. ^∗^p < 0.05; ^∗∗^p < 0.01; ^∗∗∗^p < 0.001; ^∗∗∗∗^p < 0.0001. n vary between test primarily due to outlier removal, see Methods. See also Table S2 for detailed statistics

To test whether adolescent stress increased the propensity for punishment-resistant reward-seeking, we trained adult mice (beginning at P70; 30 days after the conclusion of the stress paradigm) on a random interval operant reinforcement paradigm (RI60) for 14 days, and tested for punishment-resistant reward-seeking in a shock probe at the end of training^26^. In the shock probe session, reward-seeking nosepokes resulted in reward as well as a one-third risk of mild foot shock (1s, 0.2 mA; Figure 1B). The session timed out if mice did not nosepoke for 10 minutes. Importantly, adolescent stress did not alter learning in a two-chamber active avoidance paradigm (Figure S1J-L), indicating that any adolescent stress-induced changes in punishment-resistant reward-seeking are not due to changes in shock tolerance or the ability of mice to learn about adverse consequences.

During RI60 training, adolescent stress mice had higher RI60 nosepoke rates than unstressed littermate controls regardless of sex (Figure 1C left, Figure S2A), with rates increasing over training (Figure S2B). There were no adolescent stress-induced changes in the rate of entry into the central reward port, leading to an increased ratio of nosepokes per port entry in adolescent stress mice (Figure S2D-E). Despite increased nosepoking, due to the nature of the RI60 task, adolescent stress mice did not earn more rewards (Figure S2F). Thus, adolescent stress mice performed the task less efficiently than unstressed controls, earning fewer rewards per nosepoke (Figure S2G).

Elevated nosepoke rates could be explained by a variety of factors, including increased vigor, fine-grained bout restructuring, decreased disengagement, decreased rate of satiety, or global hyperactivity. To disambiguate these variables, we fit a five component hierarchical lognormal mixture model to pooled inter-poke intervals (IPIs) during RI60 training (Figure 1E, Figure S3A-B). The five-component model identified distinct components of the IPI distribution, which we labeled as fast-motor (c0), engaged-fast (c1), engaged-slow (c2), between-bout (c3), and disengaged (c4) (Figure 1E). Within this framework, increased vigor would manifest as increased weight on c0-1, bout restructuring would manifest as increased nosepokes per bout, disengagement as increased weight on c4, decreased rate of satiety as shallower slope of disengagement (c4 weight) across the session and hyperactivity as a lack of weight change (as global hyperactivity would increase poke rate across all components). Group weights did not differ for c0-3, ruling out increased vigor (Figure 1F), and bout microstructure did not change (Figure S3C). However, adolescent stress mice spent significantly less time in the disengaged state (c4), also reflected as decreased inter-bout interval (Figure S3D). Finally, disengagement slope across the session did not differ across groups, suggesting similar satiation rates between groups (Figure S3E-F). We concluded that adolescent stress mice have higher RI60 nosepoke rates due to continuously high levels of engagement throughout each session.

During the shock probe session, stressed mice tolerated more shocks than unstressed controls (Figure 1C right, Figure S2C). Nosepoke rates were elevated throughout the entire shock probe session in adolescent stress mice, demonstrating long-lasting persistence in reward-seeking under threat of punishment (Figure S2H). Nosepoke behavior decayed at the same rate across groups (Figure S2I), but unstressed mice timed out earlier (Figure 1G), indicating that stressed mice had a higher threshold for quitting. Adolescent stress mice also returned to poking at a significantly shorter latency post-shock compared to unstressed controls (Figure 1H) and were much likelier to return to poking within 5 seconds of a shock (compulsive returns, Figure 1I). Shocks tolerated and poke rate during training were not correlated in either group, indicating the observed increase in RI60 poke rate was not related to the observed increase in punishment-resistance (Figure S2J). Together, these results demonstrate that mice with a history of chronic stress in adolescence show increased punishment-resistant reward-seeking in adulthood, coupled with other markers of compulsive engagement in reward-seeking.

Stress can have markedly different behavioral effects depending on the developmental timepoint at which the stress is experienced^33^. To investigate if the effect of stress on punishment resistance is developmentally specific to adolescence, we applied the same 12-day CUS protocol to adult mice (P70-P82), then trained these mice and unstressed littermate controls on the same RI60 task starting 30 days later (P112; Figure 1A). In contrast to the adolescent stress group, male and female mice that underwent stress in adulthood did not differ from unstressed controls in poke rate or number of shocks tolerated (Figure 1D, Figure S2K-M). Groups also did not differ in entry rate, pokes per entry, efficiency, or reward rate (Figure S2N-Q). These data demonstrate that adolescence is a particularly vulnerable time point where chronic stress results in long-lasting increases in susceptibility to punishment-resistant reward-seeking. Given the developmental specificity of stress effects, we focused specifically on adolescence for the remaining experiments.

### Adolescent stress leads to projection-specific changes in intrinsic properties of central amygdala neurons

Extensive evidence indicates that the central amygdala (CeA) plays a central role in stress processing and is dramatically affected by chronic stress throughout the lifespan^28,34–37^. There is also plentiful evidence that the CeA is involved in addiction and addictive-like behaviors^27,29,38,39^, including punishment resistance^27^. Thus, the CeA is in a prime position to mediate the relationship between adolescent stress and punishment-resistance in adulthood. However, the CeA is a heterogenous structure that projects to many downstream targets^40^. We examined the effects of adolescent stress on the physiological properties of three major projection-defined subpopulations of CeA neurons with minimal cross-collateralization^40,41^, namely the CeA populations projecting to the bed nucleus of the stria terminalis (BNST), periaqueductal gray (PAG), and substantia nigra pars lateralis (SNL). We injected red retrobeads into the BNST, PAG, or SNL of adolescent stress mice and littermate controls and recorded intrinsic properties in current clamp starting at P70 (Figure 2A-C). Recordings were performed in the presence of glutamatergic and GABAergic blockers (D-AP5, NBQX, and GBZ) to isolate intrinsic excitability (see Methods). All electrophysiology data were analyzed using hierarchical mixed-effects modeling to account for the nested structure of the data (multiple cells within individual mice).

**Figure 2.**
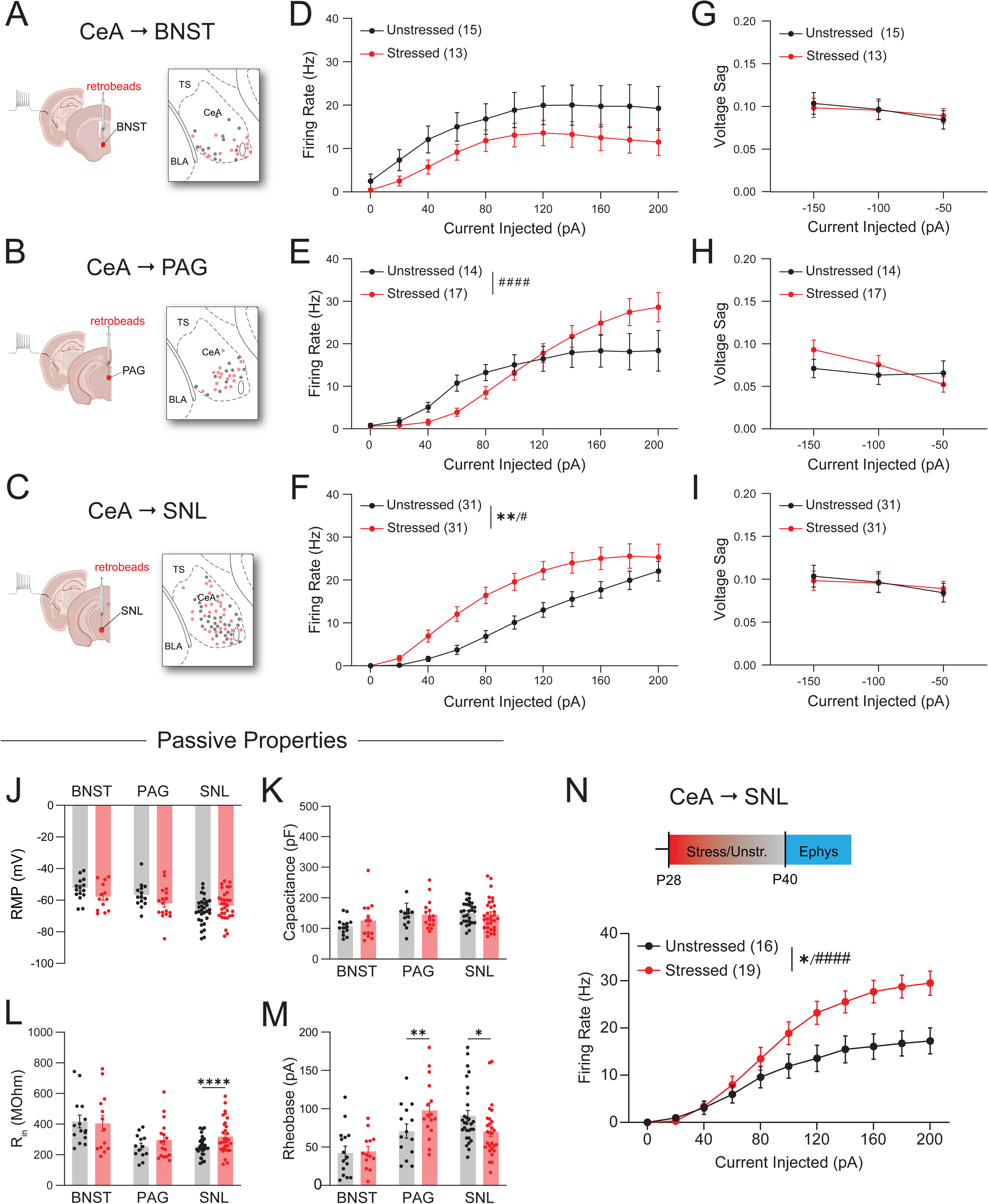
Adolescent stress alters the intrinsic properties of CeA neurons in a projection-specific manner. (A-C) Experimental schematics and maps of recording locations for CeA neurons that project to the BNST (A), PAG (B), or SNL (C). Red retrobeads were injected in the target output region for retrograde tracing and intrinsic properties of bead positive CeA cells were recorded using whole-cell patch-clamp electrophysiology. TS = tail of the striatum, BLA = basolateral amygdala, CeA = central amygdala. Gray dots show locations of CeA neurons recorded from unstressed mice; red dots show locations of CeA neurons recorded from stressed mice. (D-F) Firing rates in response to increasing depolarizing current injection in CeA→BNST (D, current: ****p < 0.0001, stress: p = 0.1753, interaction: p = 0.9714), CeA→PAG (E, current: ****p < 0.0001, stress: p = 0.6254, interaction: ^####^p < 0.0001) and CeA→SNL (F, current: ****p < 0.0001, stress: **p = 0.0037, interaction: ^#^p = 0.0165) neurons. Stress did not significantly affect the CeA→BNST projection, but led to complex effects in the CeA→PAG projection, and broad hyperexcitability in the CeA→SNL projection. (G-I) Voltage sag in response to hyperpolarizing current injections. There were no effects of adolescent stress on voltage sag properties in CeA neurons projecting to BNST (G, current: ****p < 0.0001, stress: p = 0.9399, interaction: p = 0.7305), PAG (H, current: **p = 0.0094, stress: p = 0.6037, interaction: p = 0.0521), or SNL (I, current: ***p = 0.0009, stress: p = 0.7775, interaction: p = 0.4106). (J) No effects of stress on resting membrane potential (RMP) in any studied projection (BNST, p = 0.124; PAG, p = 0.064; SNL, p = 0.075) (K) No effects of stress on capacitance in any studied projection (BNST: p = 0.114; PAG: p = 0.332; SNL: p = 0.590) (L) Stress increased input resistance (R_in_) in CeA→SNL neurons (BNST: p = 0.937; PAG: p = 0.118; ****SNL: p < 0.0001) (M) Stress increased rheobase (current required to evoke a spike within 50 ms) in CeA→PAG neurons and decreased rheobase in CeA→SNL neurons (BNST: p = 0.419; PAG: **p = 0.003; SNL: *p = 0.025) (N) CeA→SNL hyperexcitability was already present in recordings from CeA→SNL neurons made at P40, immediately after the end of stress (current: ****p < 0.0001, stress: *p = 0.0218, interaction: ^####^p < 0.0001) All data presented as mean ± SEM. For all P70 experiments, BNST n = 3 mice, 13-15 cells/group; PAG n = 2-3 mice, 14-17 cells/group; SNL n = 5-6 mice, 29-31 cells/group. For P40 experiment, n = 2 mice, 16-19 cells/group. All data analyzed with hierarchical linear mixed effects models. ^∗^*p* < 0.05; ^∗∗^*p* < 0.01; ^∗∗∗^*p* < 0.001; ^∗∗∗∗^*p* < 0.0001 for main effects or pairwise comparisons; ^#^*p* < 0.05; ^##^*p* < 0.01; ^###^*p* < 0.001; ^####^*p* < 0.0001 for interactions. See also Table S2 for detailed statistics

Excitability was assessed using depolarizing current injections. In BNST-projecting neurons, stress caused a non-significant trend towards hypoexcitability (Figure 2D). In PAG-projecting neurons, stress caused an increased dynamic range of firing with hypoexcitability at lower currents (higher rheobase) and hyperexcitability at higher currents (Figure 2E). Finally, in SNL-projecting neurons, stress caused hyperexcitability across current injection levels (Figure 2F). Analyses of spike train properties showed that stress had subtle effects on spike adaptation rates in CeA→BNST neurons (Figure S4A-C) but led to a higher latency to fire and increased mean inter-spike intervals (ISIs) in CeA→PAG neurons (Figure S4D-F). The most consistent impacts were in CeA→SNL neurons, with stress impacting latency to fire, spike rate adaptation, and mean ISI (Figure S4G-I). Hyperpolarizing current injections did not reveal any stress-induced differences in voltage sag ratios, indicating adolescent stress did not change I_h_ (hyperpolarization activated current) in any projection (Figure 2G-I). We did not observe stress-induced changes in resting membrane potential (RMP) or capacitance across projections (Figure 2J-K). CeA→SNL neurons from stressed mice had higher input resistance (Figure 2L) and lower rheobase (Figure 2M), while CeA→PAG neurons had higher rheobase (Figure 2M). Altogether, adolescent stress had the strongest effects on CeA→SNL neurons. Notably, the hyperexcitable phenotype in CeA→SNL neurons had already emerged by the end of the adolescent stress paradigm (P40; Figure 2N). We concluded that CeA→SNL hyperexcitability is a persistent neural change induced by adolescent stress with the potential to underlie the observed change in punishment-resistant reward-seeking.

Given previous reports of distinct electrophysiological subtypes of CeA neurons^42^, we further investigated heterogeneity within the CeA→SNL population using PCA and k-means clustering (see Methods). After regressing out the effects of stress, we selected three k-means clusters, and found neurons belonging to each of the three widely observed non-spontaneous electrophysiological subtypes in the CeA (Figure 3A-B)^42,43^: regular-spiking (cluster 1), low threshold bursting/adapting (cluster 2), and late firing (cluster 3; Figure 3D). All three clusters had similar proportions of stressed and unstressed cells (Figure 3C) and displayed signs of hyperexcitability following adolescent stress (Figure 3E-J). Although passive properties differed between clusters, markers of hyperexcitability (input resistance, rheobase) displayed main effects of stress, indicating a common, adolescent stress-induced shift towards hyperexcitability.

**Figure 3.**
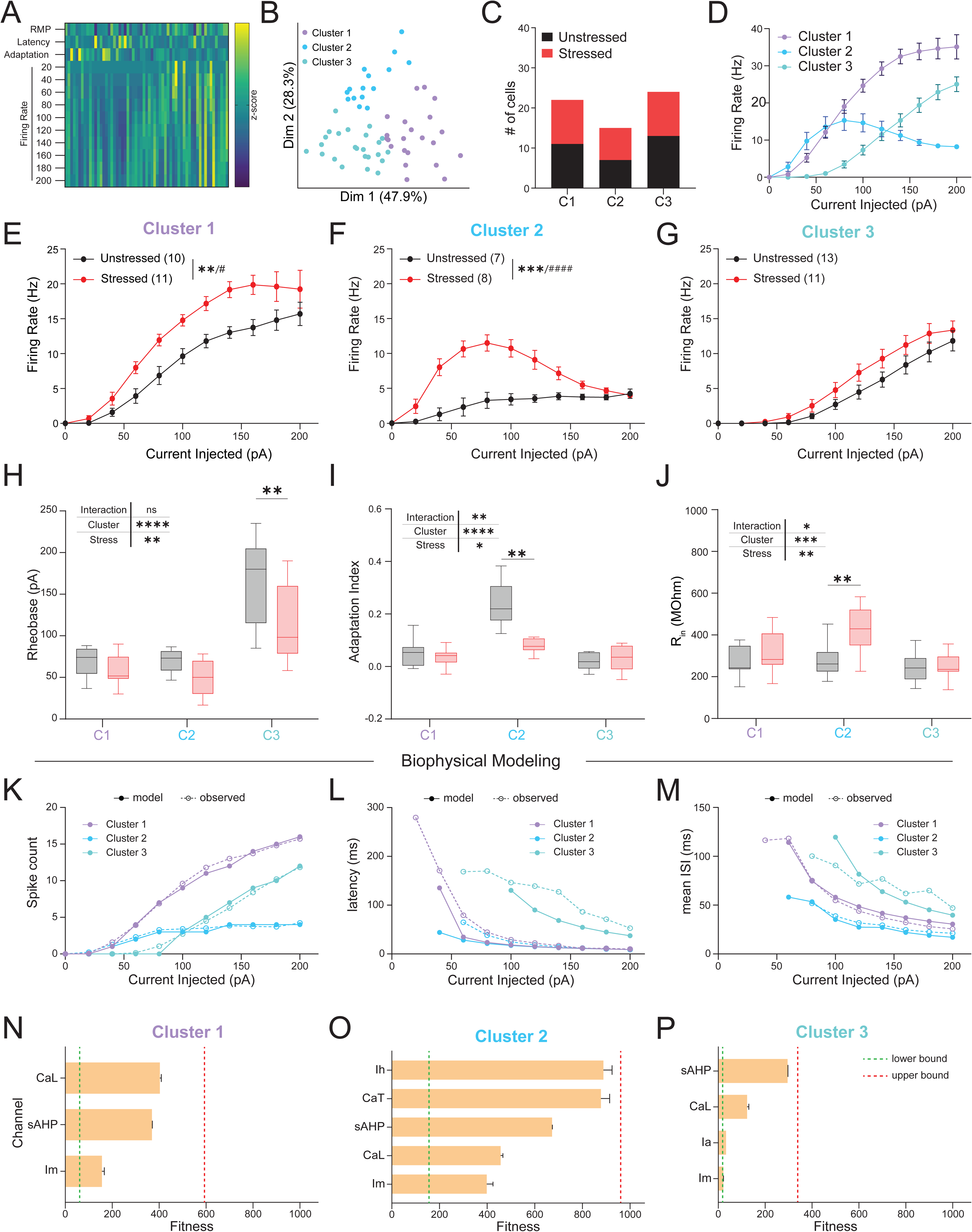
Adolescent stress induces hyperexcitability across three electrophysiologically distinct CeA→SNL subtypes. (A) Heat map showing z-scored features used for clustering CeA→SNL neurons into electrophysiological subtypes: resting membrane potential (RMP), mean latency to fire across currents (Latency), mean adaptation index across currents (Adaptation), and firing rate at each current injection level. Each column represents a recorded neuron. (B) PCA analysis showing 3 identified cell clusters projected onto the first two principal components of the 4-component PCA. (C) Number of neurons from stressed and unstressed mice assigned to each cell cluster. Each cluster contained nearly equivalent numbers of stressed and unstressed cells (C1: 11 stressed/10 unstressed; C2: 8 stressed/7 unstressed; C3: 11 stressed/13 unstressed). (D) Mean firing rates of each cluster, shown with stressed and unstressed cells combined. The firing properties of these clusters closely match previously described electrophysiological subtypes within the CeA: regular spiking (cluster 1), low-threshold bursting (cluster 2), and late firing (cluster 3). (E-G) Firing rates of each cluster split by stress condition. Clusters 1 (E, stress: **p = 0.0014, ****current: p < 0.0001, interaction: ^#^p = 0.0162; n = 10-11 cells/group) and 2 (F, stress: ***p = 0.0004, current: ****p < 0.0001, interaction: ^####^p < 0.0001; n = 7-8 cells/group) displayed increased firing rates in response to depolarizing currents; Cluster 3 was not statistically significant (G, stress: p = 0.1138, current: ****p < 0.0001, interaction: p = 0.1634; n = 11-13 cells/group) (H) Rheobase shown by cell cluster and stress condition. Stress led to decreased rheobase across clusters that is particularly potent in cluster 3 (stress: **p = 0.0025, ****cluster: p < 0.0001, interaction: p = 0.2607; post-hoc: C1, p = 0.5959, C2, p = 0.5807, C3, **p = 0.0033; n = 7-13 cells/cluster/group) (I) Adaptation index shown by cell cluster and stress condition. Stress led to a decrease in adaptation index driven by cluster 2 (stress: *p = 0.0305, ****cluster: p < 0.0001, ^##^interaction: p = 0.0025; post-hoc: C1, p = 0.9584, **C2, p = 0.0012, C3, p = 0.6577; n = 7-13 cells/cluster/group) (J) Input resistance shown by cell cluster and stress condition. Stress led to an increase in input resistance, driven by cluster 2 (stress: **p = 0.0023, ***cluster: p = 0.0004, interaction: ^#^p = 0.0498; post-hoc: C1, p = 0.4519, C2, **p = 0.0037, C3, p = 0.9796; n = 7-13 cells/cluster/group) (K-M) Comparisons between observed kdata (dashed lines) and model output (solid lines) for spike count (K), latency to fire (L), and mean inter-spike interval (ISI; M). (N-P) The single conductances that most closely recapitulated observed stressed data (when all other parameters were left frozen at unstressed fit) are shown for cluster 1 (N), cluster 2 (O), cluster 3 (P). Across all clusters, M-type potassium currents (I_m_) were the closest fit. Mean + SEM of 10 replicate runs. The green dotted lines mark fitness on stress data when all values were allowed to vary (lower bound); the red dotted lines mark fitness on stress data when all values were frozen at unstressed fits (upper bound). CaL: L-type Ca^2+^ current; sAHP: slow afterhyperpolarization; I_m_: M-type K^+^ current; I_h_: hyperpolarization-activated current; CaT: T-type Ca^2+^ current; I_a_: A-type K^+^ current Linear mixed-effects models were used for all analyses. D-G are presented as mean ± SEM. H-J are presented as box and whisker plots (IQR, median line, Tukey whisker method, points outside whiskers not shown). ^∗^*p* < 0.05; ^∗∗^*p* < 0.01; ^∗∗∗^*p* < 0.001; ^∗∗∗∗^*p* < 0.0001 for main effects or pairwise comparisons; ^#^*p* < 0.05; ^##^*p* < 0.01; ^###^*p* < 0.001; ^####^*p* < 0.0001 for interactions. See also Table S2 for detailed statistics.

The three CeA→SNL clusters differ substantially in baseline electrophysiological properties, yet adolescent stress led to hyperexcitability in all three. This convergence raises the possibility that a shared conductance underlies the effect. To explore the potential mechanisms causing hyperexcitability following adolescent stress, we modeled each electrophysiological CeA subtype using NEURON^44^, basing our channel kinetics and initial parameters on previous modeling^45^. Parameters were fit to control experimental data using a Covariance Matrix Adaptation – Evolution Strategy (CMA-ES) algorithm, which has been shown to accurately fit ionic conductances based on current clamp data (Figure S5A)^46^. This approach produced accurate fits for each cluster, with model predictions closely approximating input-output curves, latencies to fire, and mean ISI values (Figure 3K-M).

We then asked if any single conductance could account for the stress-induced shifts in each cluster. To identify candidate conductances causing stress-induced hyperexcitability in each cluster, we re-ran the model against the stressed data in three ways: first, we set the upper bound of error by freezing all parameters at the unstressed values; second, we set the lower bound of error allowing all parameters to vary; and third, we systematically varied single channel conductances while keeping all other parameters frozen (Figure S5B-C). The bounds establish the unstressed-to-stressed gap that individual conductances need to close. Using this approach, we identified the best single candidates accounting for stress-induced changes in each cluster. Strikingly, the M-type potassium current (I_m_) was the best performing single channel for all clusters (Figure 3N-P). I_m_ is a subthreshold voltage-gated outward current carried by KCNQ/Kv7 channels, and decreased I_m_ has been implicated in CeA hyperexcitability related to hypertension^47,48^. Consistent with I_m_ being available to modulate the observed projection-specific hyperexcitability of the CeA→SNL population, re-analysis of single-cell RNA-seq data from CeA^40^ indicates that putative SNL-projecting CeA neurons (Drd1^+^/Sema3c^−^) have higher Kcnq2/3 expression compared to putative BNST-projecting (Dlk1^+^) or PAG-projecting (Crh^+^) neurons (Figure S5D). This further implicates KCNQ channels as a plausible substrate for CeA→SNL hyperexcitability. These results provide a potential launching point for molecular investigations into the role of KCNQ channels in mediating the effects of adolescent stress.

### Reward-related potentiation of dopamine release in the tail of the striatum (TS) causes punishment-resistant reward-seeking following adolescent stress

We next investigated how CeA→SNL hyperexcitability might influence downstream circuits. Although the SNL contains both GABAergic and dopaminergic neurons, previous evidence indicates a stronger projection of CeA neurons onto SNL GABAergic neurons, which has the net effect of disinhibiting dopaminergic neurons^41^. SNL dopamine neurons project to the tail of the striatum (TS)^49^, a striatal subregion implicated in threat and sensory processing in rodents^31,50^ and outcome insensitive habits in primates^32,51^. Given that adolescent stress induced CeA→SNL hyperexcitability (Fig. 2) and that activation of the CeA→SNL projection disinhibits SNL dopamine neuron activity^52^, we hypothesized that adolescent stress would potentiate TS dopamine release during RI60 training. To test this hypothesis, we performed TS-targeted in vivo fiber photometry recordings with the dopamine sensor GRAB-DA2m^53^. We compared dopamine dynamics during RI60 training in adolescent stress mice to unstressed controls (Figure 4A-B, Figure S6A).

**Figure 4.**
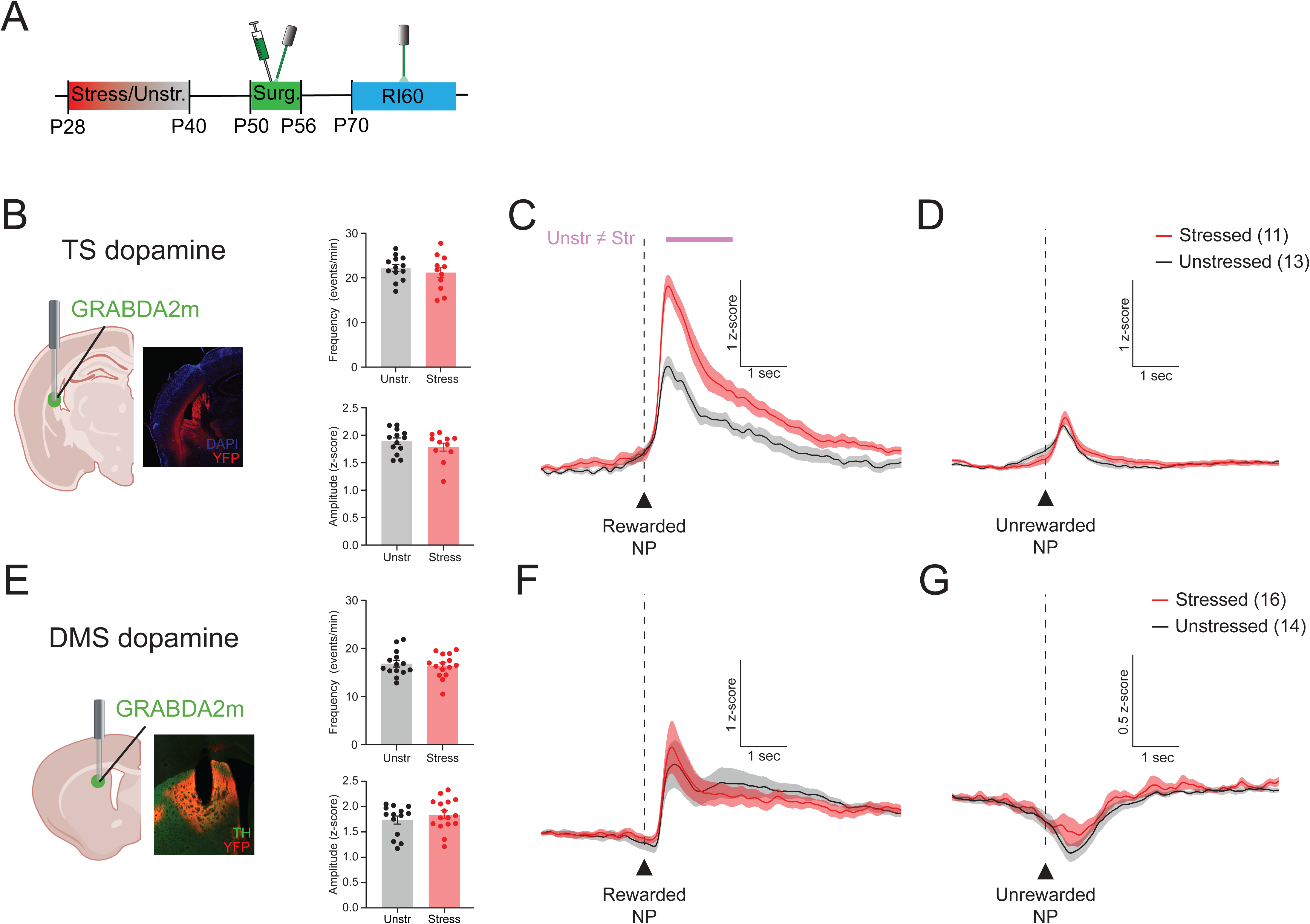
Mice with a history of adolescent stress have potentiated reward-related dopamine release in the TS but not DMS. (A) Schematic of experimental design. After adolescent stress (P28-40), mice were injected with GRAB-DA2m and implanted with fibers over the TS or DMS, then began RI60 training at P70. (B) Left, Schematic of photometry recording strategy for the TS. Right, Overall amplitude and frequency of TS dopamine transients did not differ by stress condition (frequency: unpaired t-test, p = 0.4470; amplitude: unpaired t-test, p = 0.2754; n = 11-13 mice/group). (C-D) Peri-event time histograms of TS dopamine responses to rewarded (C) and unrewarded (D) nosepokes. Significant differences (p < 0.05 for more than 200 ms) between stressed (n = 11) and unstressed (n = 13) groups are denoted by purple bars. (E) Left, Schematic of photometry recording strategy for the DMS. Right, Overall amplitude and frequency of DMS dopamine transients did not differ by stress condition (frequency: p = 0.6393; amplitude: p = 0.3636; n = 14-16 mice/group). (F-G) Peri-event time histograms of DMS dopamine responses to rewarded (F) and unrewarded (G) nosepokes. No significant differences were detected between stressed (n = 16) and unstressed (n = 14) mice. All data presented as mean ± SEM. See also Table S2 for detailed statistics.

We did not observe general effects of stress on overall session-wide frequency and amplitude of TS dopamine transients (Figure 4B, left). However, stress strongly potentiated TS dopamine release at the time of a rewarded nosepoke (Figure 4C)^54^. Unrewarded nosepokes led to smaller transient dopamine responses that did not differ based on stress history (Figure 4D). TS dopamine responses to rewarded port entries were less prominent than responses to nosepokes (Figure S6B), demonstrating that dopamine in the TS is more closely linked to action than reward.

We did not observe effects of adolescent stress on the intrinsic excitability of TS-projecting dopamine neurons in the SNL (Figure S6C-F), indicating that changes in the properties of dopamine neurons themselves were not likely to be responsible for increased in vivo dopamine release. This similarity also indicated that adolescent stress-induced CeA→SNL hyperexcitability is not compensated for by downstream homeostatic adaptations, allowing excessive CeA neuron activity to propagate through the circuit.

Because our previous work indicated that DMS dopamine release controls the development of punishment-resistant reward-seeking in unstressed, standardly raised and housed mice^26^, we also performed DMS-targeted fiber photometry recordings with GRAB-DA2m in a separate group of mice during RI60 training to assess whether there were any similar increases in DMS dopamine that could explain our behavioral results. However, we found no changes in overall DMS dopamine activity (Figure 4E, Figure S6G) or in DMS dopamine responses to time-locked to rewarded or unrewarded nosepokes following adolescent stress (Figure 4F-G, Figure S6H). Therefore, we hypothesized that stress-induced elevations in TS dopamine were overriding DMS dopamine signals, inducing a fundamental switch in the striatal subregion driving the development of punishment-resistant reward-seeking.

### Elevated TS dopamine release on rewarded nose pokes represents a computational gain-of-function following adolescent stress

How might increases in TS dopamine signals promote punishment-resistant reward-seeking? Since previous work has proposed that TS dopamine encodes an action prediction error (APE)^55^, one possibility is that TS dopamine is a value-free signal that directly reinforces nosepoking. However, we observed that TS dopamine responses to rewarded nosepokes were larger than responses to unrewarded nosepokes (Figure 4C-D), suggesting that TS dopamine in this task carries an additional signal, rather than reflecting a pure value-free APE.

Given the established role of the TS in auditory processing^56^ and its proposed role in processing salience^57^, it is possible that the rewarded nosepoke response reflects added salience due to an audible solenoid click associated with the reward dispenser. If TS dopamine reflects a combination of APE and auditory salience, we should observe: 1) a negative relationship between lifetime nosepokes and dopamine release in all nosepokes (as predicted by APE, where signals should decrease with repeated performance), and 2) the negative relationship should be larger at rewarded nosepokes (to account for APE + auditory habituation). To examine this possibility, we used a functional linear mixed model (FLMM; Figure 5A-C)^58^ to query the relationship between lifetime nosepoke number and trial-by-trial variance in TS dopamine release. All predictors were z-scored before modeling to enable direct comparison of the influence of each variable (see Methods). Dopamine release followed an APE and habituation-like pattern in unstressed mice, with negative correlations between total nosepoke number and dopamine release around the peak of both rewarded and unrewarded nosepokes. In stressed mice, however, this effect was significantly attenuated for rewarded nosepokes, and nearly absent for unrewarded nosepokes (Figure 5D-F, Figure S7A-B). Thus, adolescent stress blunts encoding of APE and may prevent habituation to the salient reward delivery-related auditory cue during RI60 training.

**Figure 5.**
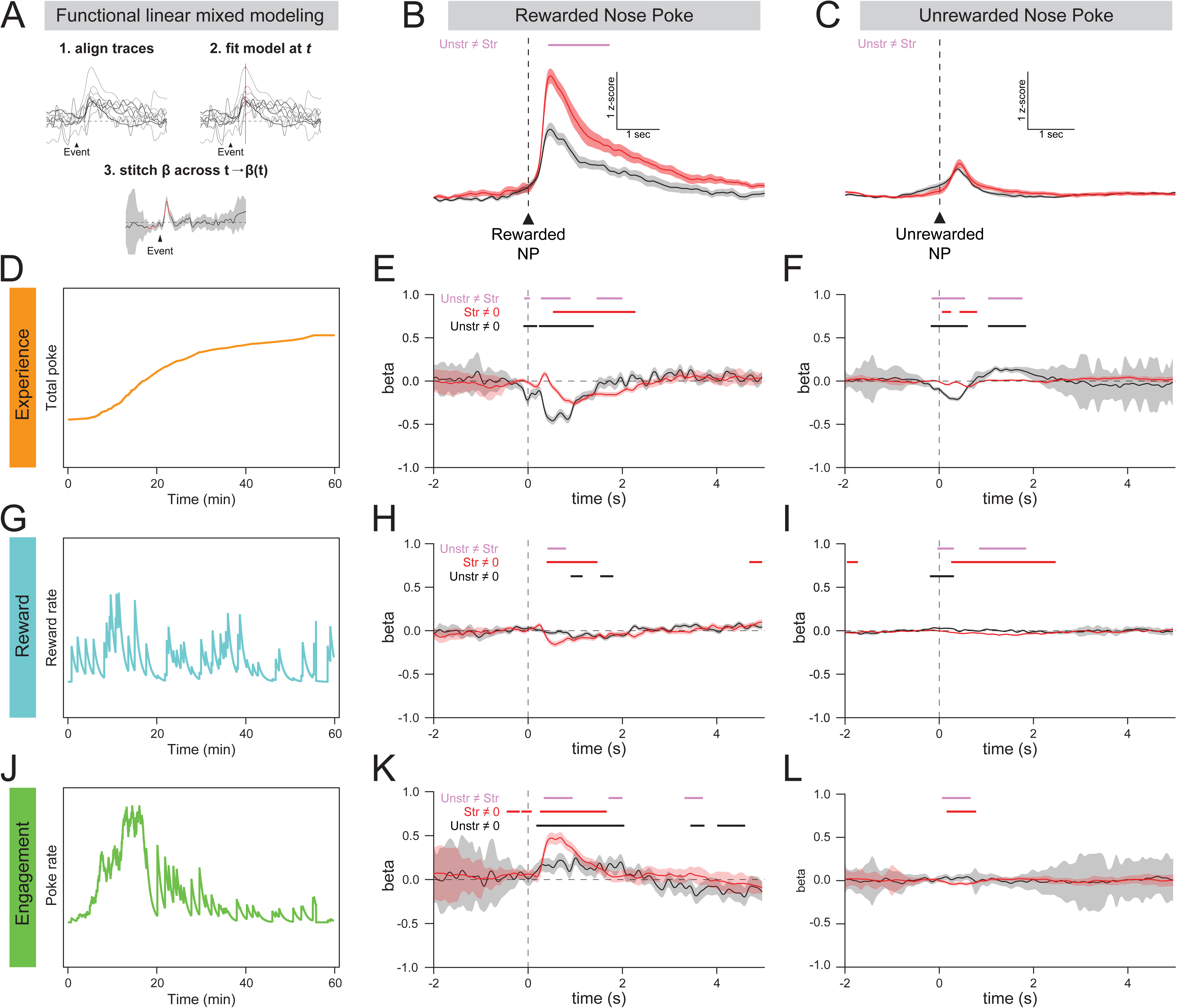
Functional linear mixed modeling reveals a computational switch in trial-by-trial dopamine encoding. (A) Schematic representing functional linear mixed modeling (FLMM). After aligning traces to events, a mixed model is fit at each time point within a window surrounding the event, and a trace showing the β coefficient (β(t)) is constructed across the event window. (B-C) TS dopamine responses to rewarded (B) and unrewarded (C) nosepokes (same as Fig. 4C-D, to allow easy comparison of dopamine traces with FLMM model outputs). (D) Total number of nosepokes across time for an example RI60 session. (E-F) FLMM beta values showing the relationship between the total number of nosepokes performed (“experience”) and TS dopamine responses to rewarded (E) and unrewarded (F) nosepokes. Black trace = unstressed β(t), red trace = stressed β(t). Black bars are where unstressed trace ≠ 0, red bars are where stressed trace ≠ 0, and purple bars are where unstressed trace ≠ stressed trace. (G) Recent reward rate across time for an example RI60 session. (H-I) FLMM beta values showing the relationship between the recent reward rate and TS dopamine responses to rewarded (H) and unrewarded (I) nosepokes. (J) Recent nosepoke rate across time for an example RI60 session. (K-L) FLMM beta values showing the relationship between the recent nosepoke rate and TS dopamine responses to rewarded (K) and unrewarded (L) nosepokes.

Another prediction of APE is that TS dopamine is a value-free signal, and thus should not correlate with reward- or engagement-related measures. To test this prediction, we ran FLMMs on TS dopamine and recent reward rate (calculated as a causal leaky integrator over rewarded pokes, τ = 60, see Methods; Figure 5G) and recent nosepoke rate (as a proxy for engagement, calculated same as reward rate, Figure 5J). In unstressed mice, dopamine responses to rewarded and unrewarded nosepokes were only weakly and briefly related to reward rate, consistent with a lack of value or state encoding. In stressed mice, however, TS dopamine release for rewarded nosepokes was higher on trials with low recent reward rate (Figure 5H, Figure S7C), consistent with value-based encoding that is not typically seen in the TS but is commonly associated with dopamine signals in the dorsomedial striatum (DMS) and nucleus accumbens (NAc)^55,59^. Dopamine release at unrewarded nosepokes was significantly more related to reward rate compared to unstressed controls, but at a lower magnitude and with less temporal precision compared to rewarded nosepokes (Figure 5I, Figure S7D).

TS dopamine signals for rewarded nosepokes also encoded engagement in the task to a significantly larger degree in stressed mice (Figure 5K, Figure S7E). If this encoding was purely motor, TS dopamine for unrewarded nosepokes would also track engagement; however, this was not the case (Figure 5L, Figure S7F), suggesting reward-specific processing. Taken together, these analyses show that adolescent stress leads to a computational switch in TS dopamine signaling, namely a loss of APE-like encoding and a gain of value and engagement encoding.

### Normalization of stress-induced phenotypes prevents adolescent stress-induced increase in punishment resistance

We next asked if we could prevent an increase in punishment-resistant reward-seeking by counteracting the stress-induced changes either in CeA→SNL neurons or in downstream TS dopamine release (Figure 6A). First, to dampen persistent hyperexcitability in CeA→SNL neurons, we overexpressed Kir2.1, an inwardly-rectifying potassium channel, in a projection specific manner by injecting a retrograde cre virus (CAV-cre) into SNL and an AAV carrying Cre-dependent Kir2.1-eYFP (AAV8-hSyn-FLEX-loxP-Kir2.1-2A-GFP) into CeA. Control mice were instead injected with an AAV carrying Cre-dependent eYFP (AAV5-EF1α-DIO-eYFP-WPRE-AGH). We compared both stressed and unstressed mice overexpressing Kir2.1 in CeA→SNL neurons with controls, resulting in four groups of mice (unstressed::eYFP, unstressed::Kir2.1, stressed::eYFP, stressed::Kir2.1; Figure 6B).

**Figure 6.**
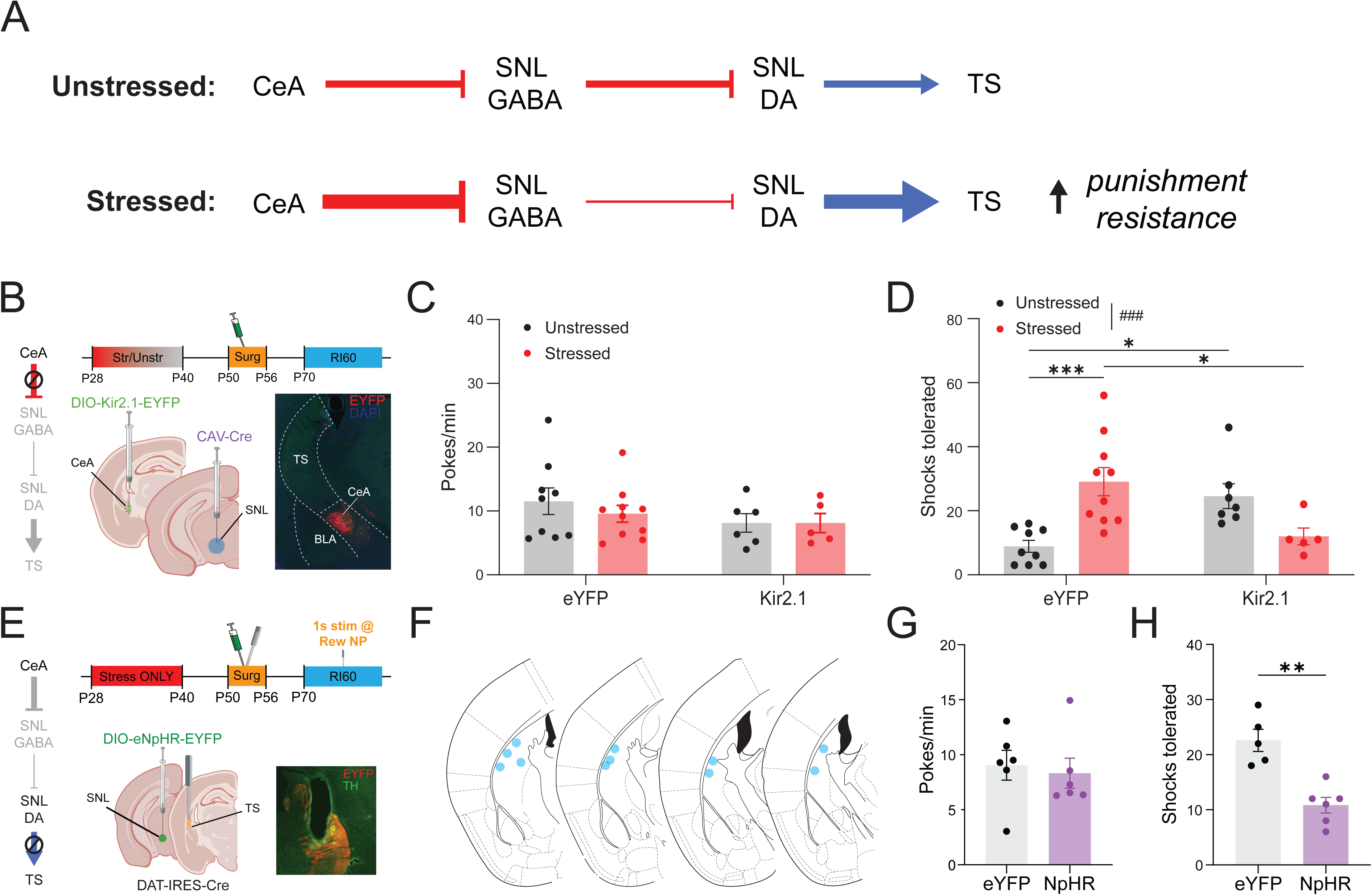
Normalizing CeA→SNL excitability or counteracting potentiated TS dopamine release prevents adolescent stress-induced increases in punishment-resistant reward seeking. (A) Schematic of hypothesized circuit changes induced by adolescent stress. In stressed mice, CeA hyperactivity increases inhibition onto SNL GABA neurons, leading to disinhibition of TS dopamine release, promoting punishment resistance. The hypothesis is tested via causal manipulations in this figure. (B) Schematic of Kir2.1 overexpression experiment to reduce CeA→SNL neuron hyperexcitability. Stressed and unstressed mice were injected with CAV-Cre in SNL and Cre-dependent Kir2.1 (or Cre-dependent EYFP as a control) in the CeA to specifically overexpress Kir2.1 in CeA→SNL neurons. Example histology (right), shows Kir2.1 overexpression limited to CeA. (C) RI60 nosepoke rates were not affected by stress or virus condition (2-way ANOVA, stress: p = 0.1869, virus: p = 0.5867, interaction: p = 0.5929, n = 5-10 mice/virus/group) (D) Number of shocks tolerated during the shock probe across stress and virus conditions. In mice expressing YFP virus, stressed mice had increased punishment-resistant reward seeking compared to unstressed controls. In mice expressing Kir2.1 virus, dampening CeA→SNL activity decreased punishment resistance in stressed mice (counteracting the effects of stress), but increased punishment resistance in unstressed controls (2-way ANOVA with Tukey’s post-hoc tests, stress: p = 0.3197, virus: p = 0.8522, interaction ^###^p = 0.0002; post-hoc (stressed-unstressed): stressed::eYFP vs unstressed::eYFP, ***p = 0.0010, stressed::eYFP vs. stressed::Kir2.1, *p = 0.0228, unstressed::eYFP vs. unstressed::Kir2.1, *p = 0.0233, all other comparisons p > 0.05, n = 5-10 mice/virus/group). (E) Schematic of NpHR experiment to inhibit TS dopamine release. Stressed DAT-Cre mice were injected with a Cre-dependent NpHR virus (or Cre-dependent EYFP as a control) in SNL, and dopamine terminals in TS were inhibited using 625nm light. Dopamine terminals in the TS were inhibited for 1s following rewarded nosepoke starting during RI30 and continuing through RI60. No inhibition was delivered during shock probe sessions. (F) Map of TS probe placements for NpHR experiment. (G) Inhibition of TS dopamine terminals did not change RI60 nosepoke rates in stressed mice (Mann-Whitney exact test, p = 0.3095, n = 6 mice/group). (H) Inhibition of TS dopamine terminals decreased punishment-resistant reward-seeking in stressed mice (Mann-Whitney exact test, **p = 0.0043, n = 5-6 mice/group). All data presented as mean ± SEM. ^∗^*p* < 0.05; ^∗∗^*p* < 0.01; ^∗∗∗^*p* < 0.001; ^∗∗∗∗^*p* < 0.0001 for main effects or pairwise comparisons; ^#^*p* < 0.05; ^##^*p* < 0.01; ^###^*p* < 0.001; ^####^*p* < 0.0001 for interactions.

Behaviorally, we did not observe significant effects of adolescent stress or Kir2.1 overexpression on nosepoke rate in this cohort (likely due to smaller cohort size; Figure 6C), but stressed::eYFP mice tolerated significantly more shocks than unstressed::eYFP mice, consistent with elevated punishment-resistant reward-seeking following adolescent stress (Figure 6D). Overexpression of Kir2.1 in the CeA→SNL projection, reduced the shocks tolerated by stressed mice (stressed::Kir2.1 vs stressed::eYFP; Figure 6D), demonstrating the necessity of hyperexcitability in the CeA→SNL projection for the observed adolescent stress-induced phenotype. Intriguingly, this same manipulation increased shocks tolerated in unstressed controls (unstressed:Kir2.1 vs unstressed::eYFP; Figure 6D). The differential effect of CeA→SNL inhibition across stress conditions implies that the role of CeA→SNL activity in regulating behavior is state-dependent, and fundamentally changed by adolescent stress, consistent with our observations of a computational shift in TS dopamine encoding downstream of stress-induced changes in the CeA. Overall, these results confirm that CeA→SNL hyperexcitability is necessary for the adolescent stress-induced increase in punishment-resistant reward-seeking.

If CeA→SNL hyperexcitability increases punishment-resistant reward-seeking by elevating TS dopamine on rewarded nosepokes, we reasoned that inhibition of this dopamine signal in a temporally-restricted manner should also be sufficient to reduce the development of punishment-resistant reward-seeking. To test this hypothesis, we inhibited TS dopamine release in stressed mice during rewarded nosepokes using the inhibitory opsin eNpHR3.0. We expressed eNpHR3.0 in dopamine neurons by injecting an AAV carrying cre-dependent eNpHR3.0 into the SNL of DAT-IRES-cre mice and implanted a fiber optic probe above TS for terminal inhibition (Figure 6E-F). We applied orange light (625 nm, 1s, 10 mW at fiber tip) on rewarded nosepokes during RI30 and RI60 training. There was no difference in RI60 nosepoke rate between NpHR and YFP control mice (Figure 6G), but TS dopamine inhibition during training reduced punishment-resistant reward-seeking when stressed mice were tested in the shock probe without optogenetic inhibition (Figure 6H). Thus, increased TS dopamine release is necessary for the development of the increased punishment resistance following adolescent stress. Taken together, these causal experiments confirm that the CeA→SNL→TS pathway is necessary for adolescent stress-induced changes in punishment-resistant reward-seeking, isolating this pathway as one of great interest for investigating the relationship between adolescent stress and addiction risk.

## Discussion

Stress is a crucial risk factor in the development of psychiatric disorders, including addiction, at all stages of life. However, the specific consequences of stress may depend not only on severity and exposure length, but also on developmental stage. Here, we find that chronic unpredictable stress in early adolescence produces a strong, persistent increase in the propensity of male and female mice to develop punishment-resistant reward-seeking in adulthood. The effects of stress are specific to adolescence, since the same stress paradigm did not change punishment-resistant reward-seeking when applied in adulthood. Our findings in mice align with human data showing that adolescent stress is associated with a long-lasting predisposition towards addictive disorders^14,15,60,61^. Therefore, this mouse model offers an opportunity to probe the mechanisms by which neural circuit perturbations create addiction vulnerability. The changes we uncover here (in the CeA→SNL→TS pathway) may be specific to the experience of adolescent stress, emphasizing a need to stratify patients based on stress history to search for relevant neural changes in humans and develop personalized approaches to addiction treatment.

The need for stratification is especially indicated by our finding that the increase in punishment-resistant reward-seeking following adolescent stress is achieved by shifting the dopaminergic “balance of power” within the striatum from DMS to TS. If the experience of adolescent stress fundamentally reshapes the computational function of TS dopamine, and the subregional dependence of behavior, treatments likely must be specialized. The treatment must be tailored towards correcting the particular “gain-of-function” within the circuit following adolescent stress; otherwise, undesired, even opposite, outcomes may occur. Encouragingly, however, we observe that counteracting the gain-of-function in adulthood is sufficient to diminish elevated punishment-resistant reward-seeking even when the intervention is applied after the window of adolescent stress, and after the onset of CeA→SNL hyperexcitability.

A further understanding of the mechanisms causing CeA→SNL hyperexcitability will be important for translation. Here, we show using biophysical modeling that channels producing the M-type potassium current are promising candidates for future investigation. Changes in M-currents could explain much of the hyperexcitability across diverse CeA electrophysiological subtypes. M-currents are carried by Kv7 channels (primarily 7.2 and 7.3 in the central nervous system)^62^, and are slow, non-inactivating potassium currents that act as a brake on excitability. Chronic and acute stressors have been shown to decrease I_m_ in various brain regions, such as the prefrontal cortex^63^, paraventricular nucleus^64^, BNST^65^, and the ventral tegmental area^66^, and in CRF+ CeA neurons^47,48^. Therefore, our modeling results present a new set of testable hypotheses regarding the ion channels mediating adolescent stress-induced hyperexcitability, and if borne out, may lead to the development of therapeutic interventions.

Our results align with previous studies suggesting CeA activity has a net disinhibitory effect on downstream dopamine signaling^41,52^. However, CeA→SNL neurons innervate both GABAergic and dopaminergic neurons in the SNL^41^, and it remains unclear if the same CeA neurons synapse onto SNL GABA and SNL dopamine neurons. Future work should investigate differences in information flow in CeA→SNL GABA and CeA→SNL dopamine projections and explore whether experience dynamically re-balances inhibition and disinhibition within this pathway.

In this work, we highlight a role for TS dopamine in punishment-resistant reward-seeking in mice with a history of adolescent stress, which contrasts with previous work implicating DMS dopamine in unstressed mice^26^. The lack of adolescent stress effects on DMS dopamine signaling suggests that control over this behavior may be the result of an interplay between or competition across striatal subregions, as opposed to fixed, cross-context functions for each domain^67^.

A corollary that emerges from this hypothesis is that there may be multiple cognitive routes to punishment-resistant reward-seeking. Following adolescent stress, potentiated TS dopamine release may assign increased salience to the reward-related auditory cue^57^, overriding goal-directed DMS control. This idea is supported by the major stress-induced changes in TS dopamine encoding of task variables (Figure 5). Combined with previous findings, our results allow us to speculate about the TS-dependent cognitive route to punishment-resistant reward-seeking. In one previous study, optogenetic activation of CeA→SNL neurons was found to support intracranial self-stimulation but not conditioned place preference^41^. This result is consistent with a signal that reinforces instrumental associations but is not inherently rewarding. Our finding of the acquisition of a value signal in TS dopamine means that value signals have a new route to directly reinforce instrumental associations and promote compulsive engagement. Our behavioral analysis indicating increased continuous task engagement during RI60 training following adolescent stress is also consistent with this framing, although it will require further study to validate.

Our findings are also important to compare with other recent results on TS dopamine release in which this signal has been linked to threat and aversive processing^31,50^. Indeed, a recent paper reports no TS dopamine response to reward at all^31^—seemingly contradicting our finding of reward-related activity. However, these other studies focus on the caudal-most part of the TS (AP ∼ −1.8mm), whereas this work and others observe salience-related release at slightly more anterior coordinates (AP ∼ - 1.3mm). Thus, the in vivo functions of TS domains along the anterior-posterior axis are another area that warrants further study.

In conclusion, we show here that developmental experience can impact the computational function of the dopamine system by altering the activity of key inputs to the midbrain. In particular, the stress-sensitive CeA may be an upstream switch capable of tuning the contribution of the TS to behavior. In individuals with a history of adolescent stress, therapies that aim to rebalance control within dopaminergic subcircuits may mitigate risk for addiction-relevant behavioral phenotypes.

## Resource availability

### Lead contact

Further information and requests for resources and reagents should be directed to and will be fulfilled by the lead contact, Talia Lerner.

### Materials availability

This study did not generate new unique reagents.

### Data and code availability

- Data have been deposited at Zenodo and are publicly available as of the date of publication at 10.5281/zenodo.21500422
- All original code has been deposited on Github and is publicly available at https://github.com/jnadelneuro/aCUS-paper/releases/tag/preprint as of the date of publication.
- Any additional information required to reanalyze the data reported in this paper is available from the lead contact upon request

## Acknowledgments

We thank the Lerner laboratory for helpful discussions and critical feedback throughout the project. We thank the Center for Comparative Medicine at Northwestern University for providing animal care and husbandry. This work was supported by the National Institutes of Health (DP2MH122401, R01MH125885, R01DA063125 to T.N.L., T32MH067564 and F31DA060560 to J.A.N.).

## Author Contributions

J.A.N. and T.N.L. conceived the experiments, and J.A.N, E.S.S., M.Z., B.D., and M.H.K. executed them. J.A.N. analyzed the data, including modeling studies. S.P. provided oversight for anxiety-related experiments. T.N.L. provided oversight and support for all aspects of the project. J.A.N. and T.N.L. wrote the manuscript with input from the other authors.

## Declaration of Interests

The authors declare no competing interests.

## Key resources table

| REAGENT or RESOURCE | SOURCE | IDENTIFIER |
| --- | --- | --- |
| **Antibodies** |  |  |
| Chicken anti-Tyrosine Hydroxylase | Aves Labs | Cat#TYH; RRID:AB_10013440 |
| Goat anti-Chicken IgY (H+L) Secondary Antibody, Alexa Fluor™ 488 | Invitrogen | Cat#:A-11039; RRID:AB_253409 |
| Rabbit anti-GFP | Invitrogen | Cat# A11122; RRID:AB_221569 |
| Goat anti-Rabbit IgG (H+L) Cross-Adsorbed Secondary Antibody, Alexa Fluor™ 594 - A-11012 | Invitrogen | Cat#:A-11012; RRID:AB_2534079 |
| **Bacterial and virus strains** |  |  |
| AAV9-hSyn-GRAB_DA2m | Addgene | Cat# 140553-AAV9; titer $2 \times 10^{12}$ vg/mL |
| AAV8-hSyn-FLEX-loxP-Kir2.1-2A-GFP | WZ Biosciences (from Addgene plasmid #161574) | Cat:#161574, titer: $8 \times 10^{12}$ vg/mL |
| AAV5-EF1α-DIO-eNpHR3.0-EYFP | Addgene | Cat#:26966-AAV5; Lot#:v32533; titer: $2.4 \times 10^{13}$ vg/mL |
| AAV5-EF1α-DIO-eYFP-WPRE-AGH | UNC Vector Core | Cat# 27056; titer $3.4 \times 10^{12}$ vg/mL |
| CAV2-Cre | IGMM (Montpellier) | titer: $3.7-3.9 \times 10^{12}$ vg/mL |
| **Chemicals, peptides, and recombinant proteins** |  |  |
| Red retrobeads IX | LumaFluor Inc | Cat#:78R180 |
| D-AP5 | Cayman Chemical | Cat#:79055-68-8 |
| NBQX disodium salt | Tocris Bioscience | Cat#: 479347-86-9 |
| Gabazine (SR-95531; "GBZ") | Tocris Bioscience | Cat#:104104-50-9 |
| Isoflurane | Henry Schein | Cat#:26675-46-7 |
| Meloxicam | Covetrus |  |
| Bupivacaine | Hospira |  |
| Euthasol | Virbac |  |
| Puralube ophthalmic ointment | Dechra |  |
| Metabond | Parkell |  |
| Flow-it ALC dental epoxy | Pentron |  |
| Vetbond tissue adhesive | 3M |  |
| Precision Reward Pellets®, Sucrose | Bio-Serv | Cat#: F07595 |
| Loctite 454 | Henkel/Loctite |  |
| Normal goat serum | Jackson ImmunoResearch Laboratories | Lot#153636; RRID:AB_2336990 |
| DAPI Fluoromount-G | SouthernBiotech | Cat#:0100-01 |
| **Critical commercial assays** |  |  |
| Corticosterone ELISA kit | Enzo Life Sciences | Cat#:ADI-900-097 |
| Deposited data |  |  |
| CeA single-cell RNA-seq (Wang et al., 2023; reanalyzed) | GEO | GSE213828 |
| **Experimental models: Organisms/strains** |  |  |
| Mouse: WT: C57BL/6J | Jackson Laboratories | Strain #000664; RRID:IMSR_JAX:000664 |
| Mouse: DAT-IRES-Cre: B6.SJL-*Slc6a3tm1.1(cre)Bkmn/J* | Jackson Laboratories | Strain #006302; RRID:IMSR_JAX:006302 |
| **Software and algorithms** |  |  |
| Wavesurfer v0.945 | HHMI Janelia | RRID:SCR_021529; https://wavesurfer.janelia.org |
| GuPPy | Lerner Lab | https://github.com/LernerLab/GuPPy/wiki; RRID: SCR_022353 |
| Synapse | Tucker Davis Technologies | https://www.tdt.com/component/synapsesoftware/; RRID: [SCR_024878](https://www.tdt.com/component/synapsesoftware/) |
| MED-PC V | Med Associates | https://www.med-associates.com/medpc-v/; RRID: [SCR_012156](https://www.med-associates.com/medpc-v/) |
| GraphPad Prism 10 | GraphPad | RRID:SCR_002798 |
| Python 3 | Python | RRID:SCR_008394 |
| RStudio 4.3.5 | R | RRID:SCR_000432 |
| FastFMM (R) | Loewinger et al., 2024 | https://cran.r-project.org/web/packages/fastFMM/index.html |

## Methods

### Animals and Breeding

Male and female C57BL/6J and (DAT)::IRES-Cre knockin mice (JAX006660) were obtained from The Jackson Laboratory and bred in house. For transgenic lines, only heterozygous mice were used for experiments. A timed breeding paradigm was used to generate large cohorts that could be stressed simultaneously. Breeding occurred on a conventional 12h:12h light cycle (light 9:00 am to 9:00 pm). Male mice were isolated at least 10 days prior to pairing and females were placed in cages of four to synchronize estrous cycles. 4 days prior to pairing, male bedding was placed into each female cage. One male and one female were then paired for three days. After three days, males were removed and females were paired to prevent isolation stress from affecting *in utero* development and maternal behavior. Pups were weaned at P21 and immediately moved to a reverse light cycle room (12h:12h, light 9:00 pm to 9:00 am), where they remained for the duration of experiments. All behavioral tests occurred during the dark cycle in red light rooms. Mice were group housed and had *ad libitum* access to food and water unless undergoing operant behavior. All experiments were approved by the Northwestern University Institutional Animal Care and Use Committee.

### Stress Paradigm

Adolescent stress was applied from P28-40 using a condensed chronic unpredictable stress paradigm adapted from Yohn and Blendy^33^. Behavioral testing began 30 days later, at P70. Details of the stressors applied are listed in Supplementary Table 1. Briefly, mice were exposed to stressors three times a day: in the morning, afternoon, and overnight, for 12 consecutive days. Adult stress was applied from P70-82 using the same schedule of stressors, and behavioral testing began 30 days later, at P112 (Figure 1A). Unstressed controls were only handled for ear-tagging; otherwise, they remained undisturbed until surgeries or behavioral testing began.

### Blood Collection and Corticosterone Enzyme Linked Immunosorbent Assays (ELISAs)

Mice were gently restrained in a Tailveiner® Restrainer for Mice (Braintree Scientific), and 30 μL of blood was sampled from the lateral tail vein via a perpendicular slit with a razor. Blood draws occurred between ZT15-16 (zeitgeber times, 15-16 hours after lights on). Plasma was separated by centrifugation at 10,000 RPM and 4°C for 10 minutes. Plasma samples were stored at −80°C prior to ELISAs. Plasma corticosterone was quantified using a Corticosterone ELISA kit from Enzo Life Sciences, according to the manufacturer’s instructions.

### Blood Glucose Measurements

At ZT16, mice were gently restrained in a Tailveiner® Restrainer for Mice (Braintree Scientific) and a drop of blood was taken from the lateral tail vein directly into a test strip (Medilax). The test strip was immediately inserted into a glucometer (Medilax) and values were recorded.

### Open Field Test

Mice were placed into the center of a 28×28×28 cm arena in a dimly lit room and recorded for 10 minutes. The arena was split into periphery and center for analysis. The center was illuminated to 400 LUX while the corners ranged between 30-40 LUX. Cumulative distance traveled, time spent in each zone, and entries into each zone were calculated by ANY-Maze tracking software based off the mouse’s center body point (Stoetling, Wood Dale, IL).

### Elevated Plus Maze Test

EPM was performed as described in Kondev et al 2022^68^. Briefly, the elevated plus maze consisted of two sets of 68 cm arms (one set open, one set closed) intersecting in a central platform. Closed arm walls were 15 cm high, and the maze was elevated 40 cm off the floor. Mice began by being placed in one of the closed arms, and each mouse’s position was tracked for 10 minutes using ANY-maze software.

### Stereotaxic Surgery

Viral injection, retrobead injection, and optic fiber implant surgeries were performed on young adult mice at P50-56. For P40 electrophysiology experiments, retrobead injections were performed at P40 and experiments began at P41. Mice were anesthetized in an isoflurane chamber at 4-5% isoflurane (Henry Schein) and then placed on a stereotaxic frame (Stoelting). Anesthesia was maintained at 1-2% for the duration of the surgery. Mice were injected with meloxicam (Covetrus, 20 mg/kg) and 0.9% sterile saline subcutaneously prior to surgery to minimize post-surgical pain and dehydration. Ophthalmic ointment (Puralube, Dechra) was applied to the eyes to prevent ocular dehydration. A far infrared heating pad (Kent Scientific) was used to maintain body temperature at ∼37°C. Hair was removed from the top of the head using Nair, and exposed skin was disinfected with alcohol and a povidone-iodine solution. Prior to scalp incision, bupivacaine (Hospira, 2 mg/kg) was injected subcutaneously at the incision site. The scalp was opened using a sterile scalpel and the skull was leveled before drilling holes at the appropriate stereotaxic coordinates (CeA: AP −1.2, ML ±2.9-3.0, DV −4.6; SNL: AP −3.0, ML ±1.5, DV −4.5; TS: AP −1.3, ML ±3.4, DV −3.5-3.6; DMS: AP +0.8, ML ±1.5 DV −2.8; BNST: AP +0.5, ML ± 0.9, DV −4.5; PAG: AP −4.3, ML ±0.4, DV −3.2) for the given experiment. Bregma was calculated as the crossing pointing between the sagittal suture with the best-fitting parabola on the coronal suture^69^. To drill skull holes, a micromotor drill (Stoelting, 51449) was moved to the appropriate coordinates with the aid of a digital stereotaxic display. Viruses and retrobeads were infused at 100 nL/min through a blunt 33-gauge injection needle using a syringe pump (World Precision Instruments). The needle was left in place for 10 minutes following the end of injection, then slowly retracted to avoid leakage up the injection tract. For all surgeries that did not involve fiber implants, the scalp incision was closed with non-absorbable sutures (Ethicon, 661H) and tissue adhesive (Vetbond, 3M).

For surgeries involving implants, implants were secured to the skull with Metabond (Parkell) and Flow-it ALC blue light-curing dental epoxy (Pentron). Mice were allowed to recover until ambulatory on a heated pad, then returned to their homecage with moist chow and DietGel available. The mice were checked after 24h and provided with another dose of meloxicam. Mice then recovered for ∼3 weeks before experiments began.

For retrobead electrophysiology experiments, red retrobeads (LumaFluor Inc) were diluted 1:4 (dilution factor) in sterile saline and 250 nL were injected into the appropriate brain region (SNL, BNST, PAG, or TS) in WT mice.

For fiber photometry experiments, we infused 500 nL of AAV9-hsyn-GRAB_DA2m (2e12 vg/mL, Addgene 140553-AAV9) unilaterally into the DMS and/or TS of WT mice. Fiber optic implants (Doric Lenses; 400 μm, 0.48 NA) were placed 0.1 mm above DMS and TS DV coordinates.

For Kir experiments, we infused 500 nL of AAV8-hSyn-FLEX-loxP-Kir2.1-2A-GFP (8e12 vg/mL, Addgene Plasmid #161574, fabricated by WZ Biosciences) or AAV5-EF1a-DIO-eYFP-WPRE-AGH (3.4e12, UNC #27056) into the CeA and 250 nL of CAV-cre (3.7-3.9e12 vg/mL, IGMM) into the SNL in WT mice.

For NpHR experiments, we infused 1000 nL of AAV5-EF1a-DIO-eNpHR3.0-EYFP (2.4e13, Addgene #26966) or AAV5-EF1a-DIO-eYFP-WPRE-AGH (3.4e12, UNC #27056) into the SNL and implanted bilateral optogenetic ferrules (Prizmatix; 500μm core, 0.66 NA) over the TS of DAT-cre mice.

### RI60 Operant Behavior

Mice were food restricted to 85% of ad libitum body weight for the duration of operant training. Mice were given one day of habituation to operant chambers (Med Associates) and (if undergoing photometry or optogenetics experiments) tethering with patch cords (Doric Lenses) for 30 minutes. They were then trained to retrieve sucrose pellets (20mg, BioServ) from a magazine port. For magazine training, pellets were delivered to the port on a random interval (RI60) schedule non-contingently for 30 minutes. Next, operant training began with all training sessions lasting 1 hour or until 50 rewards were obtained. Mice were first trained to associate nose poking a preassigned nose poke (right or left) with reward on a fixed ratio (FR1) schedule where each poke on the selected side delivered a reward, with the caveat that mice had to retrieve the reward (as measured by making a port entry) before they could earn the next. After a mouse completed 3 consecutive days with an average of 30 rewards earned (and a minimum of 4 days on FR1), they were moved to a random interval schedule of reinforcement.

Mice were initially trained for 4 days on an RI30 schedule, then moved to RI60. Mice underwent 14 days of RI60 before testing on the shock probe. For the RI30 schedule, a random number between 1 and 30 was drawn every second. If a 5 was drawn, the drawing stopped, and reward was available at the next nose poke. After the rewarded nose poke, random drawing began again. The RI60 schedule was identical except numbers were drawn between 1 and 60.

### Lognormal Mixture Modeling of Behavior Data

Inter-poke interval times (IPIs) were calculated as the difference between consecutive nose pokes within each RI60 session, retaining intervals greater than 10 ms. Since IPIs are always positive and heavily tailed, their distribution was modeled as a mixture of lognormal components, with each component corresponding to a distinct latency mode. Sessions with fewer than 100 IPIs were excluded, as fit degraded sharply below this threshold. We tested models with 2-8 components, ultimately choosing 5 as model improvements plateaued after this number. We fit a hierarchical mixture on the data from all sessions. Component location (μ_k) and scale (σ_k) were shared across all sessions and groups, while the weights (π_k, constrained to be positive and sum to 1) were allowed to vary and were estimated separately for each session. This method structurally isolates reallocation of responding across components, but is insensitive to any global scaling of latencies. Parameters were estimated by expectation-maximization. To ensure robustness, the model was fit from 10 random restarts, each run for up to 80 EM iterations. The highest likelihood solution was then further iterated upon (up to 400 iterations, relative log-likelihood tolerance 1×10^−7^) to convergence. Since the mixing weights are compositional (positive and constrained to sum to 1) and therefore not independent, group differences in composition were tested with a Bayesian Dirichlet regression modeling the full five-component composition jointly (see Quantification and Statistical Analysis).

### Shock Probe

Mice were subjected to a foot shock probe after RI60 training to evaluate their level of punishment-resistant reward seeking. These probes were performed under an FR1 schedule of reinforcement where a mild (1s, 0.2 mA) footshock was paired with a subset of rewarded nose pokes on a RR3 schedule, so that every 3^rd^ nose poke on average was accompanied by a shock. During shock probes, the session ended after 60 minutes or 10 minutes of inactivity (no nose pokes on either side). There was no maximum number of rewards. Pokes within 5 seconds of a shock were defined as “compulsive returns”.

### Active Avoidance Behavior

A separate subset of mice underwent the two-chamber active avoidance task described in Lopez et al^70^. Briefly, mice were placed in custom 2-chamber shuttle boxes, with chambers separated by a plastic door (Med Associates). Mice were placed in either the left or right shuttlebox chamber, chosen randomly and alternating side each day. Upon task start, sound (2900 Hz) and white light (40 lux) cues began, and the door between chambers opened. If the mouse shuttled from the initial chamber to the opposite chamber within 5 seconds, both cues terminated and the ITI (avg. 45 seconds) began (“avoid” trial). If the mouse failed to shuttle within 5 seconds, a 0.4 mA shock turned on and continued for 25 seconds or until the mouse crossed into the other chamber (“escape” trial). Mice were tested on this paradigm for 7 days, 30 trials per day.

### Electrophysiology

We followed the method from Ting and colleagues^71^ to prepare acute brain slices from adult mice. Following Euthasol injection, mice were transcardially perfused with ice-cold N-Methyl-D-Glucamine (NMDG) artificial cerebrospinal fluid (ACSF) containing (in mM): 92 NMDG, 2.5 KCl, 1.2 NaH2PO4, 30 NaHCO3, 20 HEPES, 25 Glucose, 5 Na-Ascorbate, 2 Thiourea, 3 Na-Pyruvate, 10 MgSO4, 0.5 CaCl2 (Millipore Sigma). All extracellular solutions used for electrophysiology were saturated with 95%O2/5%CO2 and their pH and osmolarity were adjusted to 7.3-7.4 and 300±5 mOsm, respectively. After perfusion, the brain was quickly removed and the cerebellum was cut off coronally. The cut face of the brain was glued (Loctite 454) to a specimen holder and immersed into ice-cold NMDG ACSF. Coronal slices (250-300 μm thick) were made using a vibratome (Leica, VT1200S) set to 0.10 mm/s speed and 1.10 mm amplitude. Slices containing bead injections were saved to confirm injection sites, while CeA or SNL containing slices were used for recordings. Slices were allowed to recover for 45 min in three 15 min baths: (1) warm (33°C) NMDG ACSF; (2) warm (33°C) recovery ACSF, containing (in mM): 92 NaCl, 2.5 KCl, 1.2 NaH2PO4, 30 NaHCO3, 20 HEPES, 25 Glucose, 5 Na-Ascorbate, 2 Thiourea, 3 Na-Pyruvate, 1 MgSO4, 2 CaCl2; and (3) room temperature (RT) recovery ACSF. Finally, slices were kept at RT in recording ACSF, containing (in mM): 125 NaCl, 26 NaHCO3, 1.25 NaH2PO4, 2.5 KCl, 1 MgCl2, 2 CaCl2, 11 Glucose. During recordings, fresh ACSF was continuously delivered to the slice chamber at ∼1.5 ml/min and warmed to 30-32°C with an inline heater (Warner Instruments).

For all intrinsic excitability experiments, the following drugs were added to the recording ACSF: ACSF: D-AP5 (50 μM, Cayman Chemical), NBQX disodium (5 μM, Tocris Bioscience), and GBZ (10 μM, Tocris Bioscience). For all intrinsic excitability experiments, we used a modified potassium gluconate internal solution (in mM, 130 K-gluconate, 15 KCl, 10 HEPES, 2 Mg-ATP, 1 EGTA, 0.3 Na-GTP). Patch pipettes (3-7 MΩ) were pulled (Narishige, PC-100) from borosilicate glass (Warner Instruments, G150TF-4) and moved with the assistance of a micromanipulator (Sensapex). Cells were visualized with a 40x water-immersion objective (NA 0.8, Olympus, #N2667700) on a microscope (Olympus, BX51WI) equipped with infrared-differential interference imaging (DIC) and a camera (QImaging, Retiga Electro Monochrome). An LED light source (CoolLED, pE-300white) was used to illuminate the slice through the objective for targeted patching of bead-positive cells. Signals were recorded at 10 kHz using Wavesurfer v0.945 (https://wavesurfer.janelia.org/), a National Instruments Digitizer (NIDAQ X series PCIe-6323) and BNC Breakout (BNC-2090A), and a Multiclamp 700B amplifier (Molecular Devices). Intrinsic properties were assessed with a battery of protocols: 1) a frequency current set of depolarizing steps from 0 to 200 pA in 20 pA increments (500 ms steps) to obtain F-I curves; 2) a finer grain rheobase protocol of 0 to 100 pA in 5 pA steps (50 ms steps, if cells did not fire at 100 pA the protocol was re-run until a current evoked firing); 3) a sag protocol of −50, −100, and −150 pA steps; 4) a 30 second recording of spontaneous activity at rest for resting membrane potential. Input resistance, access resistance, and capacitance were measured from a −5 mV voltage clamp test pulse. All protocols were run three times and results were averaged for analysis.

Action potential and spike train features were extracted with the ipfx Python package from the Allen Institute. Spikes were detected as peaks crossing 0 mV with a 5 ms inter-event interval, and firing rate was calculated as spike count divided by step duration. Latency was calculated as the time from step onset to first spike. Adaptation index was defined as the normalized mean of consecutive ISI differences within a train. Resting membrane potential was calculated as the mean voltage over the 30 second spontaneous recording sweep. Rheobase was defined as the smallest current step evoking a spike. Sag ratio was defined as |V_peak − V_ss| / |V_peak − V_baseline|, where V_peak is the most negative voltage during the step, V_ss the mean voltage over its final 10%, and V_baseline the mean pre-step voltage. Data analysis was performed offline using custom-written Python scripts.

### Clustering Analysis of Electrophysiology Data

Electrophysiological features (firing rate across current injection, resting membrane potential, adaptation index, and latency to fire) were extracted for each CeA→SNL cell used in intrinsic excitability experiments. All numeric features were z-score normalized to ensure equal weighting of features with different units. Given the observed effect of stress on intrinsic properties, we statistically removed variance associated with the stress manipulation by fitting linear mixed-effects models for each feature (feature ∼ stress + (1 + stress | mouse)). The inclusion of the random intercept and slope per mouse accounts for inter-animal variability. Model residuals were extracted and used for subsequent analysis. We confirmed the effectiveness of this strategy by comparing the proportions of stressed and unstressed cells in each group, finding they were nearly identical.

Principal component analysis (PCA) was performed on the residuals, with the first 4 components chosen to retain for clustering after inspection of elbow plots. K-means clustering was applied to the PCA scores to identify distinct electrophysiological phenotypes in 3 groups. Clustering quality was assessed using the silhouette coefficient. To interpret the identified clusters, mean values of electrophysiological features were calculated for each cluster and split by stress and unstressed groups.

### Biophysical Modeling

All biophysical modeling was performed using NEURON and its associated python packages. Model morphology consisted of a somatic (15 x 15 μm) and dendritic (5 x 300 μm, 150 Ω x cm) component. Channel kinetics and initial conductance values were based on Li et al^45^, who modeled the same three CeA electrophysiological phenotypes (regular spiking, low threshold bursting, and late firing). All three cell types included 1) fast Na^+^, 2) delayed K^+^, 3) M-current (Kv7), 4) L-type Ca^2+^, 5) HCN, 6) Ca^2+^-dependent slow afterhyperpolarization, and 7) passive leak conductances. In addition to those base channels, LTB neurons (cluster 2) additionally had T-type Ca^2+^ channels and a slow sodium inactivation gate based on Qian et al^72^; late firing neurons (cluster 3) had A-type potassium channels (Kv4). Each model had 8-10 free parameters (maximal conductances for each channel along with leak reversal potential).

Electrophysiological data from each cluster from unstressed mice (Figure 3) were used to fit parameters using a Covariance Matrix Adaptation Evolution Strategy (CMA-ES), a derivative-free optimization algorithm. The cost function was a weighted sum of squared errors across: F-I spike counts at 0-200 pA (in 20 pA increments), passive membrane properties (RMP, input resistance, sag ratio), first-spike shape (threshold, peak, trough, half-width), and spike train dynamics (first-spike latency, mean inter-spike interval, ISI CV, and adaptation index). Each feature was normalized by its experimentally observed standard deviation to ensure proportional contribution to the cost function. Optimizations were repeated with 10 independent random seeds (population size 1200, 200 generations) to ensure robustness and avoidance of local minima.

To identify candidate conductance changes underlying the aCUS-induced phenotype in each cluster, we re-ran the optimization against electrophysiology data from aCUS mice using three complementary approaches. First, all parameters were frozen at their unstressed-optimized values to establish an upper bound on model error. Second, all parameters were freed, establishing a lower bound. Third, we systematically varied single conductance values while freezing all others at unstressed values. The gap between a candidate’s fitness and the upper and lower bounds was used to quantify what fraction of the stress-induced change that the candidate explains.

### Fiber photometry

All fiber photometry experiments were performed using fiber photometry rigs with optical components from Doric Lenses and Tucker Davis Technologies (TDT) controlled by a real-time processor from TDT (RZ10X), sampled at ∼1017 Hz. TDT Synapse software was used for data acquisition. 405nm and 465nm LEDs were modulated at 330 Hz and 210 Hz, respectively. LED currents were adjusted to return a voltage between 150-200mV for each signal, were offset by 5 mA, and were demodulated using a 4 Hz lowpass frequency filter. Behavioral timestamps were transmitted from operant boxes as TTL signals for alignment with the neural data. GuPPy, an open-source Python-based photometry analysis pipeline, was used to process fiber photometry data and align signals to TTLs (see Quantification and Statistical Analysis)^73^. Mice were recorded from at least 3 times throughout RI60 training, during early, middle, and late training.

### Inhibitory Optogenetic Stimulation

Mice injected with DIO-eNpHR or EYFP controls (see Stereotaxic Surgery, above) underwent RI60 training as described above. Starting at RI30 and lasting through the remainder of RI60 training, rewarded nose pokes were paired with a continuous pulse of orange/red light (625 nm, 1s, 10 mW) generated by a high-power LED light source and pulse generator (Prizmatix). For the shock probe, mice were untethered and did not receive light stimulation.

### Perfusions and Histology

Mice received lethal i.p. injections of Euthasol (Virbac, 1mg/kg) to induce a rapid onset of unconsciousness and death. Once unresponsive (as determined by lack of responsiveness to a firm toe pinch), an incision was made up the middle of the body cavity. A needle was inserted into the left ventricle of the heart, the right atrium was punctured, and PBS followed by 4% PFA was infused into the left ventricle. The mouse was then decapitated, and its brain was removed and fixed overnight in 4% PFA at 4°C. The next day, brains were transferred to a 30% sucrose solution for at least 48 hours. Tissue was then sectioned on a freezing microtome (Leica) at 25-50 um, then stored in cryoprotectant (30% sucrose, 30% ethylene glycol, 1% polyvinyl pyrrolidone in PB) at - 20°C until immunostaining.

Selected slices were transferred to a well in a 12-well plate and washed 4x for 5 minutes in PBS to remove cryoprotectant. Slices were then blocked in 3% normal goat serum in 0.3% PBS-T for 1-2h at room temperature. The slices were then transferred to solutions containing the indicated primary antibody (Key Resources Table). Slices were incubated overnight on a rotating plate at 4°C. The next day, slices were washed 4x times for 5 minutes in PBS at room temperature, then incubated with the indicated secondary antibodies in blocking solution (Supplementary Information) for 2h at room temperature. Finally, slices underwent 3x 5 minute washes in PBS to wash off secondary antibodies. Tissue was mounted on slides in PBS and coverslips (Fisherbrand, Cat. No. 1255005) mounted with DAPI Fluoromount-G (Southern Biotech). Slides were imaged using a fluorescent microscope (Keyence BZ-X710) with a 4x objective. Probe placements and viral expression were determined by comparing location to the Allen Mouse Brain Atlas.

### Exclusion Criteria

For electrophysiology experiments, animals were excluded if the injection site was off target. Individual cells were excluded if the mean access resistance before and after current clamp experiments was over 20 MOhm. For photometry and optogenetic experiments, animals were excluded if we observed probes in unintended locations or lack of viral expression. For Kir2.1 experiments, animals were excluded if we did not observe bilateral CeA expression, or if we observed any significant spread outside of the CeA. For behavioral analyses, RI60 sessions were excluded if mice performed fewer than 30 pokes. For all analyses, outliers were excluded using GraphPad Prism’s ROUT method at the most conservative level (0.1%). For analyses conducted outside of GraphPad, we built a python function that mimicked Prism’s function with the same conservative criteria.

### Quantification and Statistical Analysis

#### Statistical analysis

Conventional statistical comparisons (t-tests, ANOVAs, correlations, and linear/non-linear regressions) were performed in GraphPad Prism 10. Specialized analyses used custom Python and R code (photometry processing and signal extraction, electrophysiology extraction and analysis, biophysical modeling in Python; electrophysiology clustering and functional linear mixed-modeling in R; see Key Resources Table). Unless otherwise noted, data are presented as mean ± SEM, or as box-and-whisker plots (Tukey method: box, 25th–75th percentile with median; whiskers extend to the most extreme value within 1.5 × the interquartile range). For all statistical tests, the significance threshold was α = 0.05. In figures, asterisks denote a significant main effect or pairwise comparison (*p<0.05, **p<0.01, ***p<0.001, ****p<0.0001) and pound signs a significant interaction (#p<0.05, ##p<0.01, ###p<0.001, ####p<0.0001); “ns” indicates p≥0.05. n denotes number of mice except in electrophysiology experiments, where it denotes number of cells (number of mice are noted in corresponding figure legends). Before testing, data were assessed for normality (D’Agostino–Pearson and Shapiro–Wilk); nonparametric tests were used when either test detected non-normality (p < 0.05). Outliers were removed using the ROUT method (Q=0.1%) in Prism; for analyses outside Prism, custom Python or R functions reproduced the ROUT/Benjamini-Hochberg test at the same threshold.

#### Group comparisons (Prism)

for comparisons involving two groups, we used unpaired two-tailed t-tests (with Welch’s correction in cases of unequal variances), or Mann-Whitney U-tests when data were non-normal. With three+ groups, we performed one-way ANOVA (Tukey’s). For factorial analysis in Prism, we used two-way ANOVA: ordinary for independent observations (region×stress, virus×stress, cluster×stress, sex×stress), or repeated-measures/mixed-effects models when measurements were repeated within subject (current steps, days, reward number, corticosterone levels, etc) or data were missing (REML, Geisser–Greenhouse correction). Three-way mixed-effects ANOVAs were used for designs with two within-subject factors (weight across age split by sex). Tukey’s post-hoc tests were used for all multiple comparisons. Pearson correlation and simple linear regression were used for associative statistics.

#### Electrophysiology analysis

Effects of stress and projection target on intrinsic features were evaluated with linear mixed-effects models (statsmodels) with random effects for mouse and cell to account for the hierarchical structure of the data. Effects and interactions were tested by Wald F/t tests on model contrasts with Šidák correction. Feature extraction is described in Method Details; clustering and biophysical modeling are described in their respective Method Details sections.

#### Fiber photometry

Data were preprocessed in GuPPy^73^. Briefly, the isosbestic channel was fit to the signal channel by iteratively reweighted least-squares robust linear regression, and ΔF/F was computed as (signal − fitted control)/(fitted control) × 100. ΔF/F was converted to a standard (whole-session) z-score, z(t) = (ΔF/F(t) − μ)/σ, using the session mean and SD. z-scored ΔF/F was used for all peri-event analyses. Spontaneous transients were detected on the whole-session z-score in 15 s moving windows by a two-stage median-absolute-deviation (MAD) procedure: samples exceeding median + 2 × MAD were first removed as high-amplitude artifacts, and transients were then identified as local maxima exceeding median + 3 × MAD of the remaining trace. Transient frequency was the event count divided by recording duration (events/min), and amplitude the z-scored peak height. For comparing group differences in photometry signals, group-level significance was assessed by hierarchical bootstrap over animals^54^. Per-mouse mean traces were computed, and 25,000 bootstrap resamples were drawn by resampling mice with replacement. Pointwise 95% confidence intervals were taken as the 2.5th–97.5th percentiles of the bootstrap distribution and expanded by √(n/(n−1)) to correct small-sample bias. A timepoint was considered significant where the 95% CI excluded zero (for within-group comparisons against baseline) or where the bootstrapped between-group difference CI excluded zero; to control false positives across the time series, significance was required to persist for ≥ 20 consecutive samples (≈0.20 s at ∼102 Hz).

#### Functional Linear Mixed Modeling

The relationship between trial-level behavior variables and TS dopamine photometry signals were calculated using the fastFMM package in R^58^ for rewarded and unrewarded nosepokes. The functional outcome was the trial level photometry trace in a −2 to +5 second window surrounding the event, downsampled to ∼50 Hz. Three models were fit for each predictor (total poke, reward rate, instantaneous poke rate). The first was a group interaction model, photoTrace ∼ group * variable + (1|mouse), fit to the combined stress and control data. In this model, the β(t) for the interaction term tests whether the variable’s coupling to the signal differs between stressed and unstressed groups across the event window. The second and third were group-specific models, photoTrace ∼ variable + (1|mouse), fit for stressed and unstressed data separately. In all models the random mouse intercept accounted for the nested nature of repeated trials from a single mouse. Prior to modeling, each predictor was z-scored using a pooled mean and SD computed across both groups, so that the same standardization was applied to unstressed and stressed traces and their β(t) curves remained on a common scale. Coefficients were considered significant at time points when 95% of their pointwise confidence band excluded zero. To guard against spurious isolated time points, significance was additionally required to last for more than 10 consecutive samples (0.2 seconds). For variables, reward rate and instantaneous poke rate were calculated as causal, exponentially-decaying leaky integrators evaluated at each nosepoke. For a given event, every preceding event of the relevant type contributed an impulse that decayed with the elapsed time since it occurred, so that the value at poke *i* was:

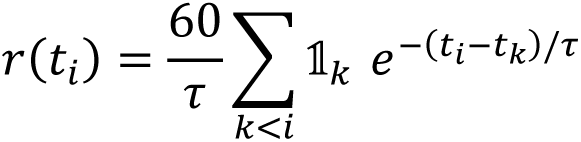

with a decay time constant τ = 60 s, and the leading factor of 60/τ converting the accumulated impulses to events per minute. For reward rate, the indicator 1ₖ was 1 only for rewarded pokes, so the measure reflects the recent density of earned rewards; for instantaneous poke rate, every poke contributed an impulse, yielding the recent density of nose poking regardless of outcome.

#### Behavioral mixture modeling

As the mixture weights are inherently compositional and thus not independent, group differences in composition were tested with a Bayesian Dirichlet regression fit in brms interfaced to Stan. The five session-level weights (π₀–π₄; floored at 10⁻⁶ and renormalized to avoid boundary values) were entered as a Dirichlet-distributed response, with sessions as the unit of observation and a per-mouse random intercept accounting for repeated sessions (weights ∼ group + day + (1 | mouse), Dirichlet family with the default multinomial-logit link and the shortest-latency “motor” component as the reference category), using brms’s default weakly-informative priors.

Models were fit with four chains of 4,000 iterations each (1,000 warmup; 12,000 post-warmup draws; adapt_delta = 0.95); convergence was confirmed by R^ < 1.01 and bulk effective sample sizes > 1,000 for all parameters. Because Dirichlet coefficients are log-ratios relative to the reference category and are not directly interpretable, group effects are reported as marginal contrasts on the proportion scale: for each component we computed the posterior difference in model-predicted weight between groups (stressed – unstressed), averaged over the empirical distribution of training days at the population level (G-computation/standardization). Effects are summarized by the posterior mean, its 95% credible interval (CrI), and the probability of direction (pd, the posterior probability that the effect has the sign of the point estimate); a component was considered credibly different between groups when its 95% CrI excluded zero (equivalently pd ≥ 0.975), and for tabulation alongside conventional tests we report a two-sided analog p = 2 × (1 − pd).

#### scRNA-seq re-analysis

We reanalyzed single-cell RNA-sequencing data from Wang et al.^40^ (GEO accession GSE213828). After filtering for quality—retaining cells with more than 2,000 genes detected and fewer than 25% ERCC reads (1,576 of 1,824 neurons)—counts were normalized to 10,000 per cell and log-transformed in Scanpy, while raw counts were retained for statistical testing. Neurons were assigned to putative projection groups using the marker genes identified by Wang et al. as most predictive of axonal projection (putative SNL, Drd1⁺/Sema3c⁻; putative BNST, Dlk1⁺; putative PAG, Crh⁺). Expression of Kcnq2, Kcnq3, and Kcnq5 was compared across these groups using Kruskal–Wallis tests followed by Dunn’s post hoc tests with Benjamini–Hochberg correction. To jointly test the effects of projection and gene, we fit a negative binomial generalized linear model and assessed main effects and interactions by likelihood-ratio tests comparing nested models; to account for excess zeros in Kcnq3 and Kcnq5, we additionally fit a zero-inflated negative binomial model.

---

**Supplementary Figure 1 (Related to Figure 1).**
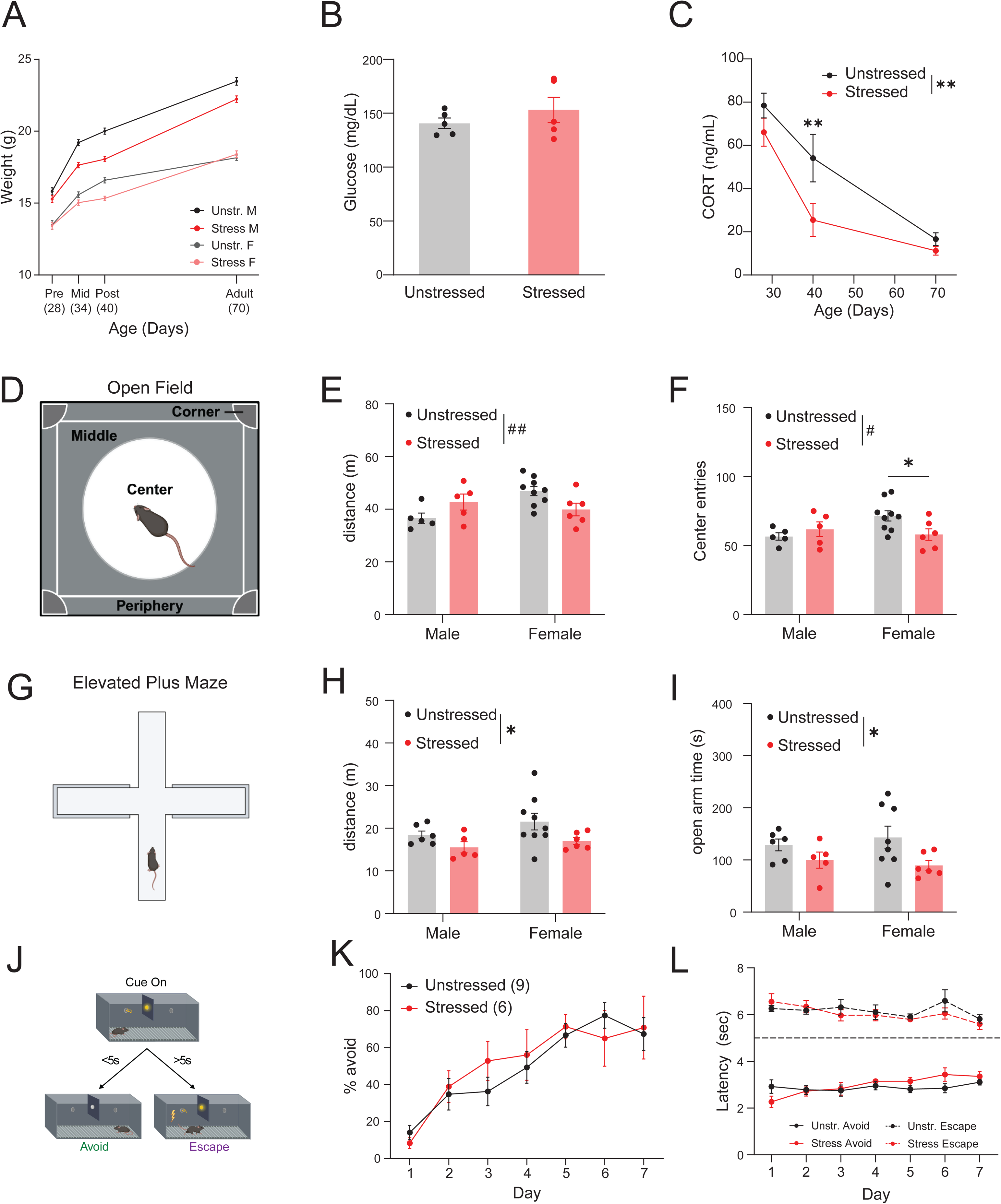
Adolescent stress leads to persistent physiological changes and anxiety-like behavior but does not change ability to learn from shock. (A) Weight of stressed vs. unstressed males and females before (P28), during (P34), and immediately after (P40) adolescent stress, and at the beginning of the experimental period (P70; Three-way repeated measures mixed effects analysis followed by two-way RM mixed effects analyses split by sex, see Table S2) (B) Groups did not differ in blood glucose concentration (Unpaired t-test, p = 0.3574, n = 5/group) (C) Stressed mice had blunted levels of circulating corticosterone (as measured at ZT16), particularly immediately after stress (P40; two-way repeated measures mixed effects model (REML), age x stress with Tukey’s post-hoc tests: age: ****p < 0.0001, stress, **p = 0.0011, interaction, p = 0.0560; post-hoc tests: P28 (n = 4-6/group): p = 0.5041, P40 (n = 6-7/group): p = 0.0031, P70 (n = 21-23/group), p = 0.5508) (D) Schematic of the open field arena with a bright circle of light in the middle to increase the center’s anxiogenic properties. (E) There were sex-dependent effects of adolescent on total distance traveled in the open field: stress males moved more, whereas stressed females moved less (2-way ANOVA + Tukey’s post-hoc, stress: p = 0.84488, sex: p = 0.1208, interaction: ^##^p = 0.0095; post-hoc tests (stressed - unstressed): M, p = 0.1877, F, p = 0.0527, n = 5-9 mice/sex/group) (F) There were sex-dependent effects of adolescent stress on anxiety-like behavior (entries into the center region) in the open field: males did not display a group difference, while stressed females entered fewer times (2-way ANOVA + Tukey’s post-hoc, sex: p = 0.2037, stress: p = 0.3370, interaction: ^#^p = 0.0387; post-hoc tests (stressed - unstressed): M, p = 0.6807, F, *p = 0.0420, n = 5-9 mice/sex/group) (G) Schematic of the elevated plus maze with two closed and two open arms. (H) There was a decrease in total locomotion in the elevated plus maze in stressed mice (2-way ANOVA + Tukey’s post-hoc, stress: *p = 0.0302, sex: p = 0.1646, interaction: p = 0.6164; post-hoc tests (stressed - unstressed): M, p = 0.4274, F, p = 0.0835, n = 5-8 mice/sex/group). (I) There was an increase in anxiety-like behavior (time spent in the open arms) in the elevated plus maze in stressed mice (2-way ANOVA + Tukey’s post-hoc, stress: *p = 0.0253, sex: p = 0.9031, interaction, p = 0.4851; post-hoc tests (stressed - unstressed): M, p = 0.4642, F, p = 0.0572, n = 5-8 mice/sex/group). (J) Schematic of active avoidance paradigm. Mice had 5 seconds after cue onset to cross to the opposite chamber of the shuttlebox, otherwise they received a footshock (0.4 mA) until they crossed (∼1s after shock onset). (K) Percent shocks avoided across days of learning in the active avoidance paradigm. No effects of adolescent stress were observed (2-way RM mixed effects analysis, day: ****p < 0.0001, stress:, p = 0.6112, interaction: p = 0.7895, n = 4-9 mice/day/group). (L) Latencies to shuttle for avoid and escape trials across days in the active avoidance paradigm. No effects of adolescent stress were observed (escape: two-way RM mixed-effects analysis, no main effects or interactions, p > 0.05 for all; latency: two-way RM mixed-effects analysis, no main effects or interactions, p > 0.05 for all). All data presented as mean ± SEM. ^∗^*p* < 0.05; ^∗∗^*p* < 0.01; ^∗∗∗^*p* < 0.001; ^∗∗∗∗^*p* < 0.0001 for main effects or pairwise comparisons; ^#^*p* < 0.05; ^##^*p* < 0.01; ^###^*p* < 0.001; ^####^*p* < 0.0001 for interactions. See also Table S2 for detailed statistics.

**Supplementary Figure 2 (Related to Figure 1).**
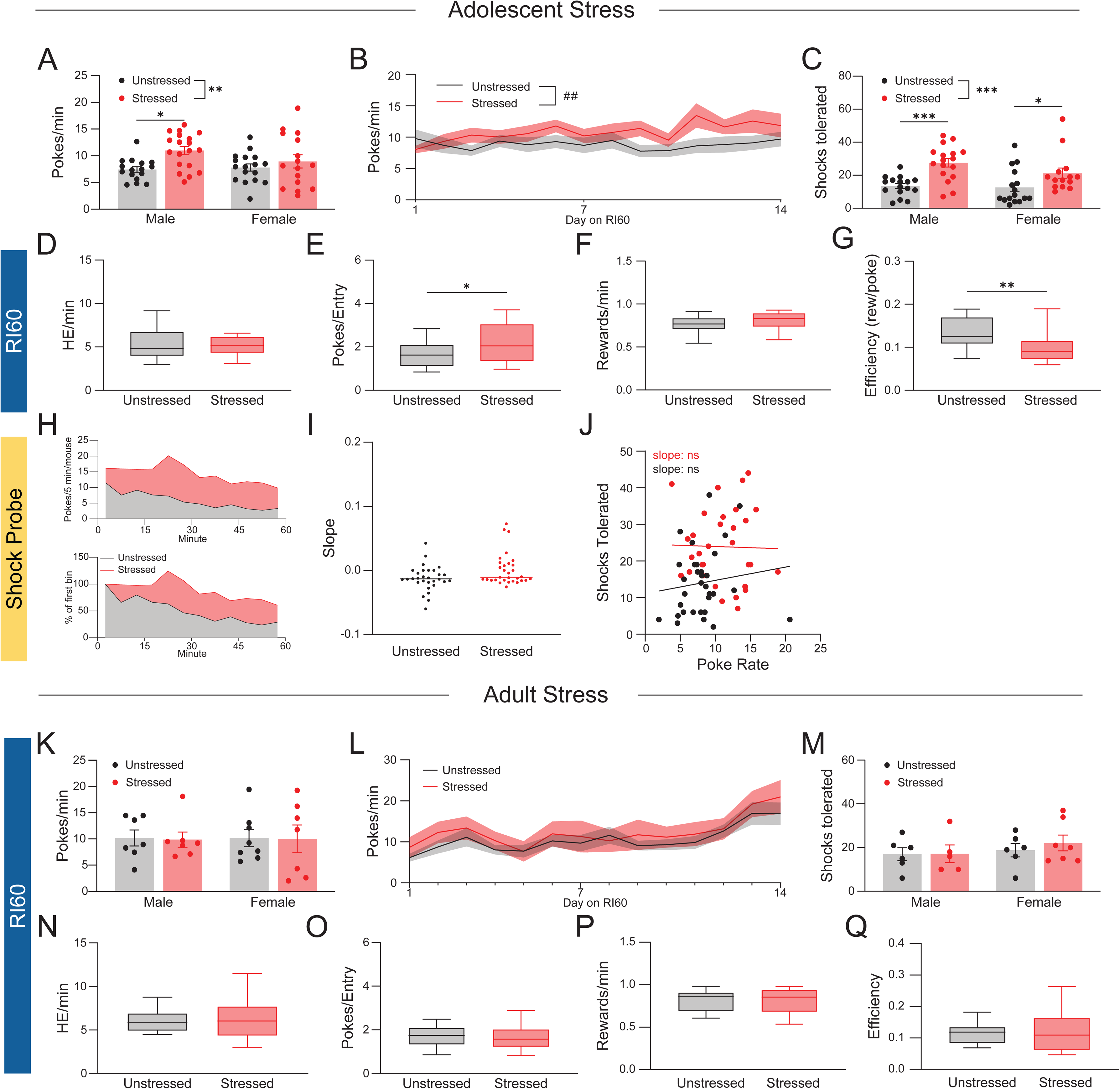
Adolescent stress effects on RI60 training and shock probe behavior in adolescents and adults. (A) There were no effects of sex on RI60 nosepoke (two-way ANOVA with Tukey’s post-hoc tests: sex: p = 0.8595, stress: **p = 0.0068, interaction: p = 0.6400; post-hoc tests (stressed - unstressed): M, *p = 0.0484, F, p = 0.1958; n = 15-19 mice/sex/group). (B) Nosepoke rate across days of RI60 training. Groups started at similar nosepoke rates, but stressed mice nosepoked more by the end of RI60 training (2-way RM mixed-effects analysis (day x stress): day: p = 0.3259, stress: p = 0.1077, interaction: ^##^p = 0.0095, n = 20-36 mice/day/group) (C) There were no effects of sex on punishment-resistant reward-seeking in the shock probe (two-way ANOVA with Tukey’s post-hoc tests: sex, p = 0.1644, stress: ****p < 0.0001, interaction, p = 0.2705; post-hoc tests (stressed - unstressed): M, ***p = 0.0002, *F, p = 0.0238; n = 15-19 mice/sex/group). (D-G) Group comparisons of port entry rate (D, Mann-Whitney exact test, p = 0.7250), pokes per entry (E, Mann-Whitney exact test, *p = 0.0393), reward rate (F, Mann-Whitney exact test, p = 0.0909), efficiency (number of rewards earned per poke, G, Mann-Whitney exact test, **p = 0.0018) during RI60 training, n = 34-35 mice/group. (H) Nosepoke behavior during the shock probe sessions. Top, total number of nosepokes across the shock probe in 5 minute bins, pooled across all mice and split by group; bottom, the same data but normalized to poke rate in the first bin. (I) Individual-level slopes of nosepoke rate across the shock probe, calculated as (pokes in first bin/pokes in last bin)/number of bins. No effects of adolescent stress were observed (Mann-Whitney exact test, p = 0.0751, n = 30-31 mice/group) (J) Correlations between RI60 nosepoke rate and number of shocks tolerated during the shock probe (unstressed: R^2^ = 0.01710, p = 0.4756; stressed: R^2^ = 0.0005279, p = 0.9024, n = 31-32/group). The lack of correlation in either group suggests that elevated nosepoke rates are not sufficient to explain elevated punishment-resistant reward seeking in the shock probe. (K) There were no effects of group or sex on nosepoke rate when stress was applied in adulthood instead of adolescence (2-way ANOVA, no significant main effects or interactions, p > 0.05, n = 7-8 mice/sex/group). (L) There were no differences in nosepoke rates across days after adult stress (2-way RM mixed-effects analysis (day x stress): main effect of day, p < 0.0001, no main effect of stress, p = 0.5291, no significant interaction, p = 0.9751, n = 9-15 mice/day/group). (M) There were no effects of group or sex on punishment-resistant reward-seeking following adult stress (2-way ANOVA, no significant main effects or interactions, p > 0.05, n = 5-7 mice/sex/group) (N-Q) There were no effects of adult stress on port entry rate (N, Welch’s t-test, p = 0.7124), pokes/entry (O, Welch’s t-test, p = 0.8188), reward rate (P, unpaired t-test, p = 0.8054), efficiency (Q, Mann-Whitney exact test, p = 0.9349) during RI60 training, n = 15 mice/group. A-C, K-M are presented as mean ± SEM. D-G, N-Q are presented as box and whisker plots (IQR, median line, Tukey whisker method, points outside whiskers not shown). I presented as line at median. J presented as scatter plot with linear regression fit. ^∗^*p* < 0.05; ^∗∗^*p* < 0.01; ^∗∗∗^*p* < 0.001; ^∗∗∗∗^*p* < 0.0001 for main effects or pairwise comparisons; ^#^*p* < 0.05; ^##^*p* < 0.01; ^###^*p* < 0.001; ^####^*p* < 0.0001 for interactions; n.s., not significant. See also Table S2 for detailed statistics.

**Supplementary Figure 3 (Related to Figure 1).**
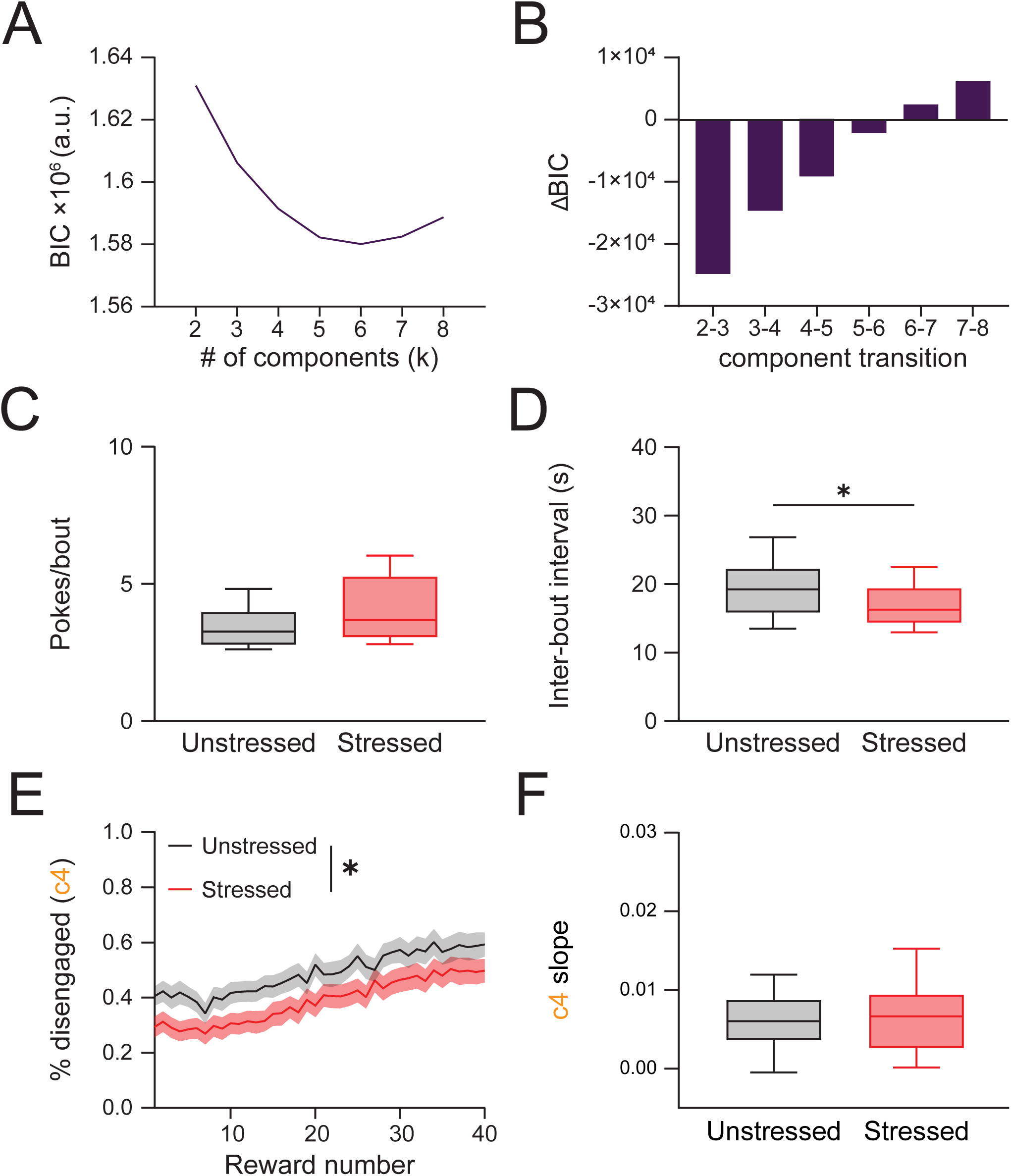
Lognormal mixture modeling of IPIs reveals decreased disengagement in adolescent stress mice. (A-B) Model fit (A) and change in fit (B) across number of components (C) Stress did not affect the number of pokes within nosepoking bouts (defined as nosepokes from components 0-2; Mann-Whitney exact test, p = 0.0571, n = 33-34 mice/group) (D) Stressed mice had shorter inter-bout intervals than unstressed controls (components 3-4; unpaired t-test, *p = 0.0295, n = 35 mice/group) (E) Stressed mice spent a lower percentage of time disengaged (component 4; c4) across the average session (2-way RM mixed-effects analysis: reward: ****p < 0.0001, stress: *p = 0.0168, interaction: p = 0.7541; n = 32-35 mice/reward/group), parceled by reward number to account for the varying total time of most sessions and trimmed to 40 rewards as most sessions reached 40 rewards, but many did not reach 50. (F) Stressed and unstressed mice displayed satiation effects (increased disengagement across the session represented by c4 slope) to similar degrees (unpaired t-test, p = 0.5509). † indicates slope is significantly different from 0 (one-sample t-test vs. 0, unstressed and stressed both p < 0.0001), n = 35 mice/group. E is presented as mean ± SEM. C-D, F are presented as box and whisker plots (IQR, median line, Tukey whisker method, points outside whiskers not shown). ^∗^*p* < 0.05; ^∗∗^*p* < 0.01; ^∗∗∗^*p* < 0.001; ^∗∗∗∗^*p* < 0.0001 for main effects or pairwise comparisons; ^#^*p* < 0.05; ^##^*p* < 0.01; ^###^*p* < 0.001; ^####^*p* < 0.0001 for interactions; n.s., not significant. See also Table S2 for detailed statistics.

**Supplementary Figure 4 (Related to Figure 2).**
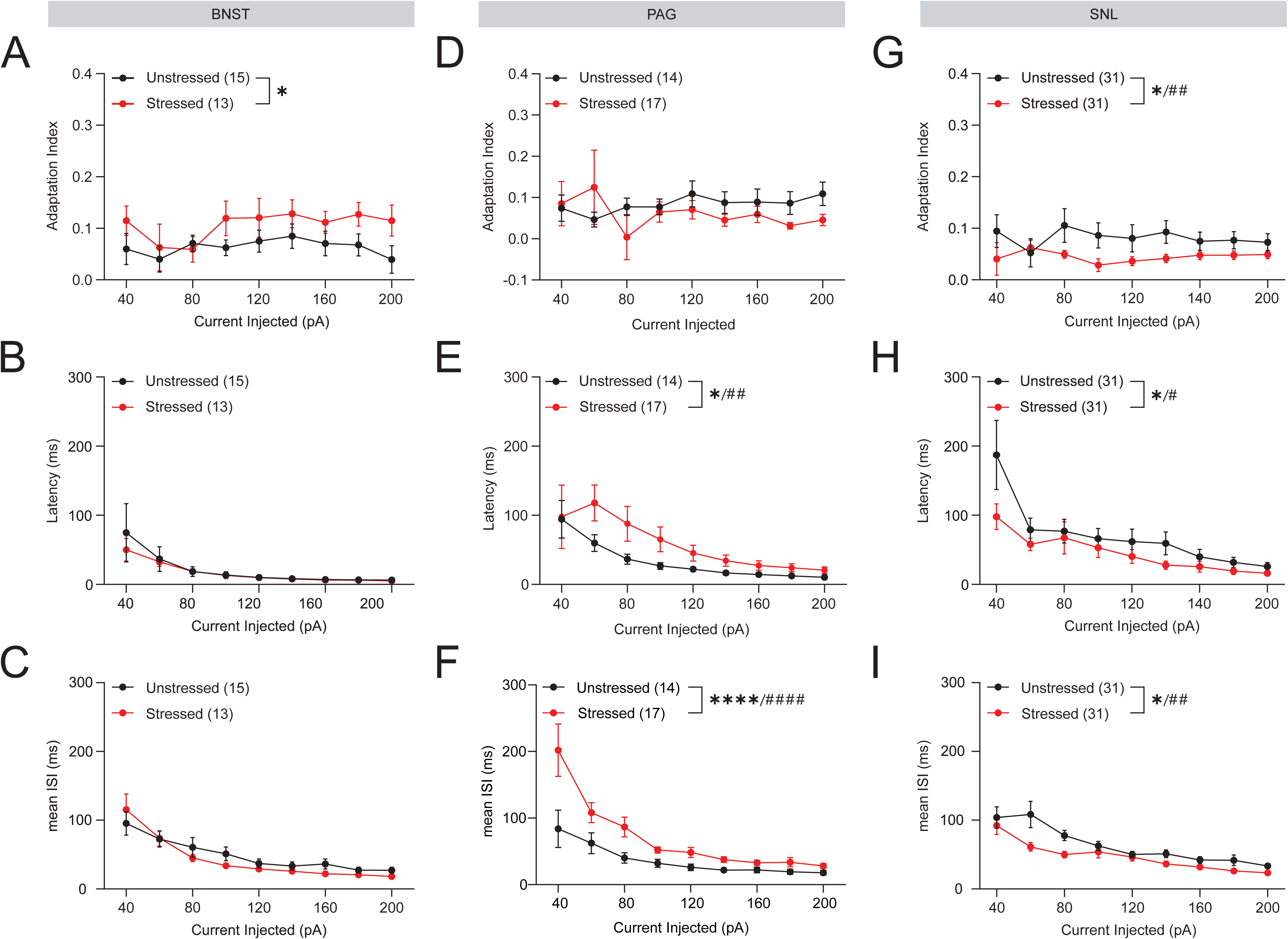
Spike train properties of CeA neurons that project to the BNST, PAG, or SNL. (A-C) Adaptation index (A), latency to spike (B), and mean inter-spike interval (C) for BNST-projecting CeA neurons. Stress increased adaptation index but did not affect the other properties in the BNST projection (Adaptation: stress, *p = 0.0295; current, p = 0.8980; interaction p = 0.8680; Latency: stress, p = 0.2868, current, ****p < 0.0001, interaction, p = 0.0505; Mean ISI: stress, p = 0.4693, current, ****p < 0.0001, interaction, p = 0.4532, n = 3 mice, 13-15 cells/group). (D-F) Adaptation index (D), latency to spike (E), and mean inter-spike interval (F) for PAG-projecting CeA neurons. Stress did not affect adaptation index but increased latency to fire and mean ISI in the PAG projection (Adaptation: stress, p = 0.2645, current, p = 0.8030, interaction, p = 0.3780; Latency: stress, *p = 0.0186, current, ****p < 0.0001, interaction, ^##^p = 0.0059; Mean ISI: ***stress, p = 0.0008, current, ****p < 0.0001, interaction, ^####^p < 0.0001; n = 2-3 mice, 14-17 cells/group). (G-I) Adaptation index (G), latency to spike (H), and mean inter-spike interval (I) for SNL-projecting CeA neurons. Stress decreased adaptation index, decreased latency to fire, and decreased mean ISI in the SNL projection (Adaptation: stress, *p = 0.0173, current, p = 0.4910, interaction, ^##^p = 0.0035; Latency: *p = 0.0463, current, ****p < 0.0001, interaction, ^#^p = 0.0182; Mean ISI: *stress, p = 0.0187, current, ****p < 0.0001, interaction, ^##^p = 0.0025; n = 5-6 mice, 31 cells/group). Hierarchical linear mixed-effects models were used for all analyses. All data are presented as mean ± SEM. ^∗^*p* < 0.05; ^∗∗^*p* < 0.01; ^∗∗∗^*p* < 0.001; ^∗∗∗∗^*p* < 0.0001 for main effects or pairwise comparisons; ^#^*p* < 0.05; ^##^*p* < 0.01; ^###^*p* < 0.001; ^####^*p* < 0.0001 for interactions. See also Table S2 for detailed statistics.

**Supplementary Figure 5 (Related to Figure 3).**
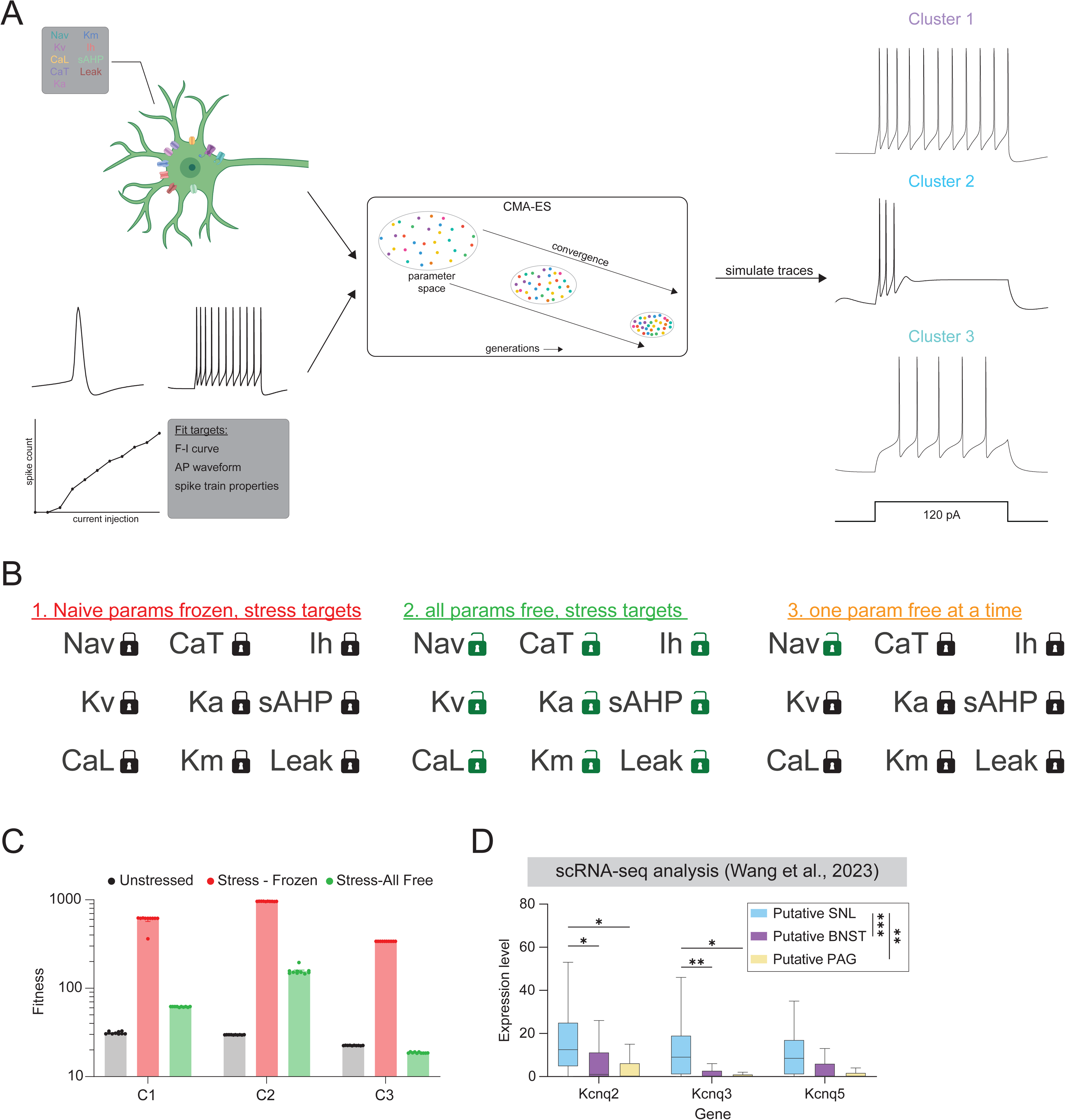
Biophysical modeling identifies M-type potassium currents as a candidate mechanism for stress-induced hyperexcitability across clusters. (A) Schematic of NEURON model strategy. Simulated neurons of each cluster were fit to unstressed data (left) using a CMA-ES optimization algorithm (middle). This strategy produced spike trains that resembled all three clusters (right). (B) Schematic of the fitting strategy for stressed data. Three fits were run: 1) a frozen fit, where all parameters were held at unstressed levels and fit to stressed data, which set the upper bound for error; 2) an “all-free” fit where all parameters were allowed to vary, this set the lower bound for error; 3) models where a given parameter was free to vary but all other parameters were held at unstressed levels (C) Fitness of each model. Unstressed: free parameters on unstressed data; Stress-frozen: frozen unstressed parameters on stressed targets; Stress-all free: free parameters on stressed targets. (D) Reanalysis of single cell RNA-seq data showing that putative CeA→SNL neurons express higher levels of Kv7/KCNQ subunits than putative CeA→BNST or putative CeA→PAG neurons (Kruskal-Wallis test + Dunn’s (BH-adjusted) post-hoc tests. Kcnq2: projection: *p = 0.0118, SN vs. BNST: *p = 0.0107, SN vs. PAG: *p = 0.0107; Kcnq3: projection: **p = 0.0024, SN vs. BNST: **p = 0.0016, SN vs. PAG: *p = 0.0240; Kcnq5: projection: p = 0.1336). ^∗^*p* < 0.05; ^∗∗^*p* < 0.01, data presented as box-and-whisker plots (IQR, median, Tukey whiskers, points outside whiskers not shown).

**Supplementary Figure 6 (Related to Figure 4).**
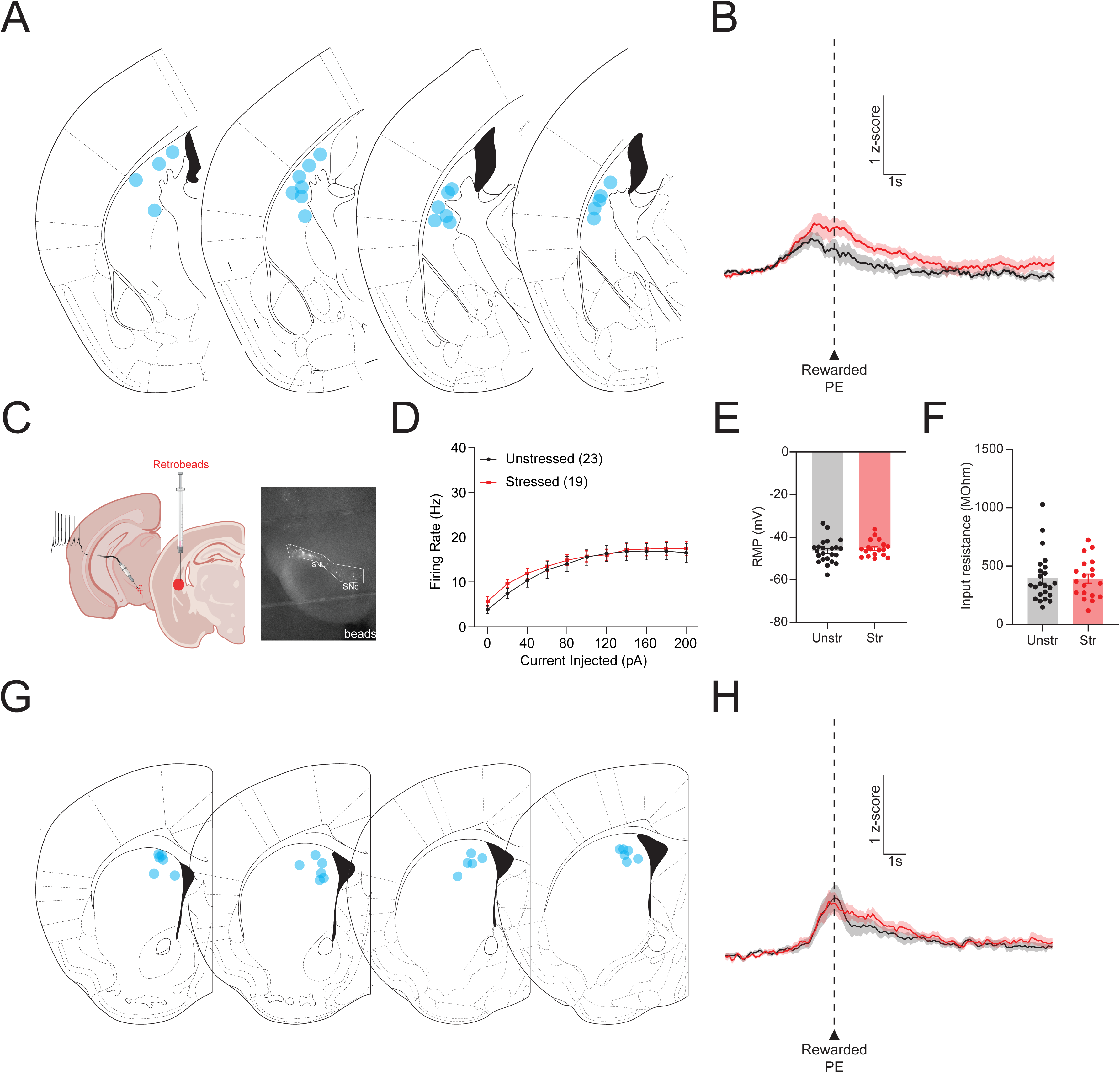
DMS and TS photometry, SNL→TS dopamine neuron electrophysiology. (A) Map of probe locations for TS dopamine fiber photometry. (B) TS dopamine response to rewarded port entry. No significant differences across groups were detected. (C) Schematic for whole-cell patch-clamp recordings of TS-projecting SNL/SNc neurons. Red retrobeads were injected into the TS and intrinsic properties of SNL dopamine neurons were recorded. (D) Firing-current curve for SNL→TS neurons. No differences were detected between groups (stress: p = 0.6468, current: ****p < 0.0001, interaction: p = 0.9478). (E-F) Resting membrane potential (E, p = 0.3928) and input resistance (F, p = 0.6360) of SNL→TS dopamine neurons were not changed by stress. (G) Map of probe locations for DMS dopamine fiber photometry. (H) DMS dopamine response to rewarded port entry. No significant differences across groups were detected. All data are presented as mean ± SEM. All electrophysiology data was analyzed using hierarchical linear mixed-effects modeling and n = 4 mice, 18-23 cells/group for all SNL→TS analyses. ^∗^*p* < 0.05; ^∗∗^*p* < 0.01; ^∗∗∗^*p* < 0.001; ^∗∗∗∗^*p* < 0.0001 for main effects or pairwise comparisons; ^#^*p* < 0.05; ^##^*p* < 0.01; ^###^*p* < 0.001; ^####^*p* < 0.0001 for interactions; n.s., not significant. See also Table S2 for detailed statistics.

**Supplementary Figure 7 (Related to Figure 5).**
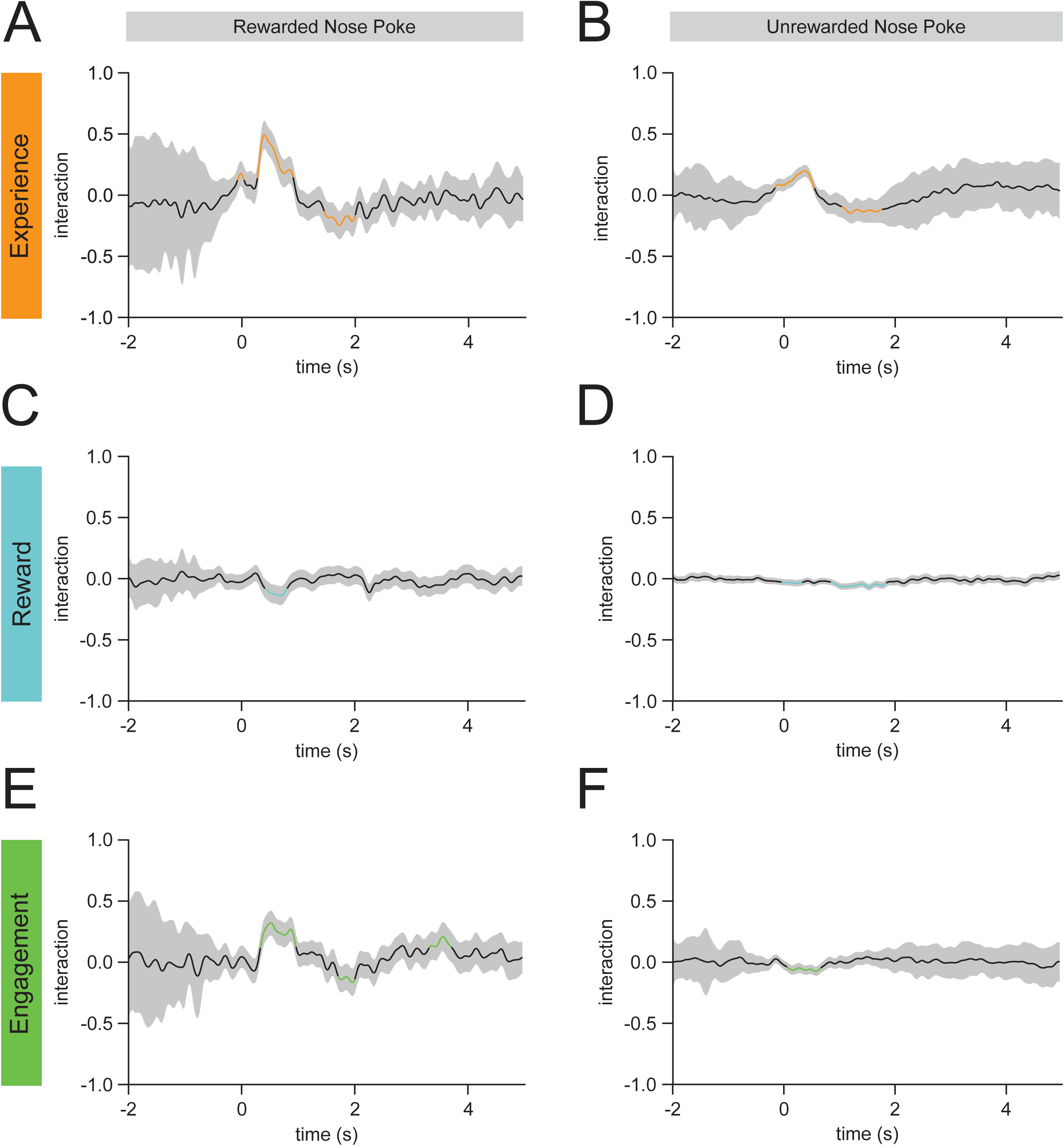
Interaction models for Figure 5 FLMMs. For all group interaction models, significant divergence from 0 indicates that stress vs. unstressed groups differ in encoding. These models were used to generate the purple bars (unstressed ≠ stressed) in Figure 5. (A-B) Group interaction model for effects of total nosepoke experience on rewarded (A) and unrewarded (B) nosepoke responses. (C-D) Group interaction model for effects of recent reward rate on rewarded (C) and unrewarded (D) nosepoke responses. (E-F) Group interaction model for effects of recent nosepoke rate on rewarded (E) and unrewarded (F) nosepoke responses.

**Supplementary Table 1:** condensed chronic unpredictable stress paradigm.

| Day | Morning (9:00) |  | Afternoon (2:00) |  | Overnight (6:00-7:00) |
| --- | --- | --- | --- | --- | --- |
| 1 |  |  |  |  | *Food dep* |
| 2 | New Partner | 3 hr | Tilt | 3 hr | *Isolation* |
| 3 | Stroboscope | 5 hr | Crowding | 3 hr | *Lights off* |
| 4 | Lights on | 3 hr | Restraint | 1 hr | *Tilt* |
| 5 | Swim | 5 min | Lights on | 3 hr | *Lights off* |
| 6 | Stroboscope | 5 hr | Crowding | 3 hr | *Wet bedding* |
| 7 | New Partner | 3 hr | Restraint | 1 hr | *Food dep* |
| 8 | Crowding, light on | 3 hr | Tilt, wet bedding | 3 hr | *Isolation* |
| 9 | Restraint | 1 hr | Swim | 5 min | *Tilt* |
| 10 | Stroboscope | 5 hr | Lights on | 3 hr | *Wet bedding* |
| 11 | Crowding, light on | 3 hr | New Partner | 3 hr | *Food dep* |
| 12 | Stroboscope, wet bed | 5 hr | Restraint | 1 hr | *Isolation* |

**Table S2.**
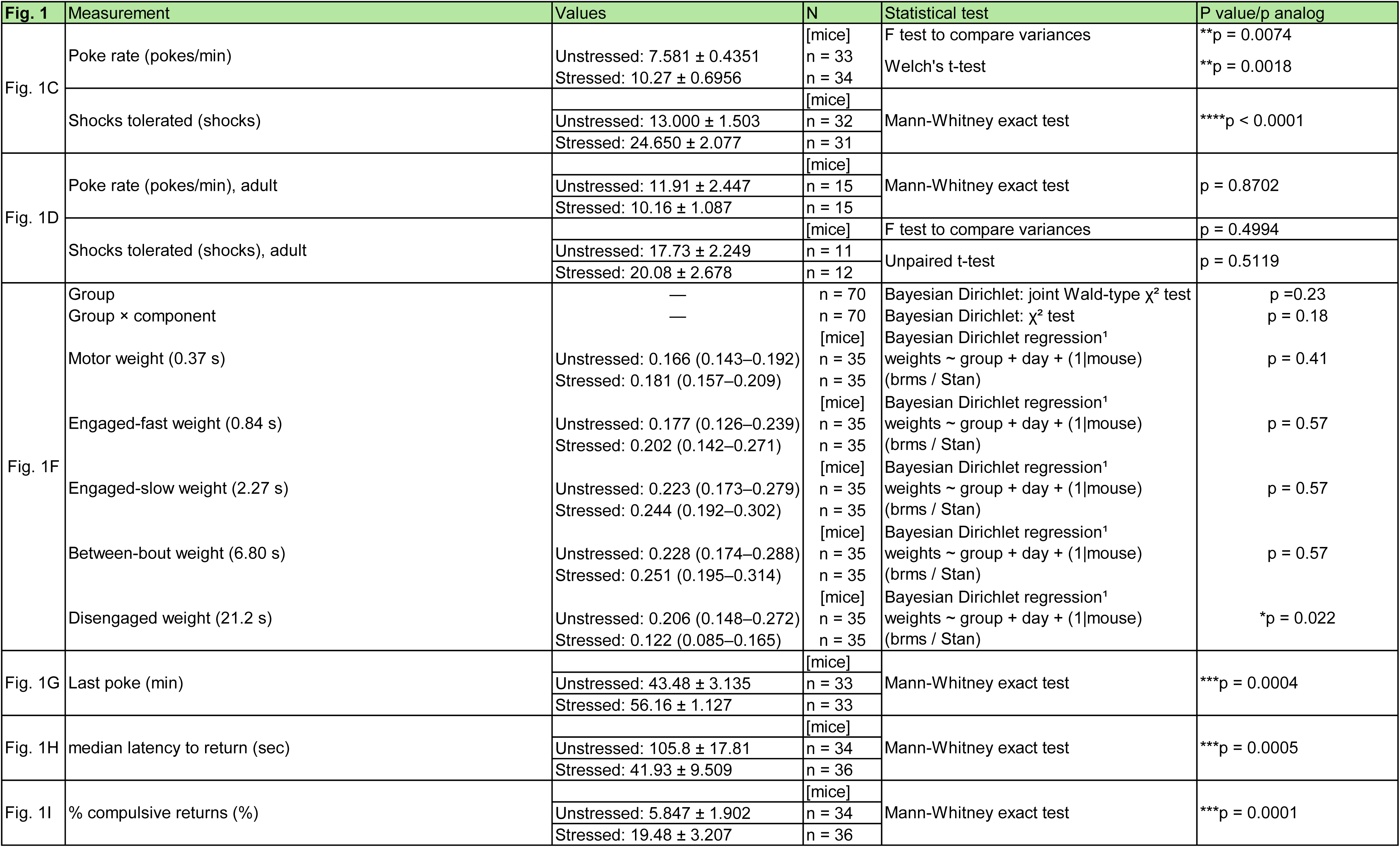
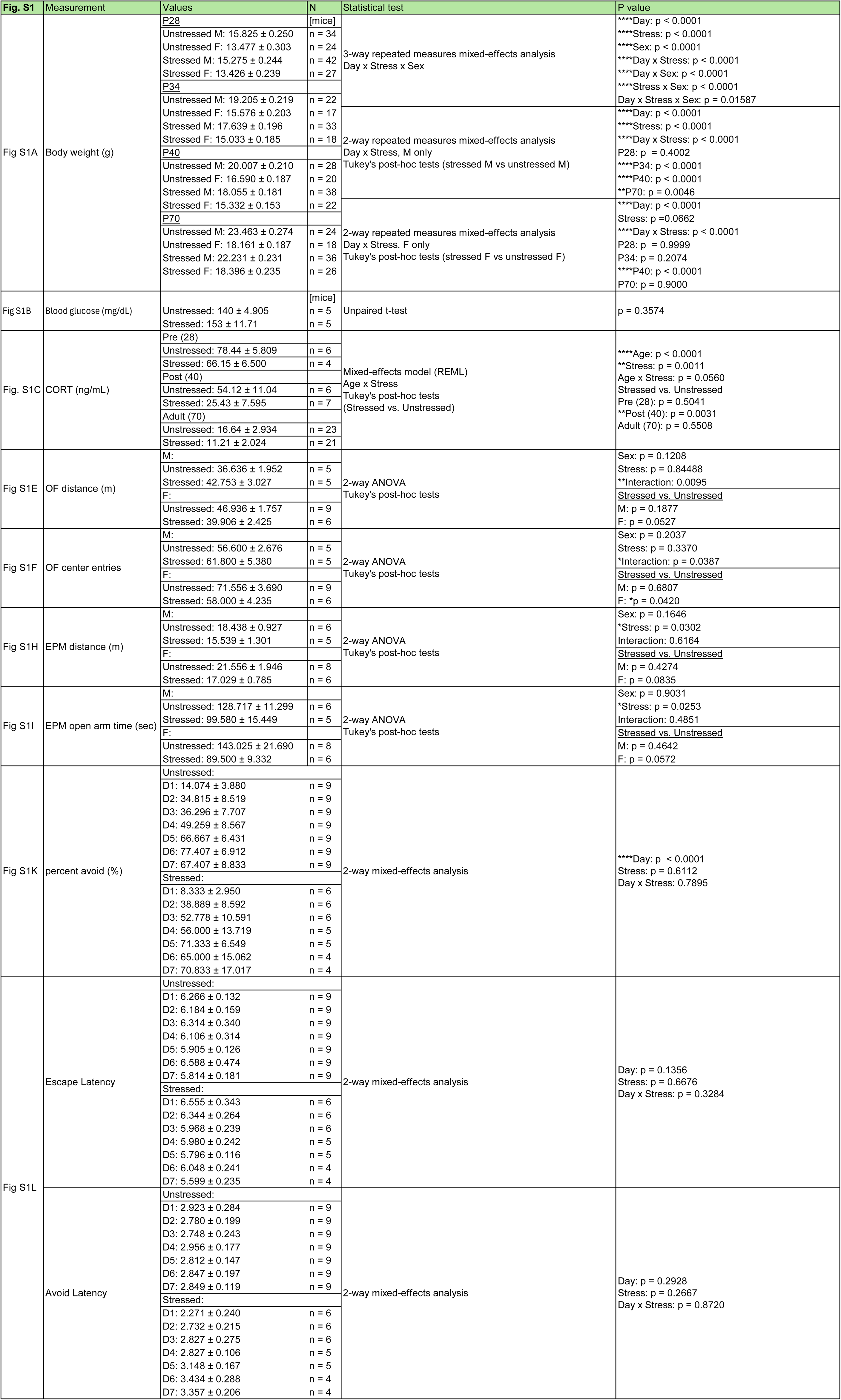
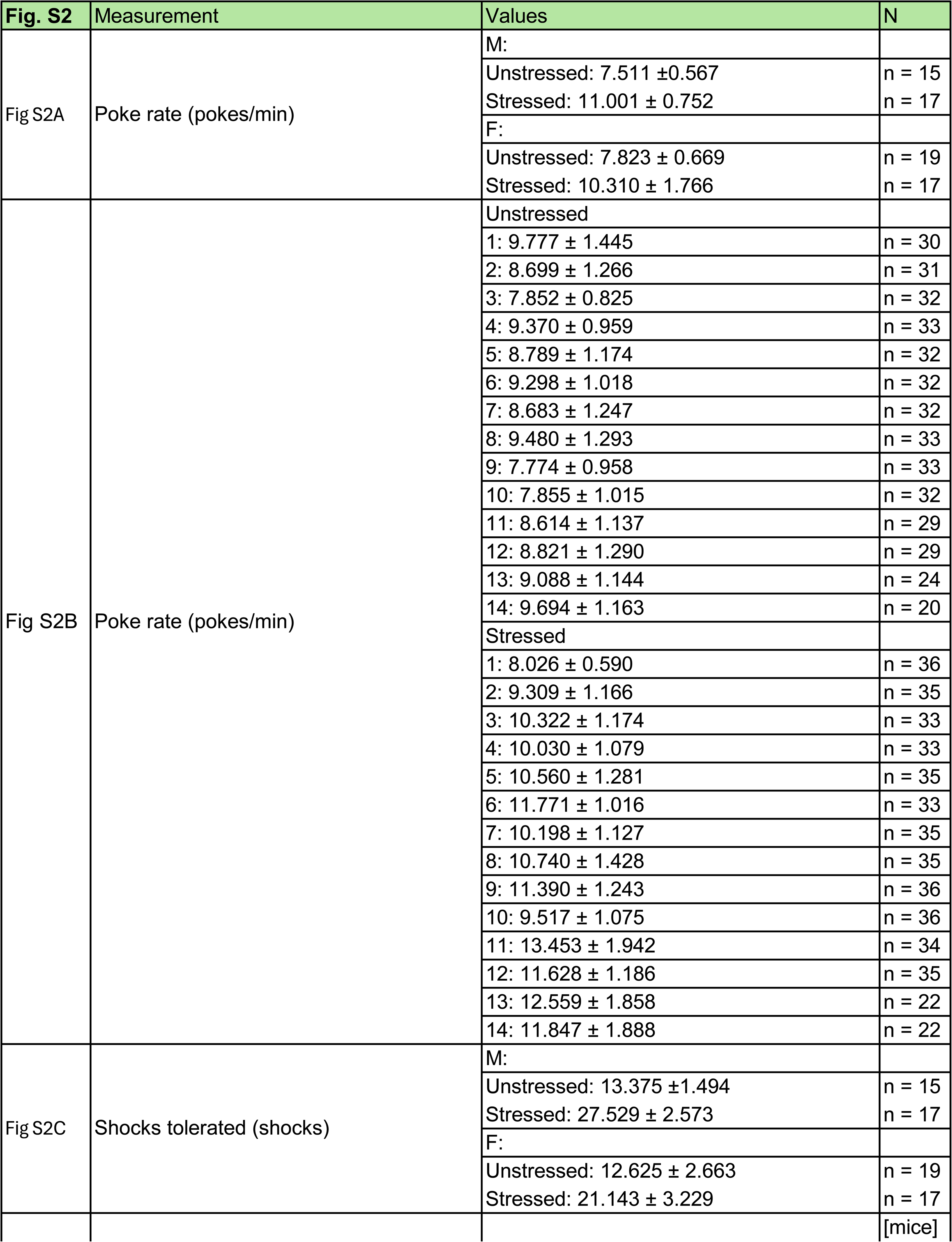
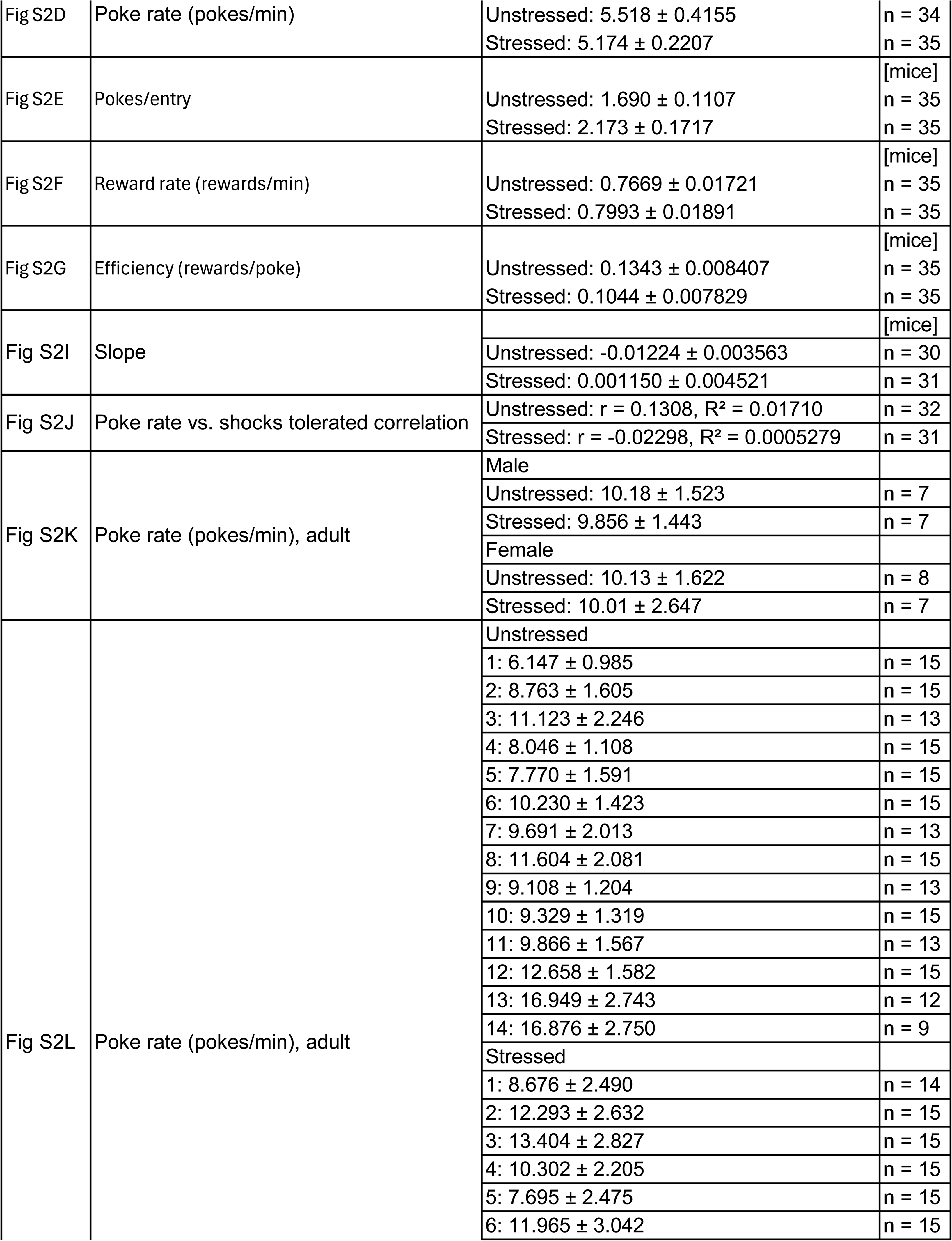
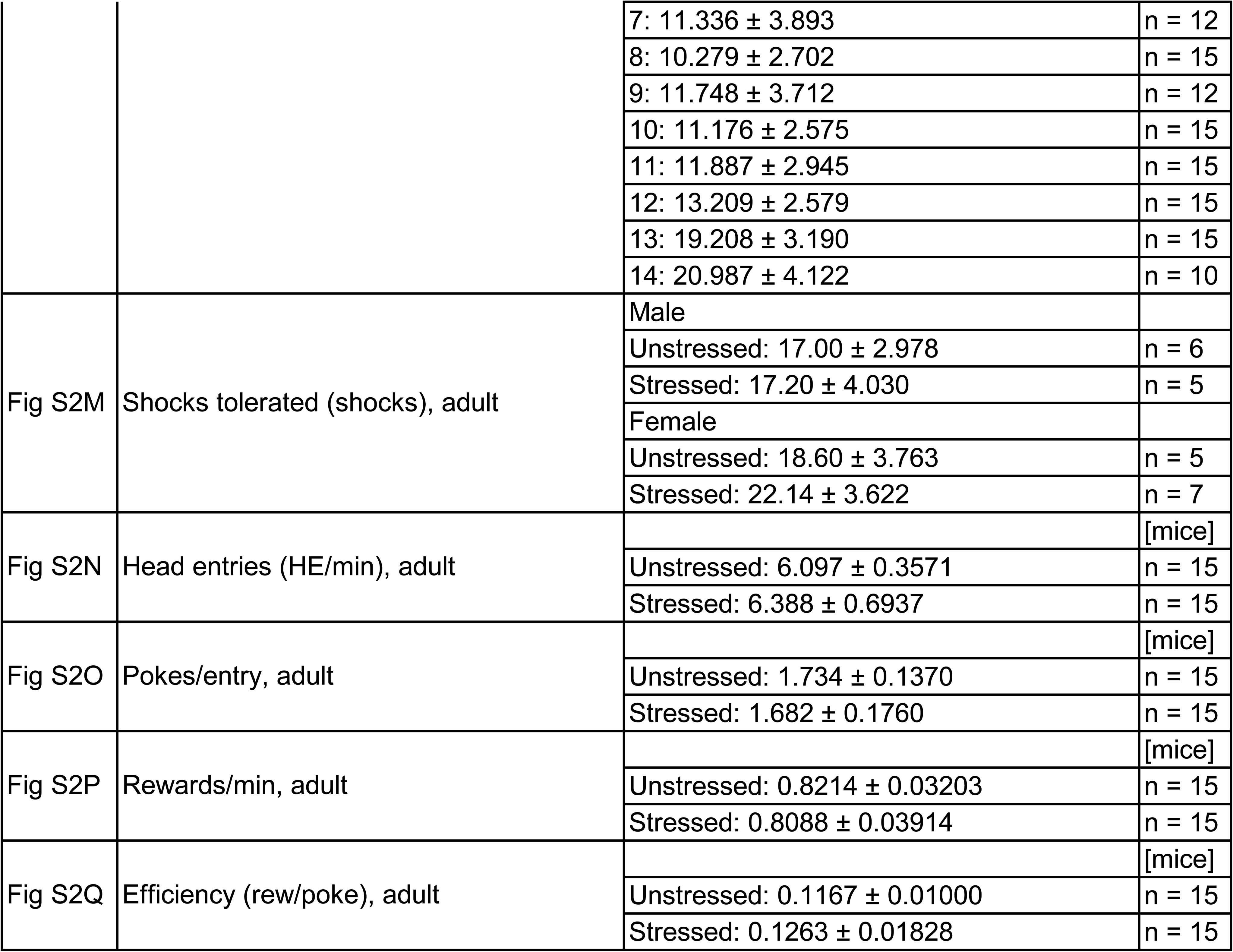
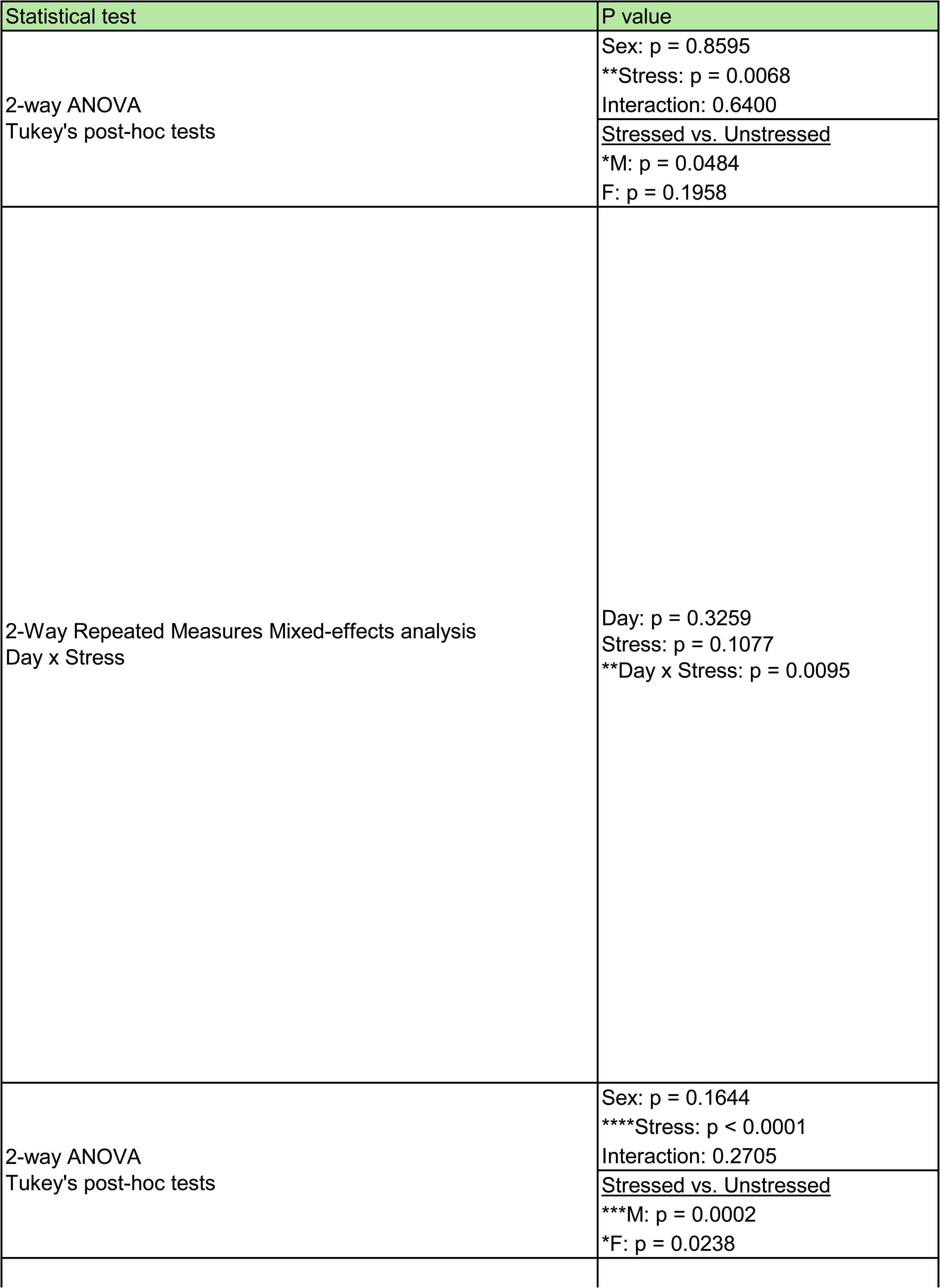
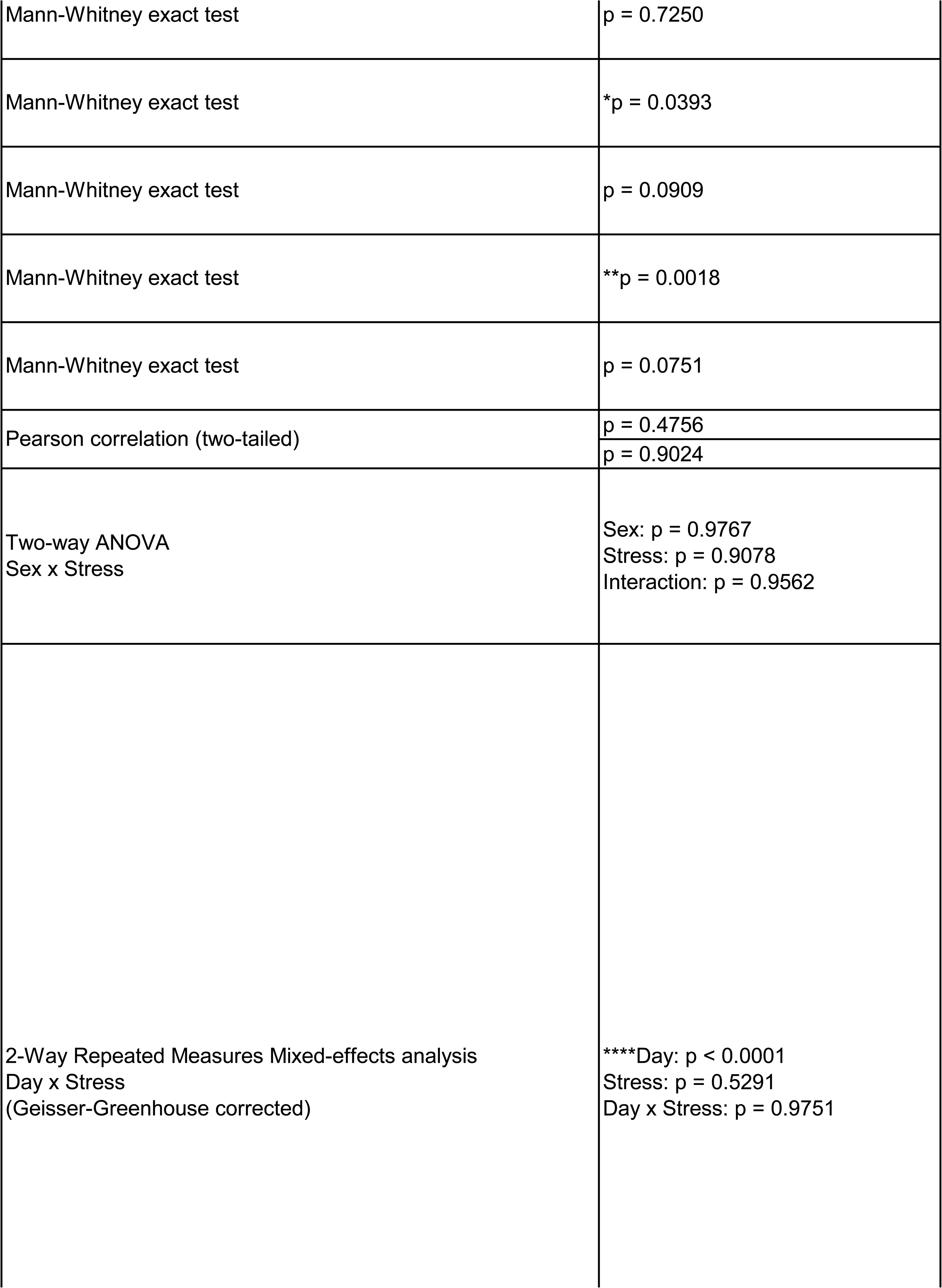
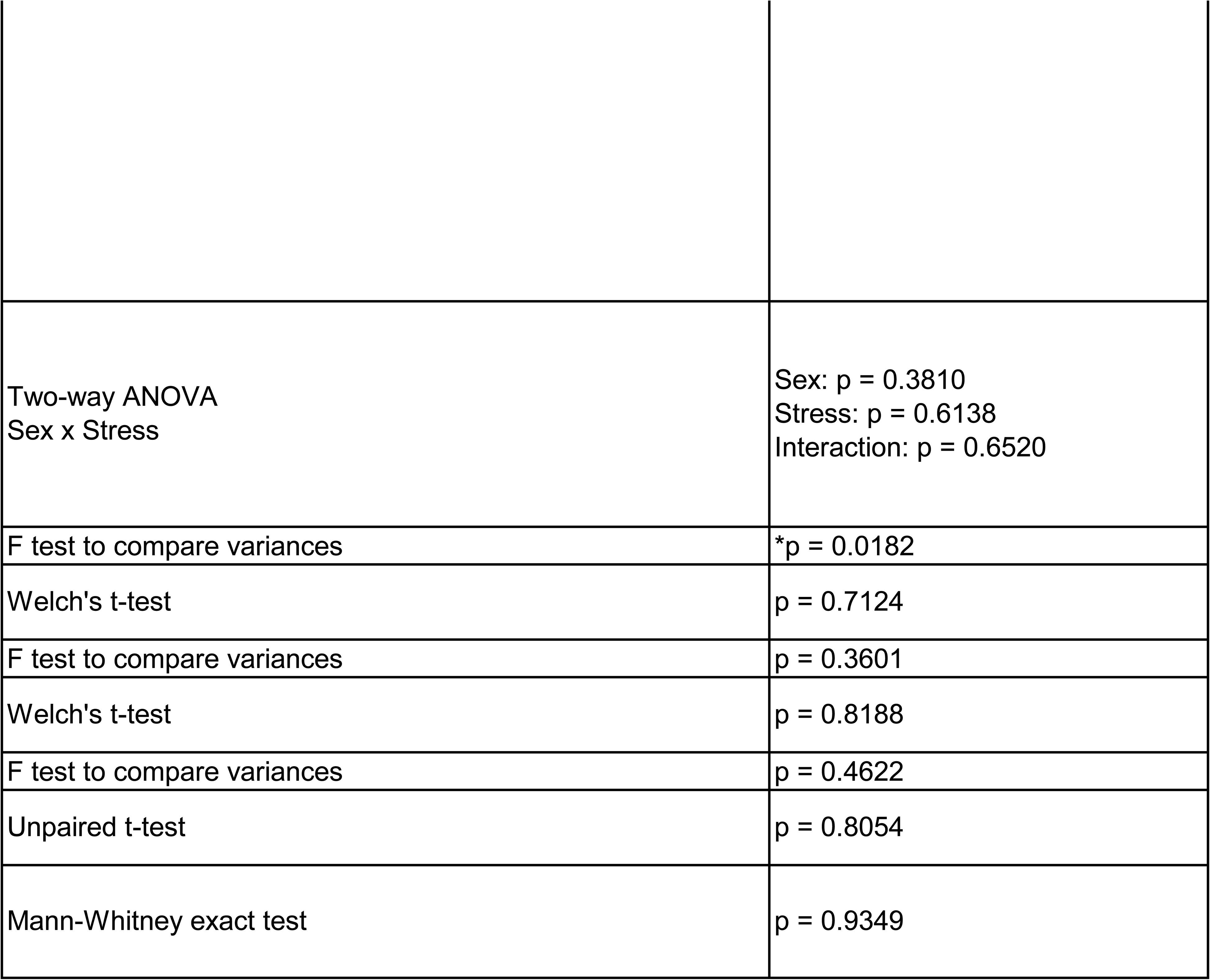
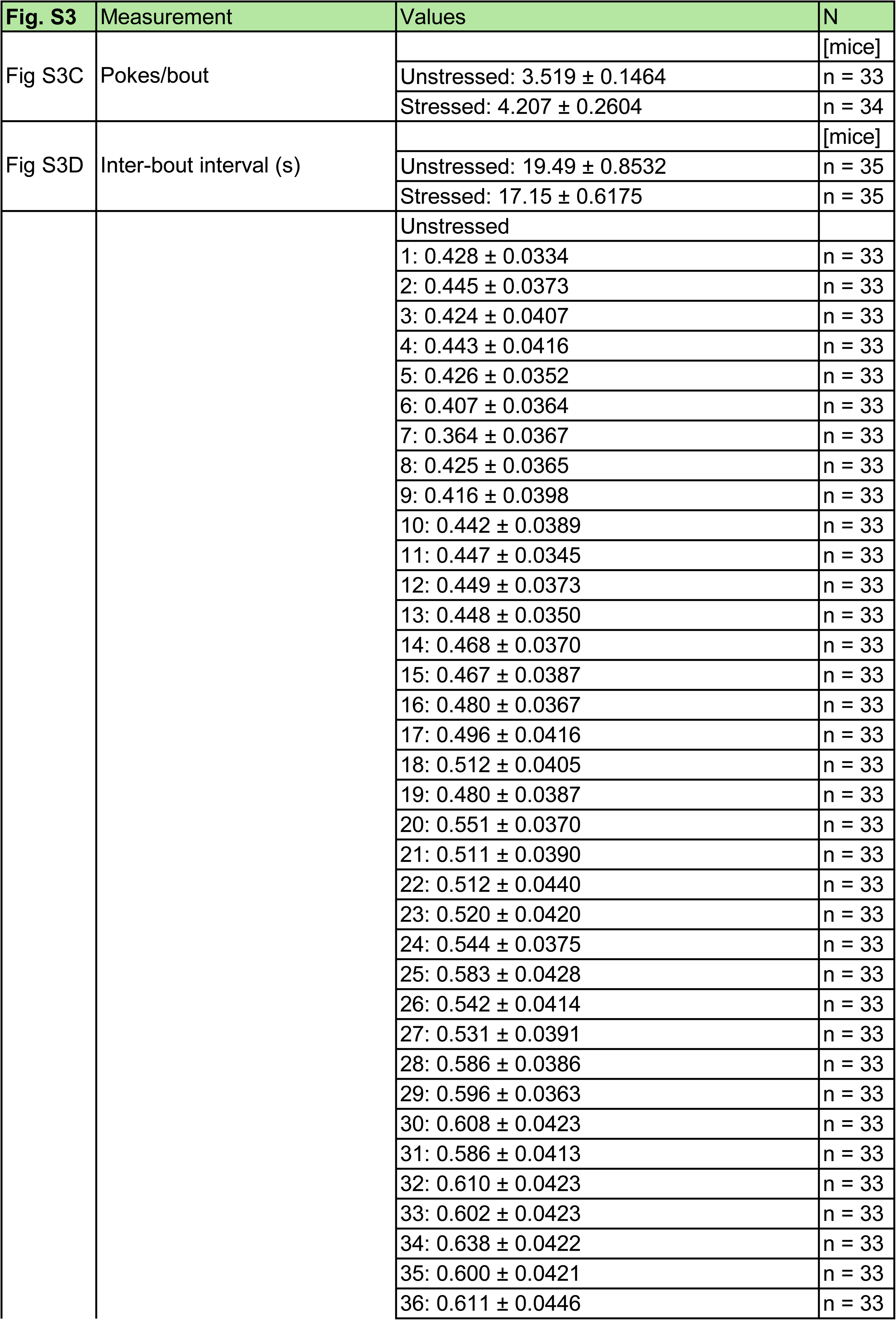
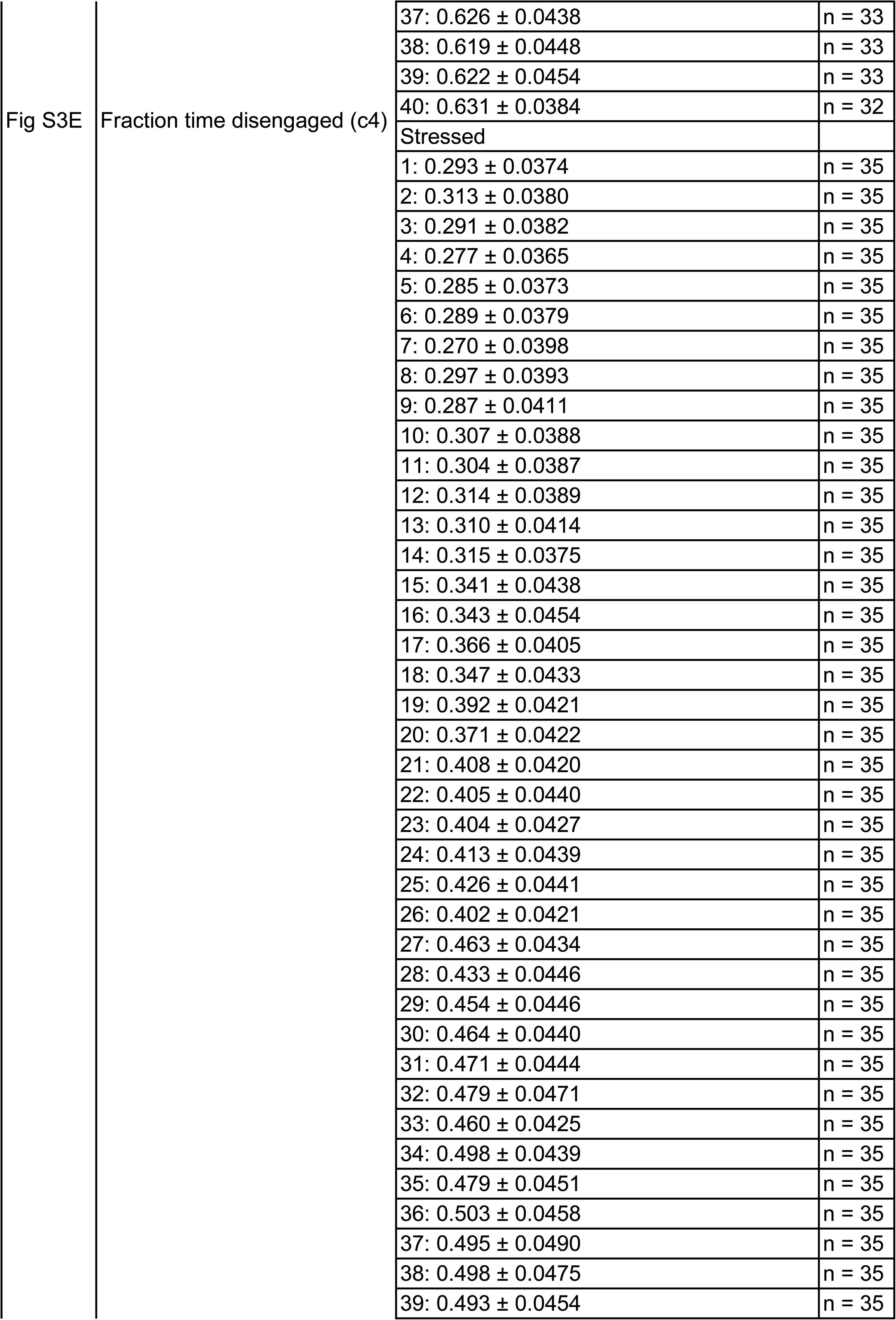
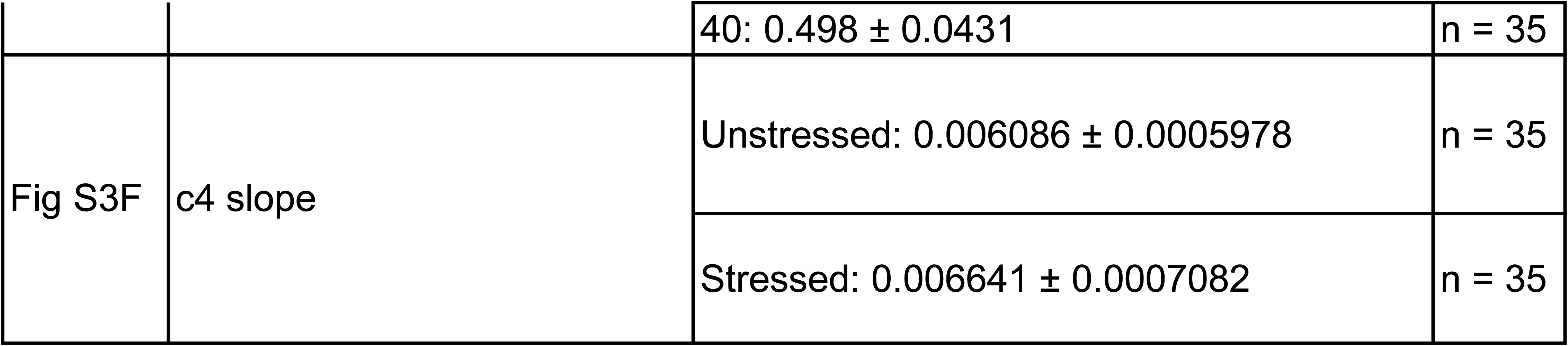
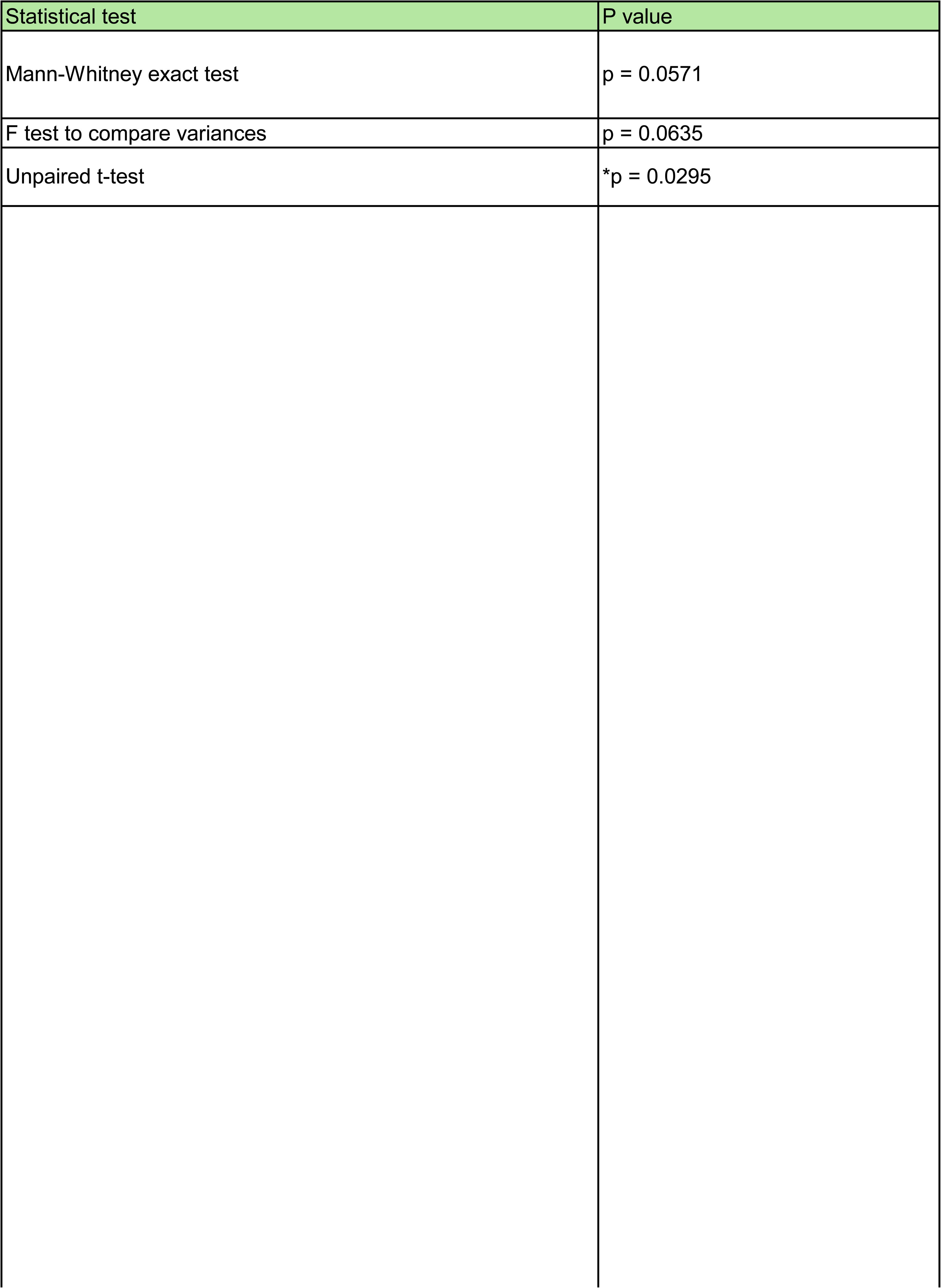
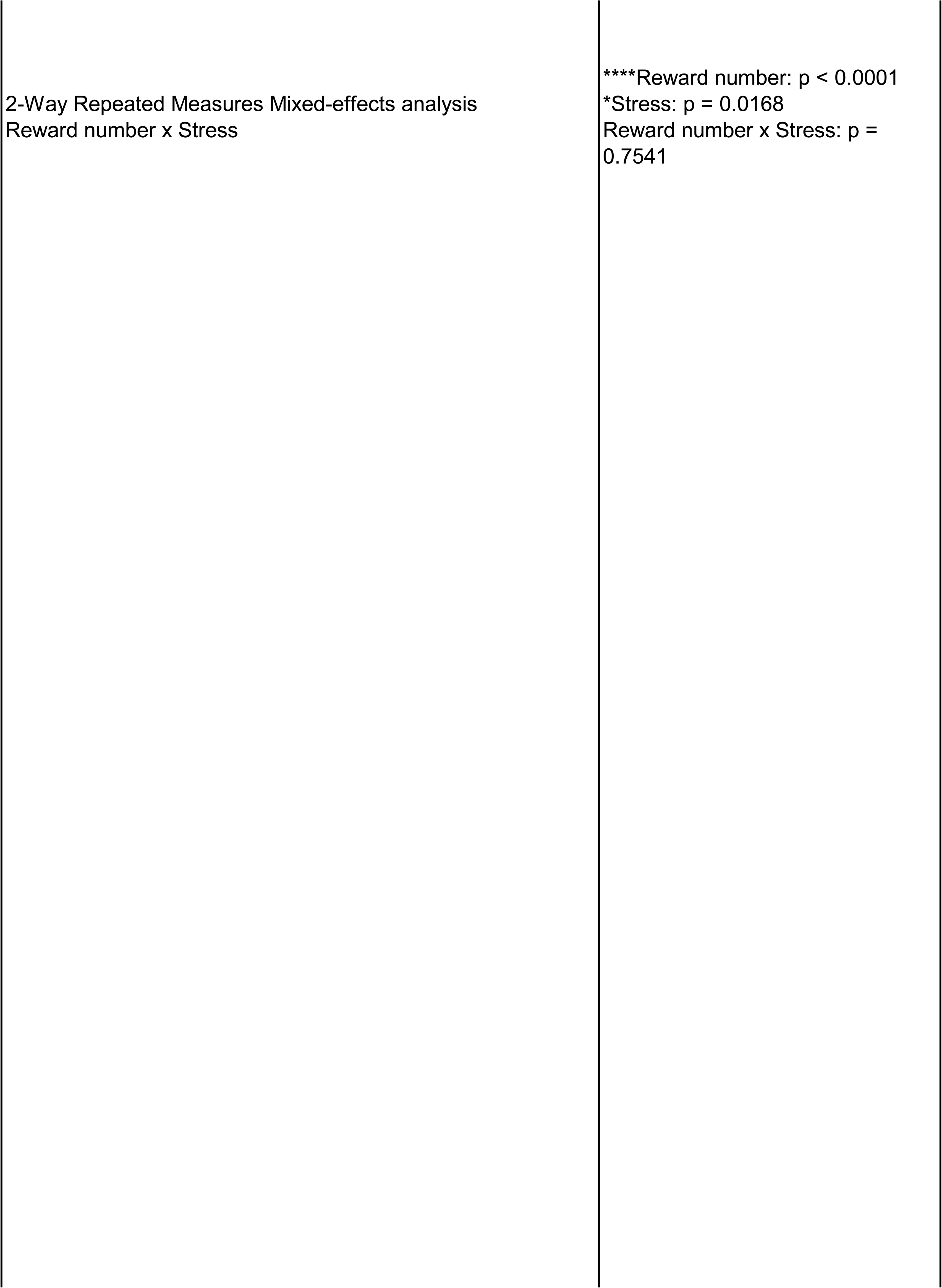
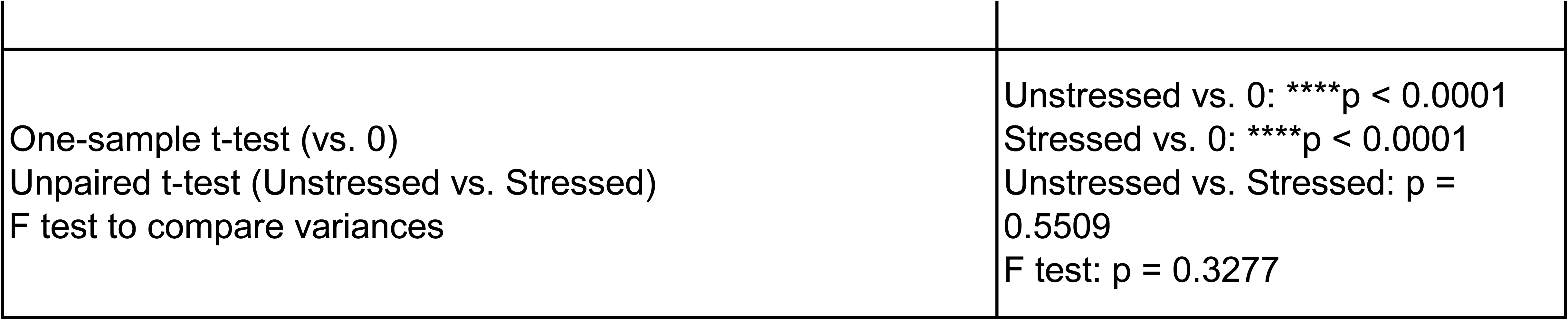
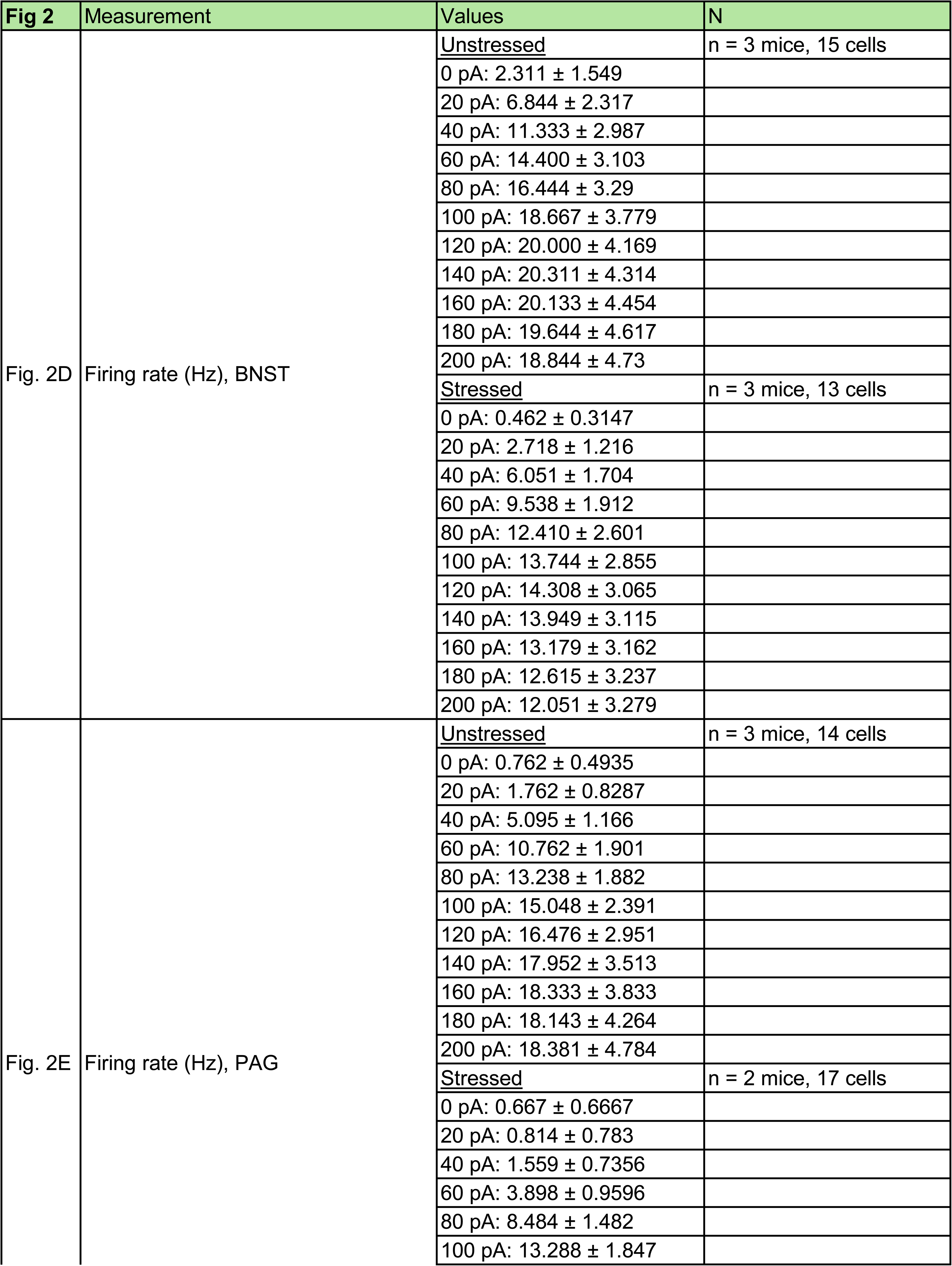
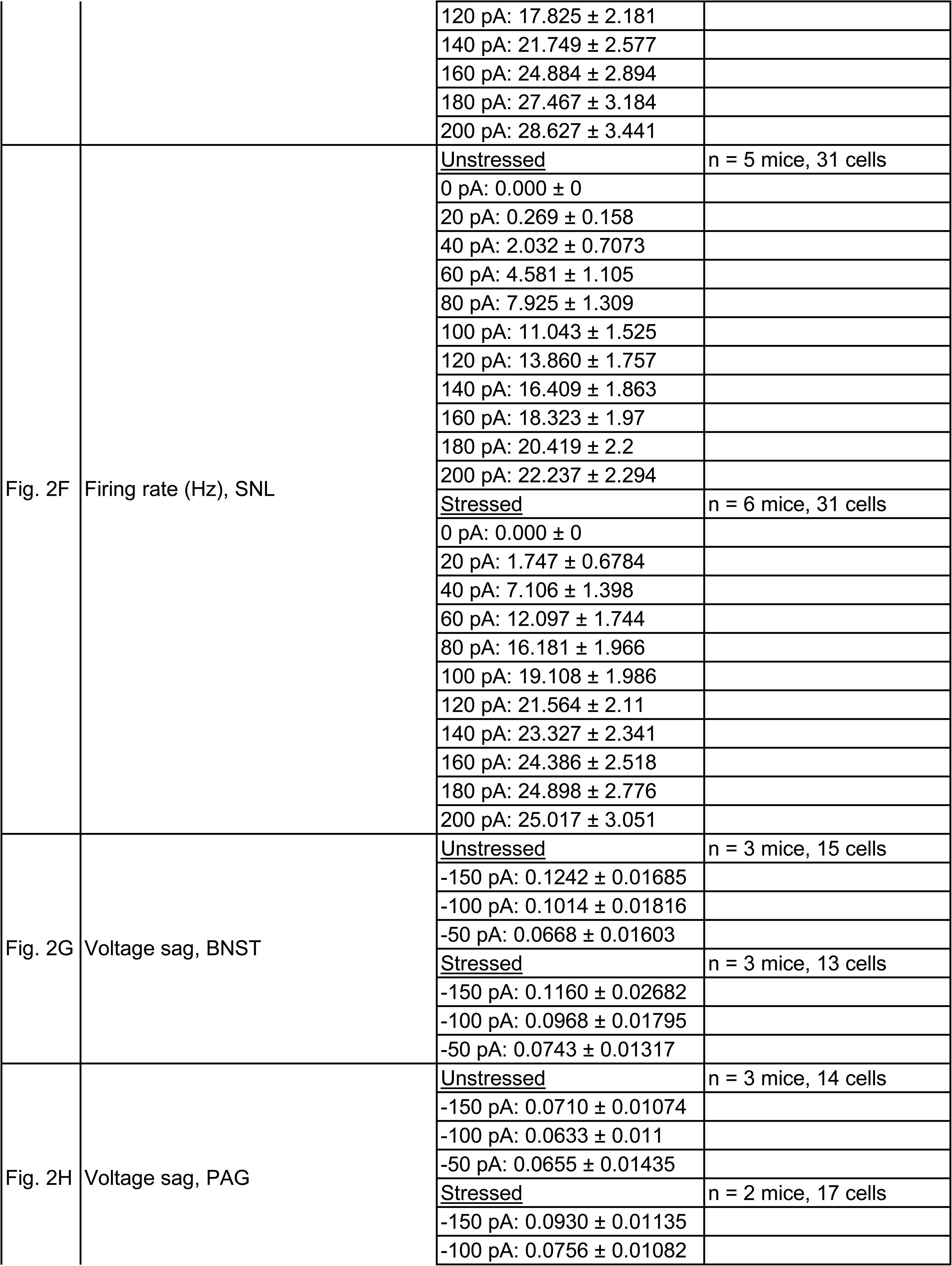
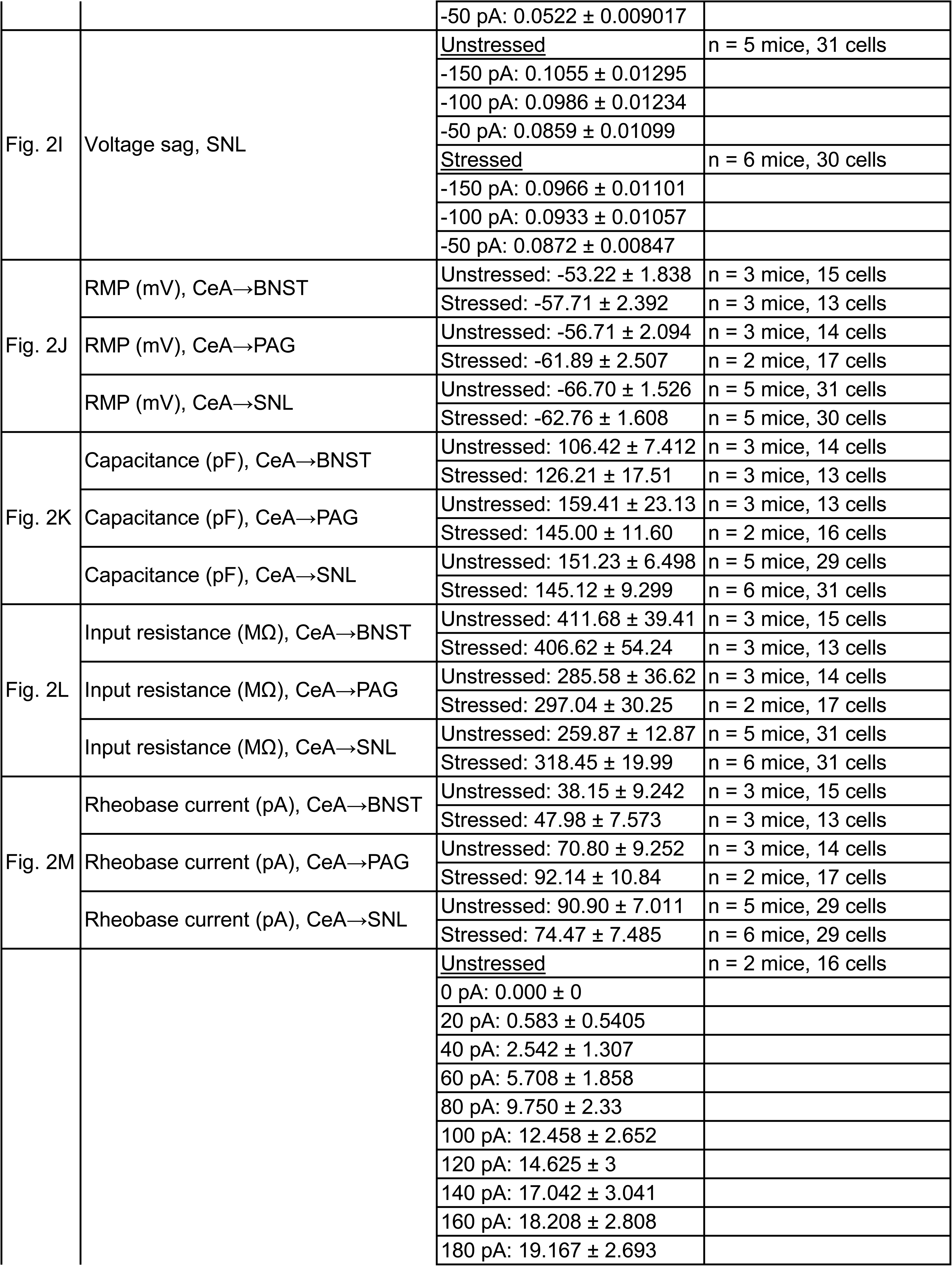
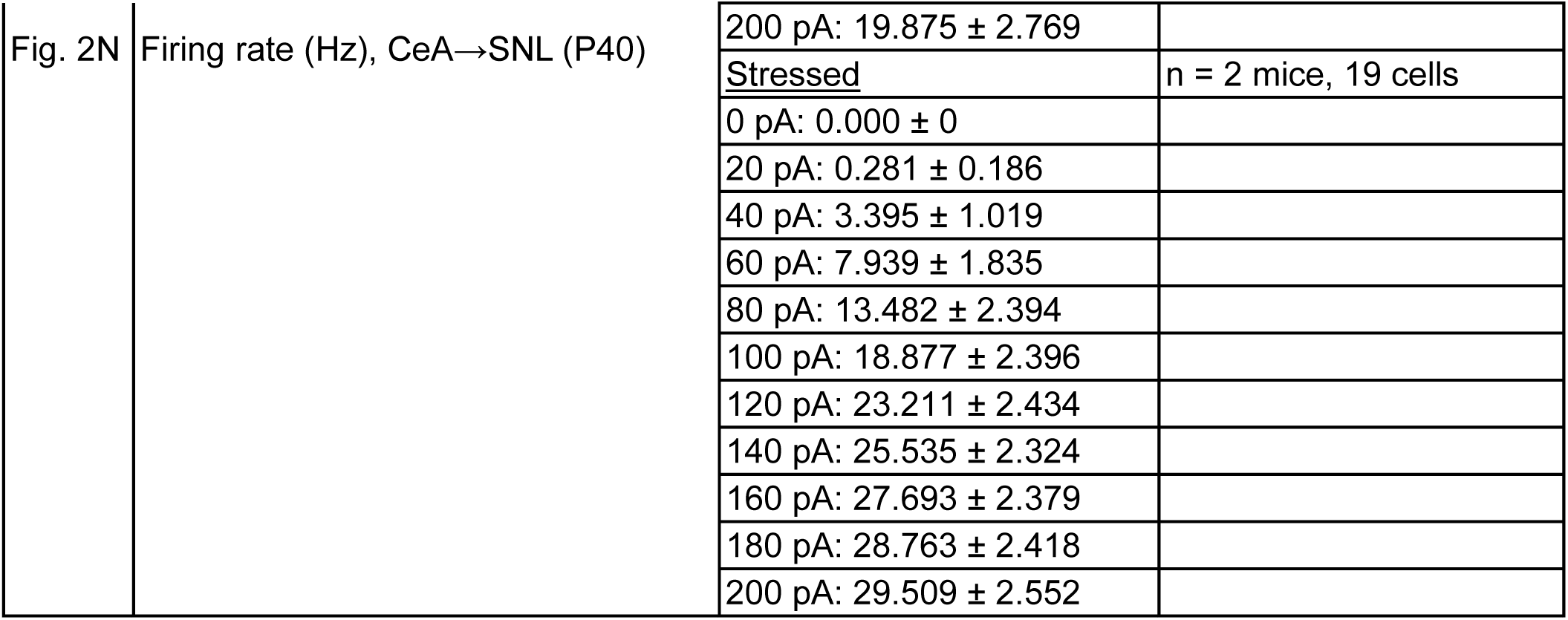
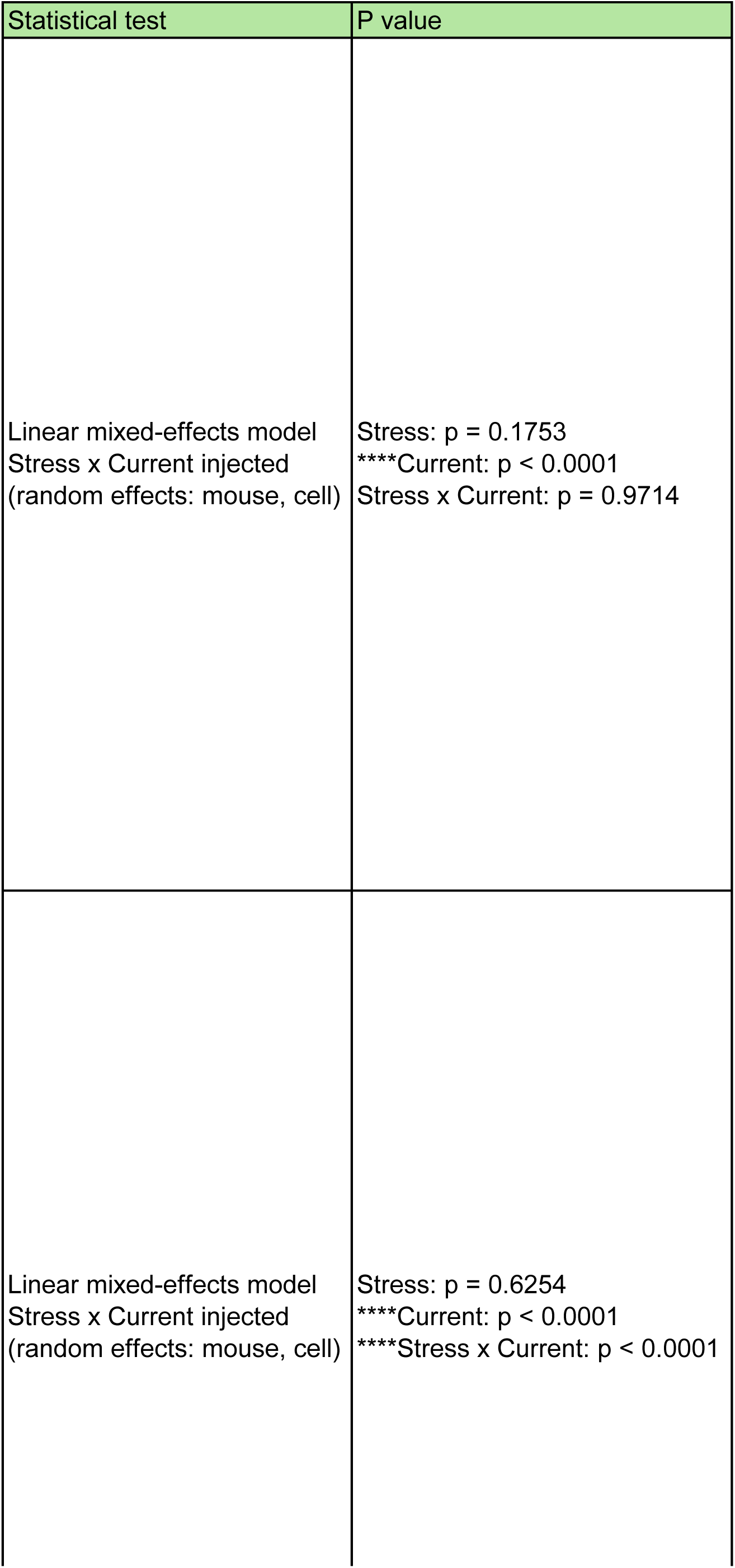
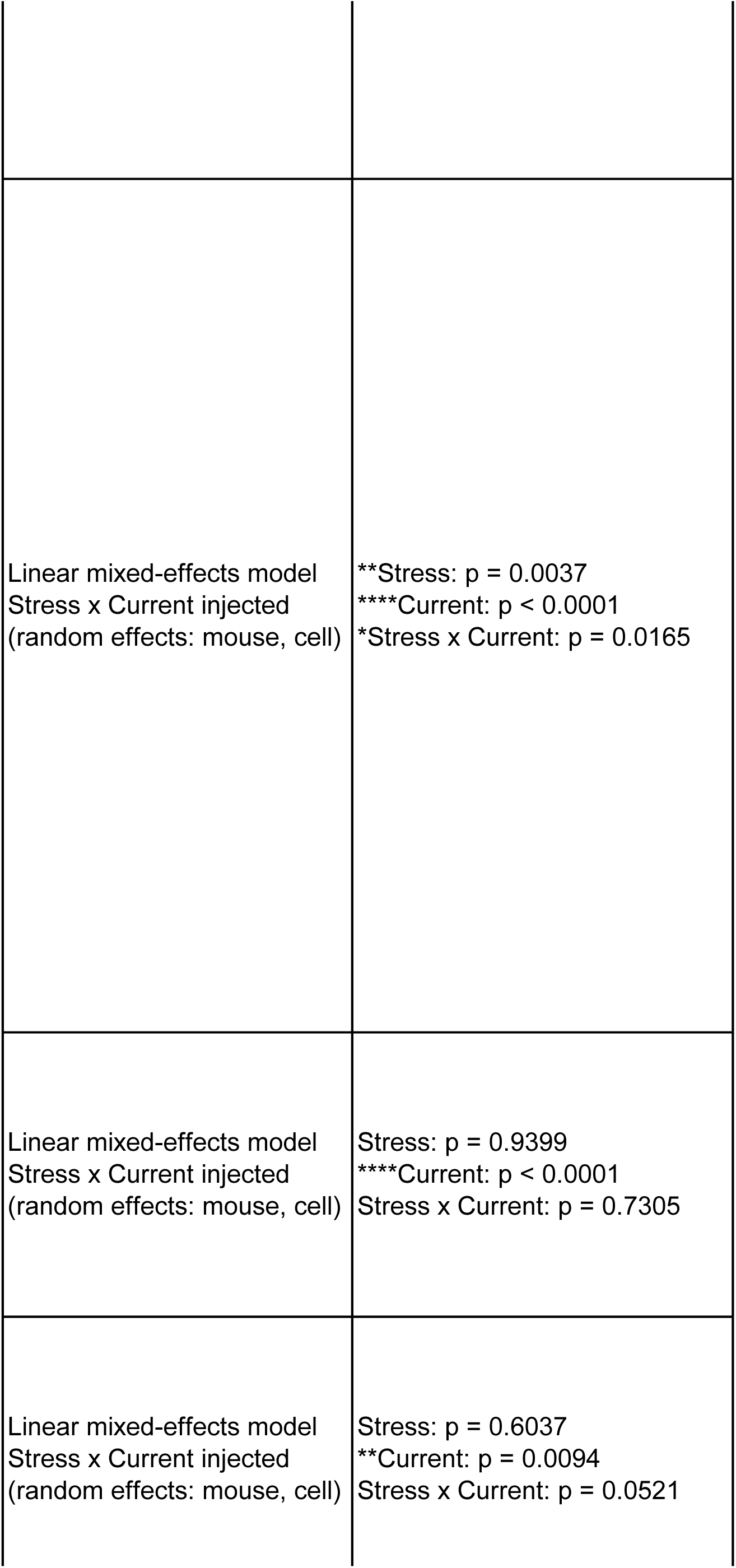
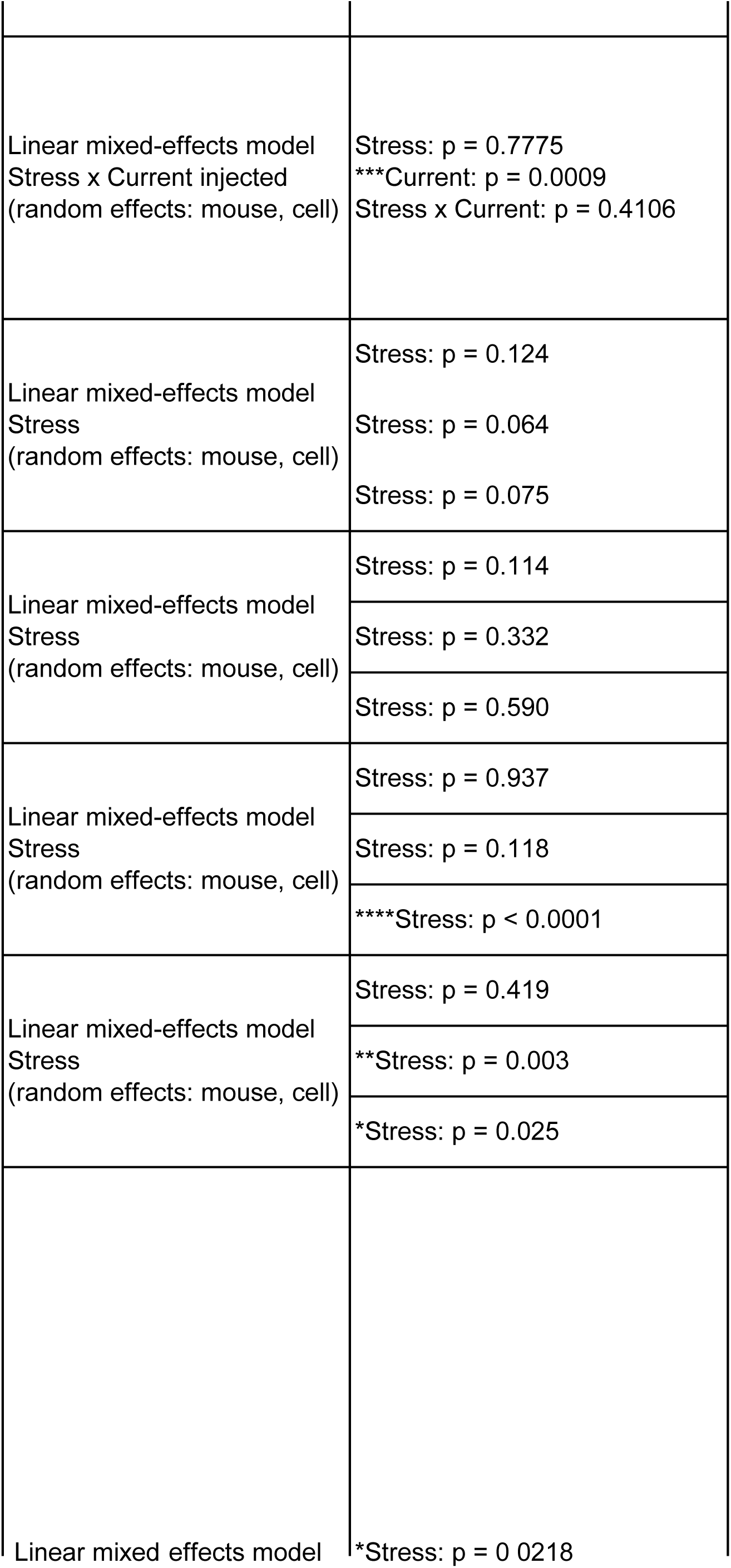
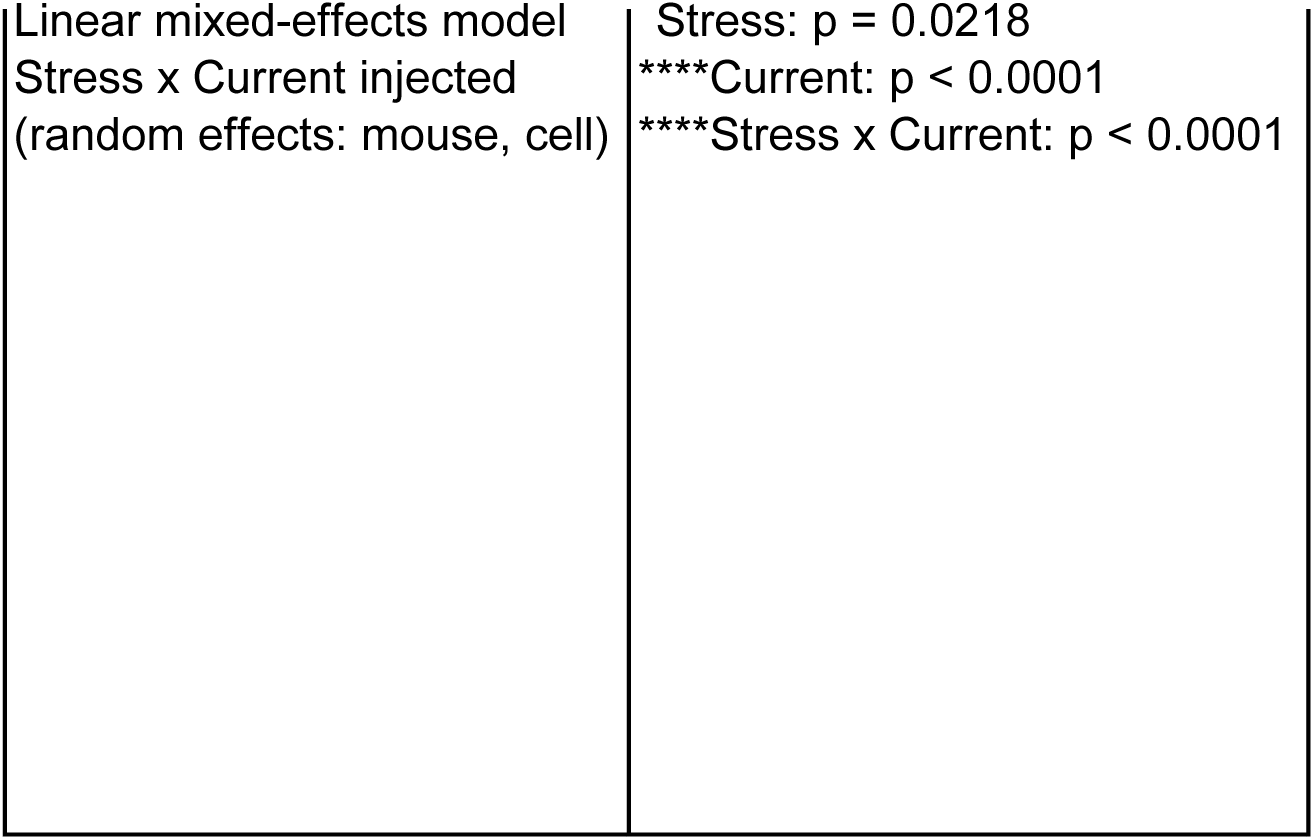
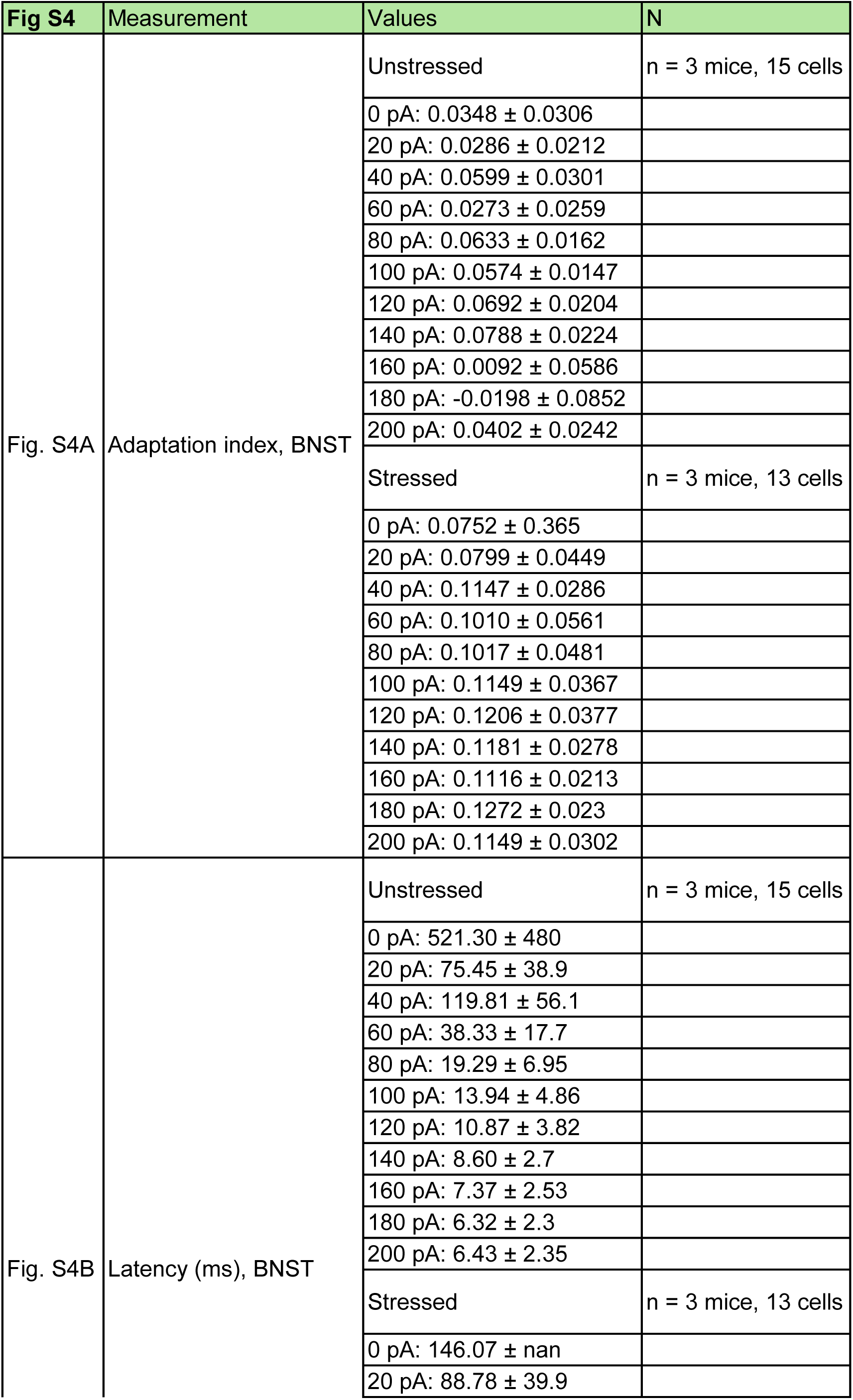
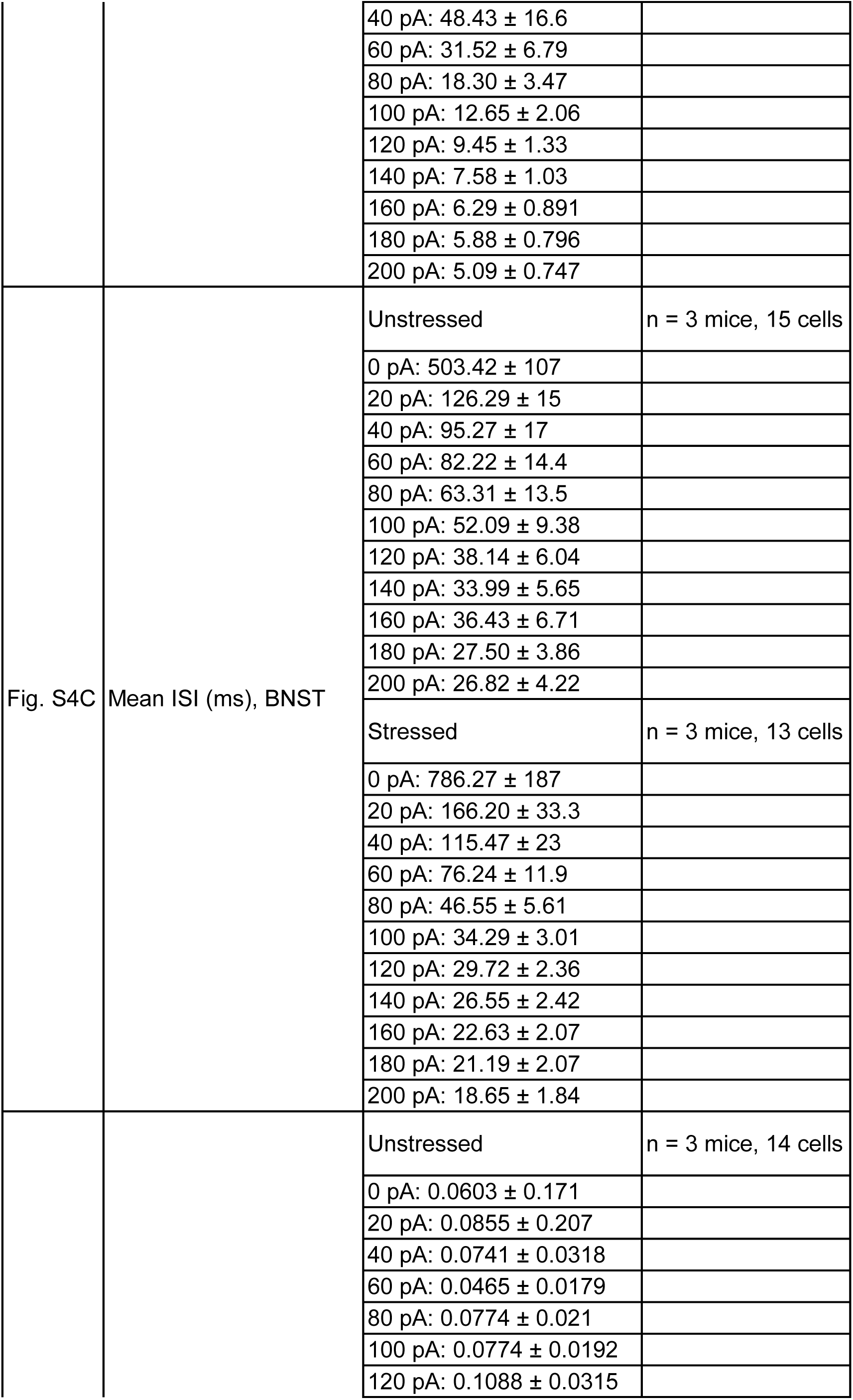
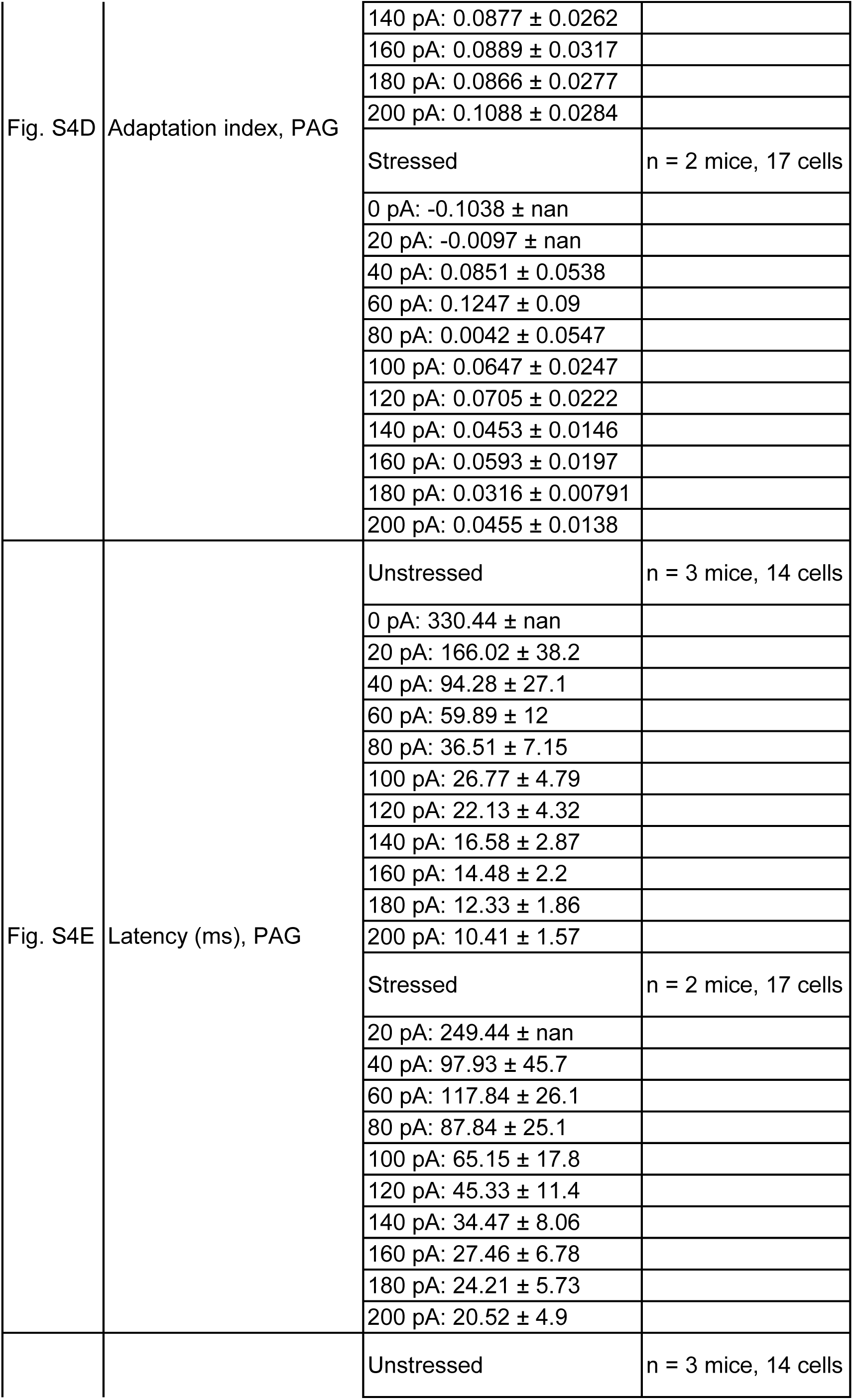
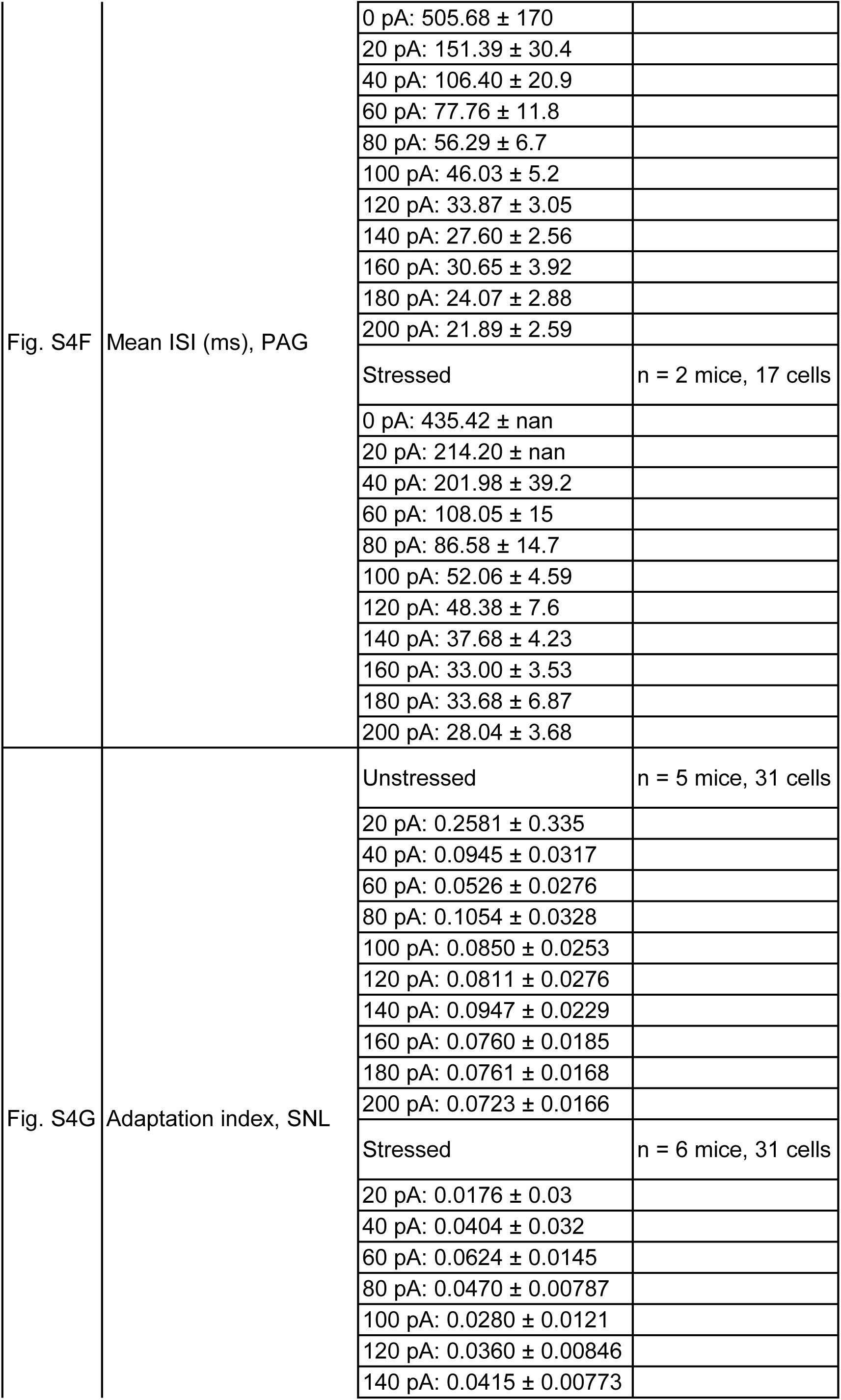
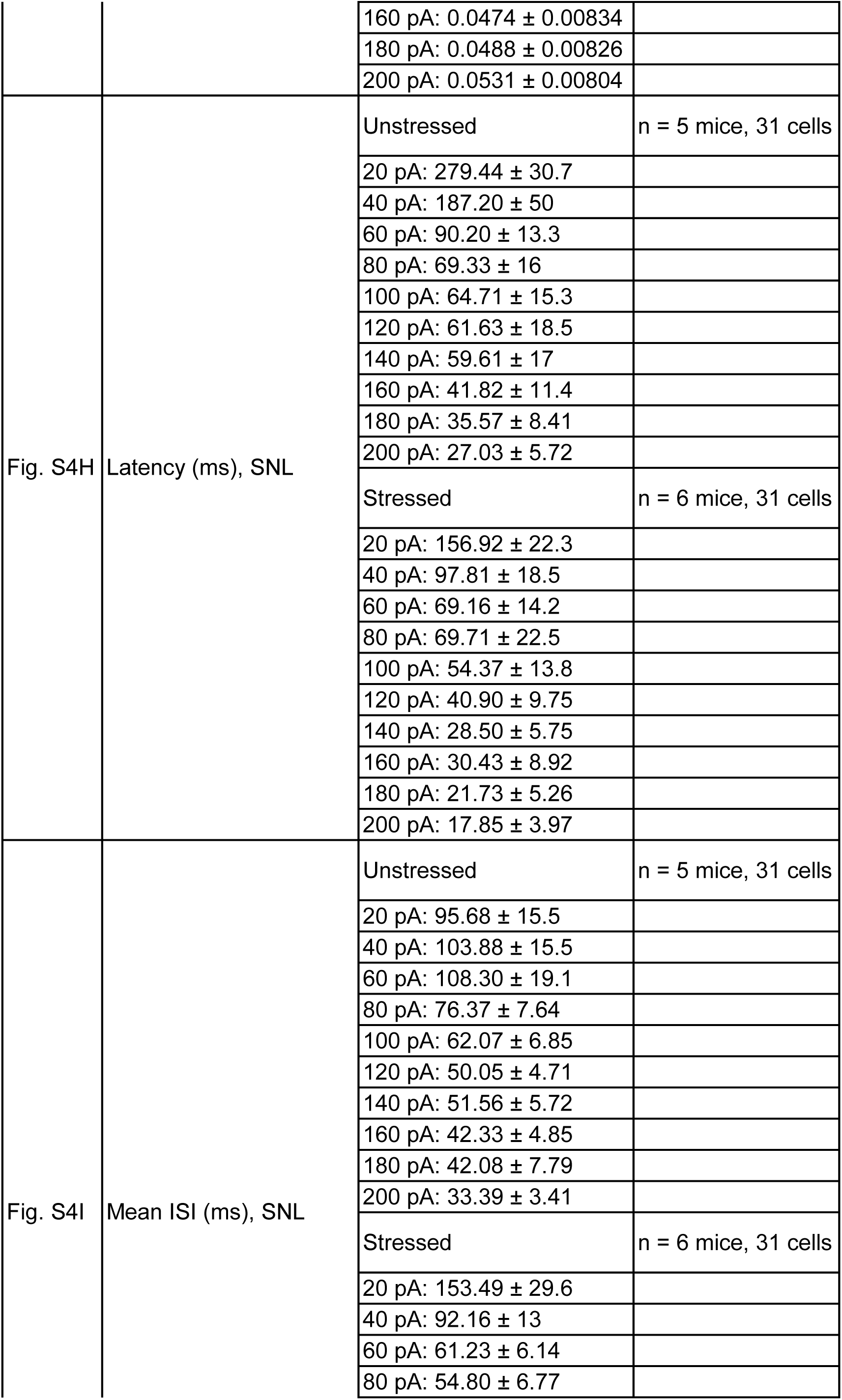
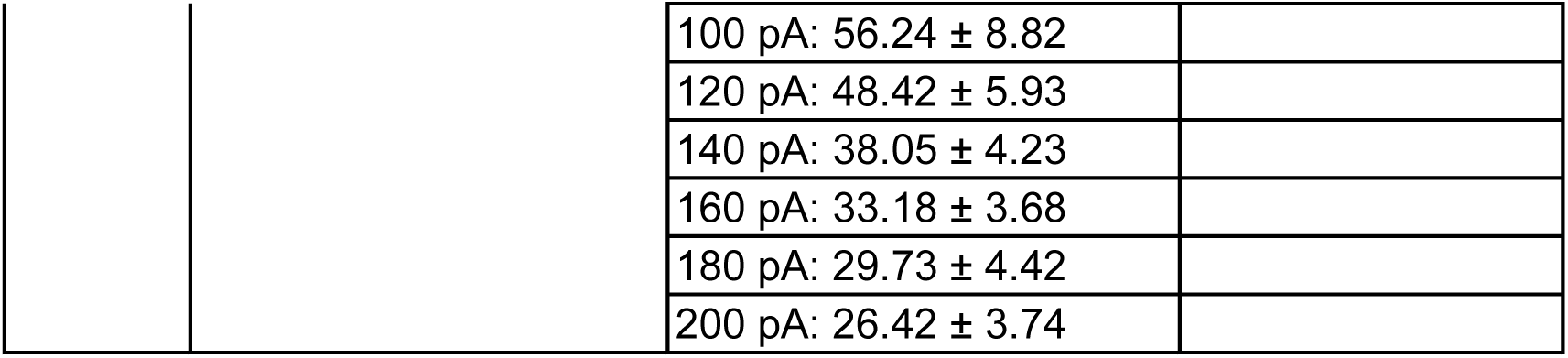
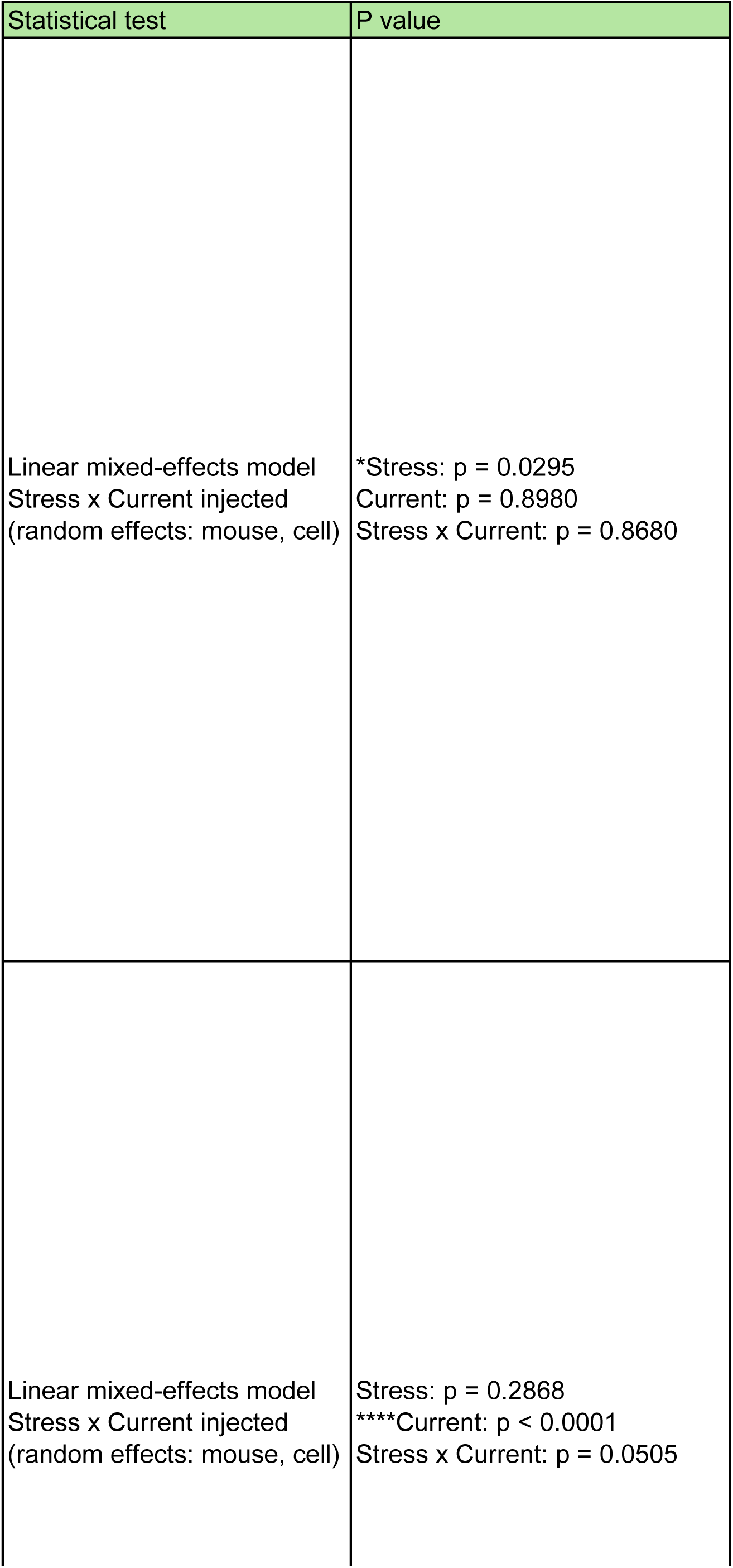
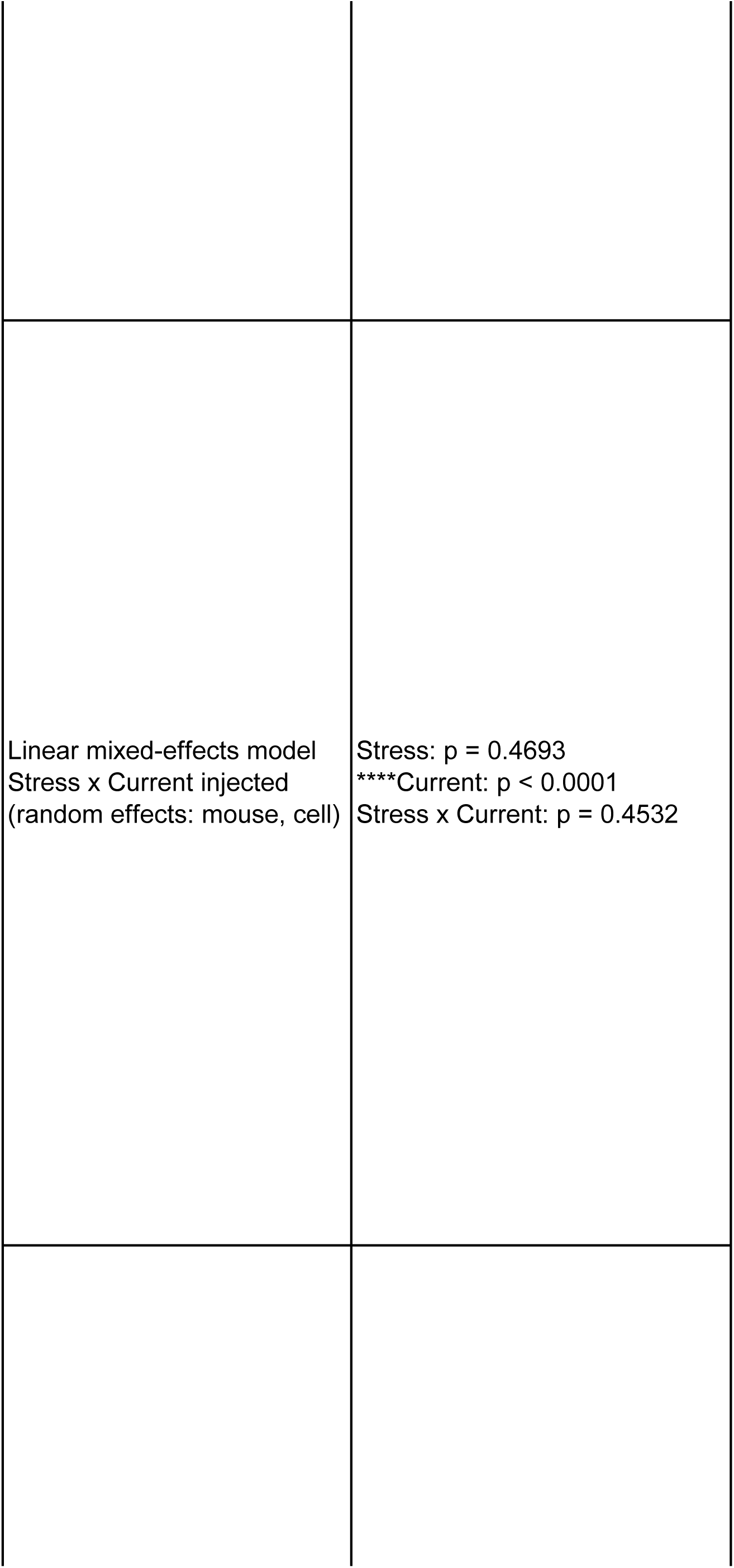
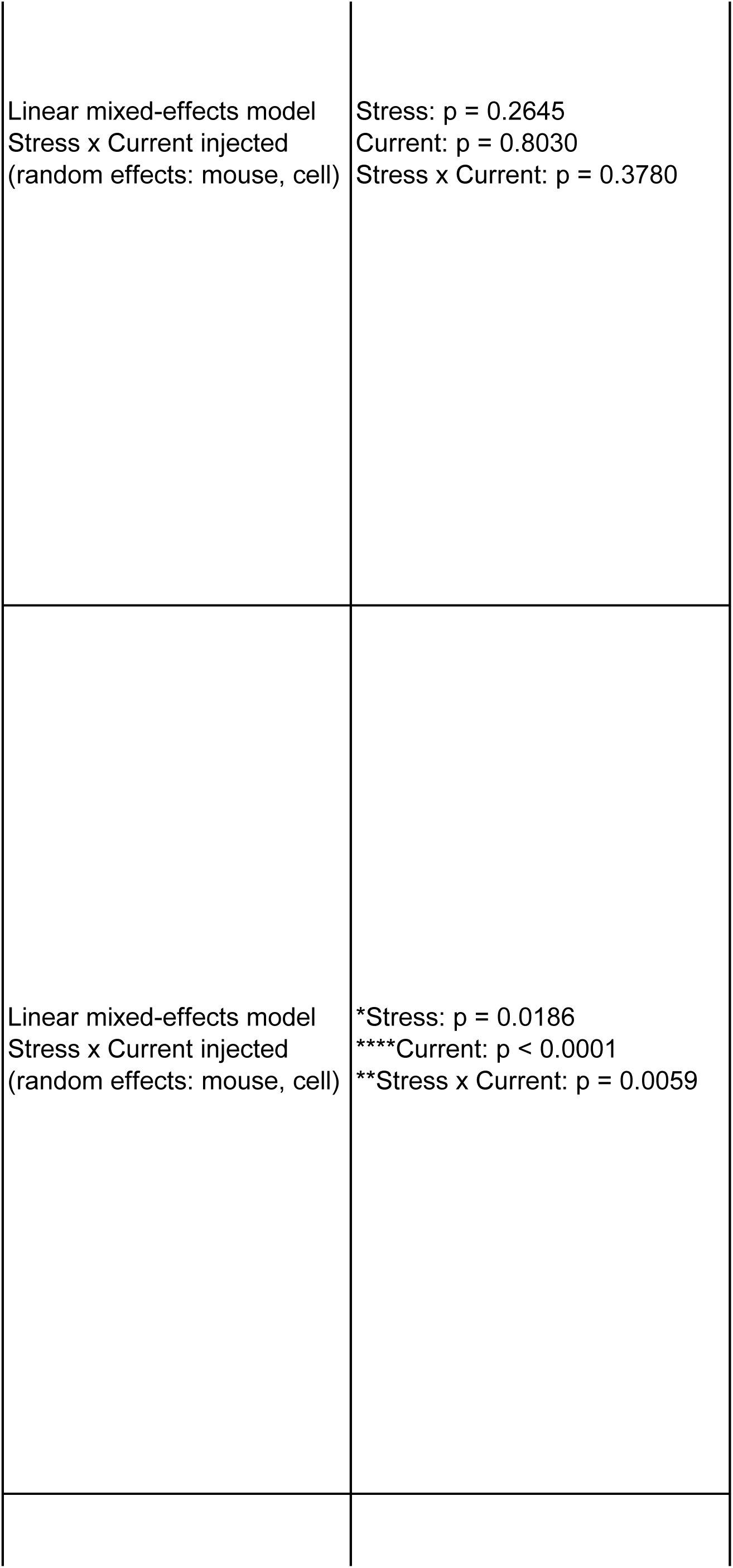
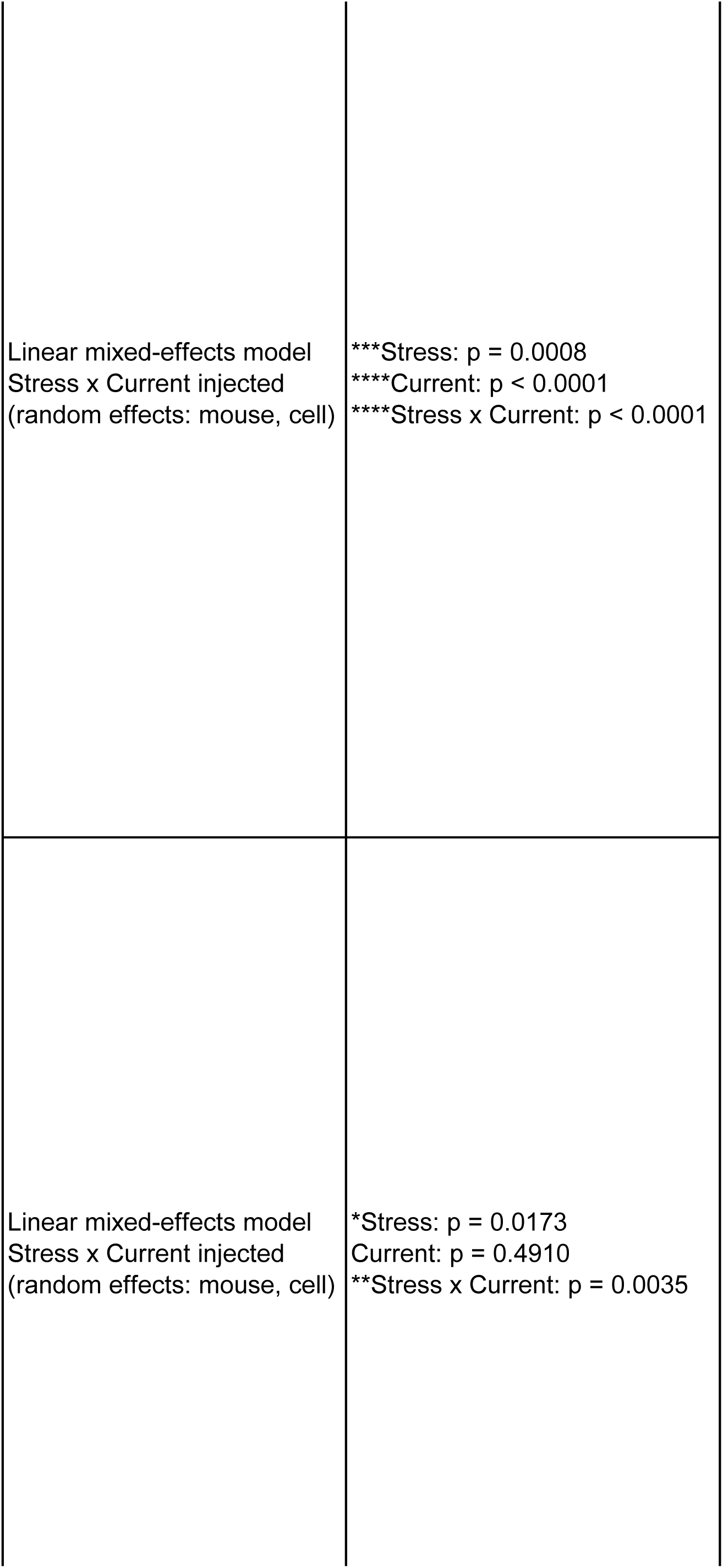
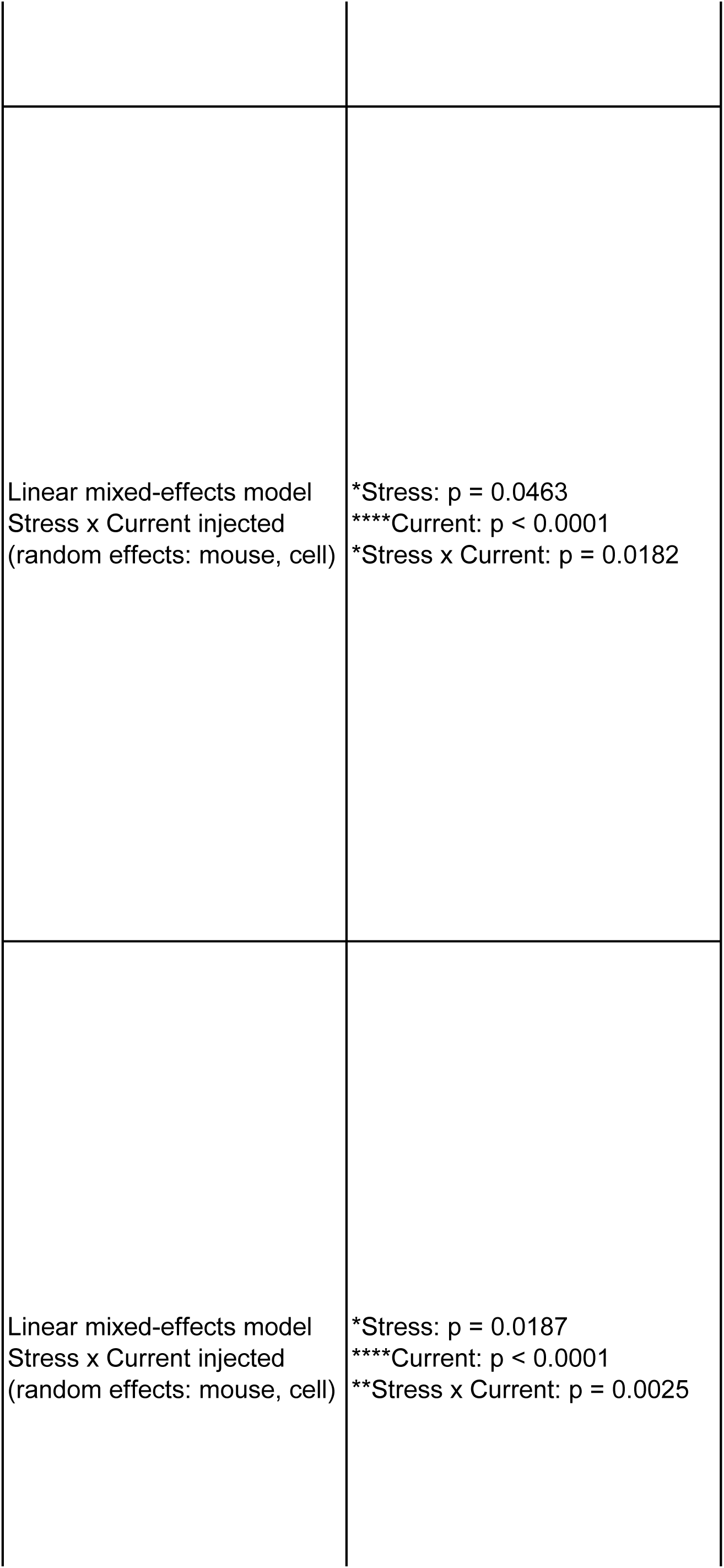
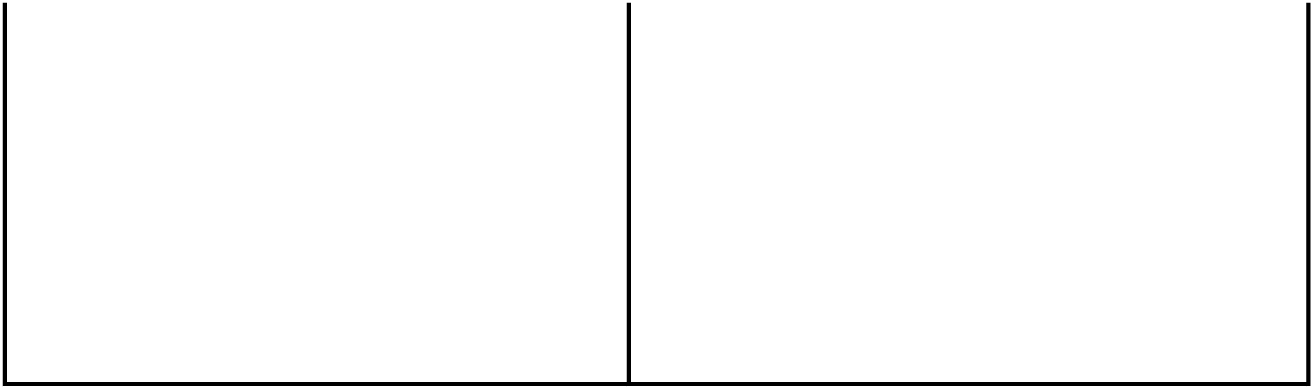
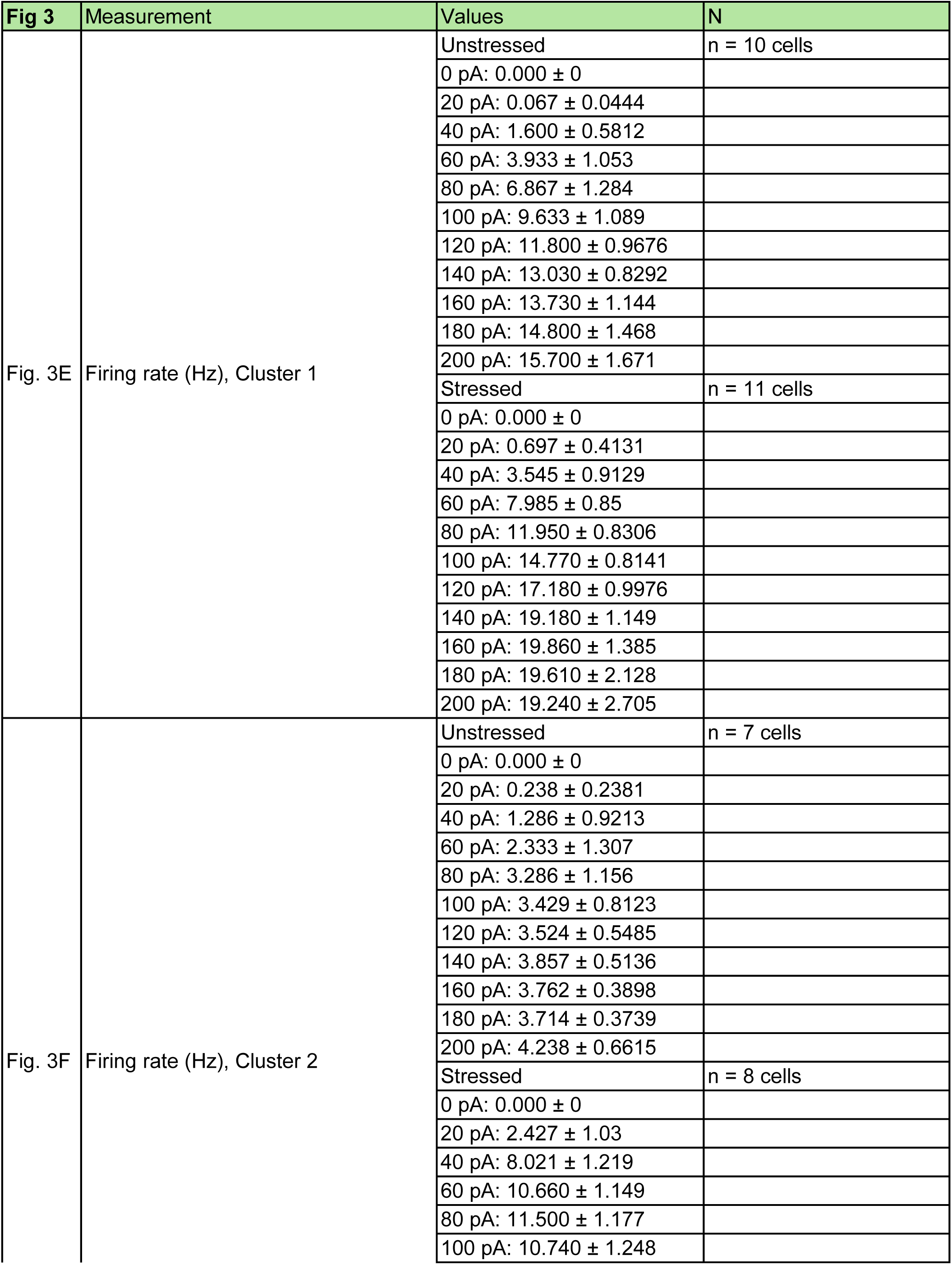
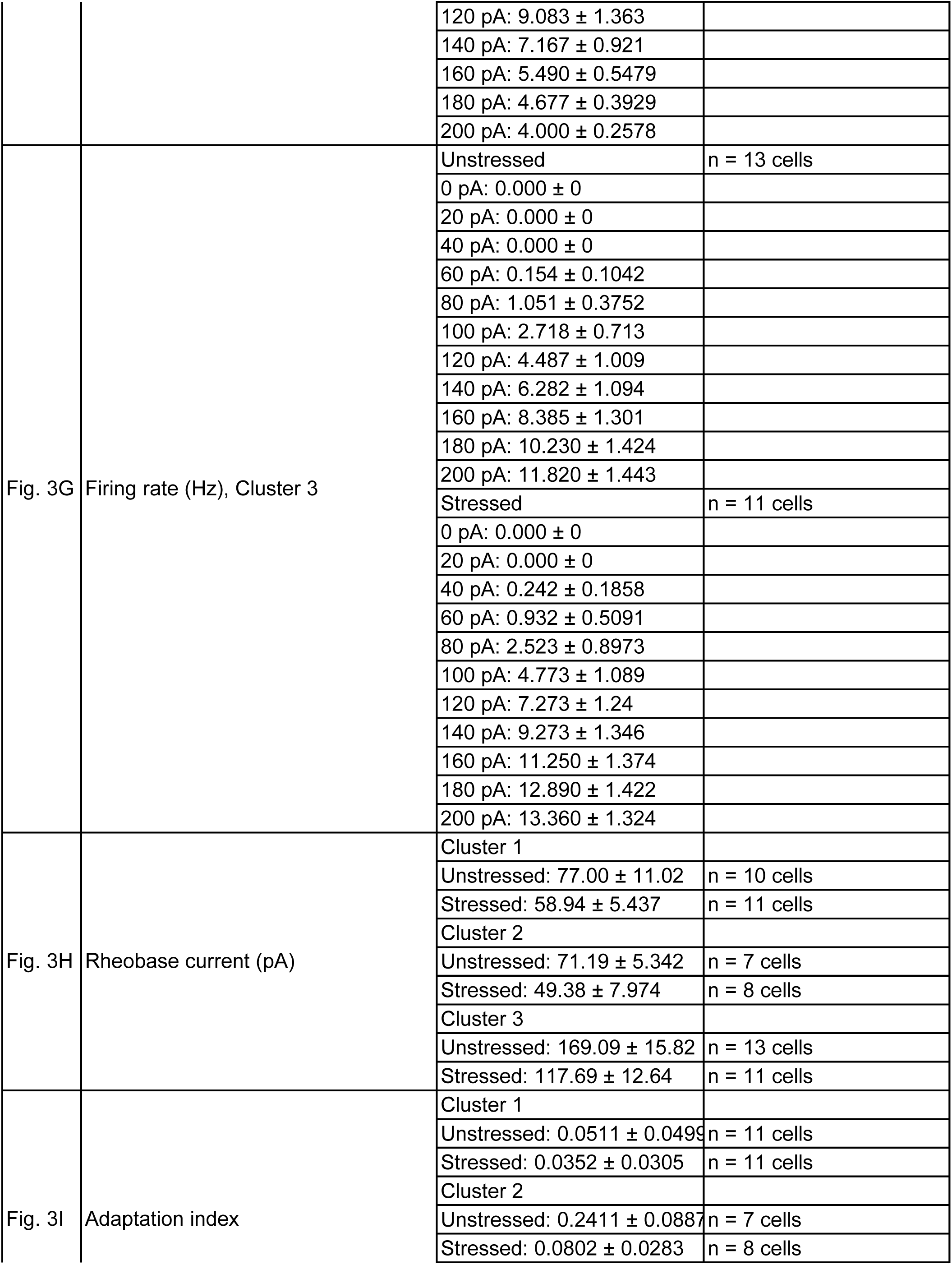
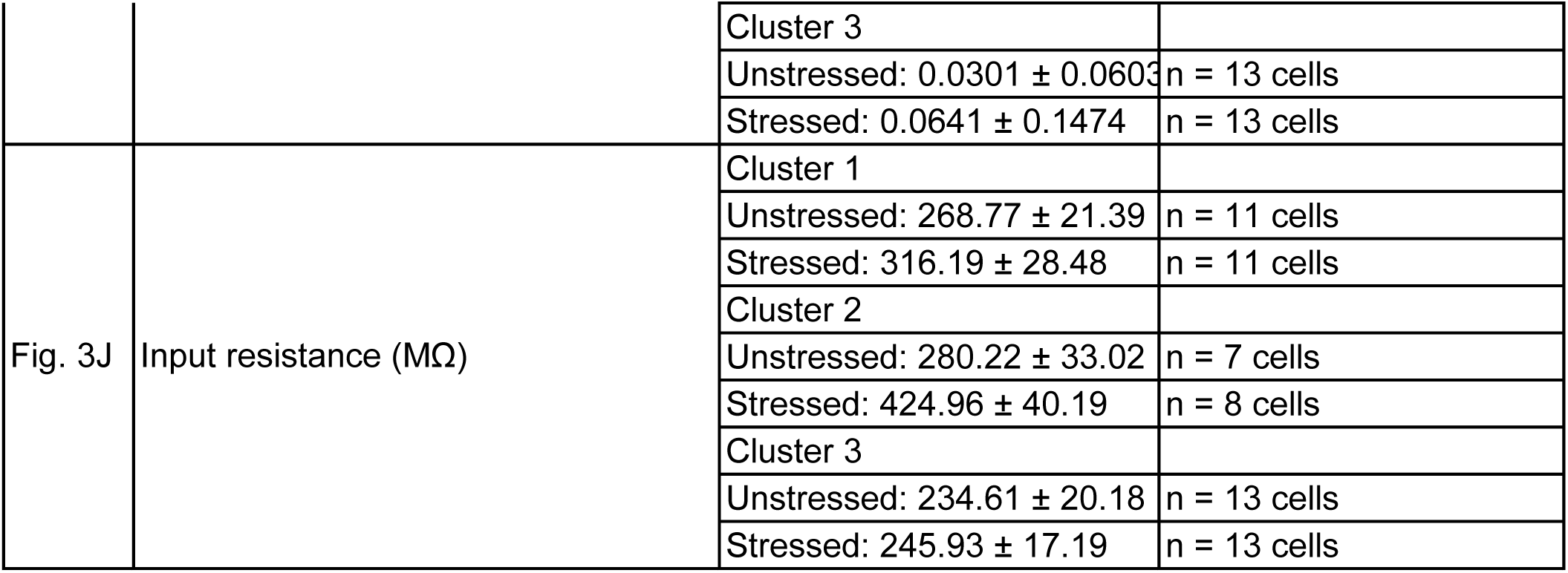
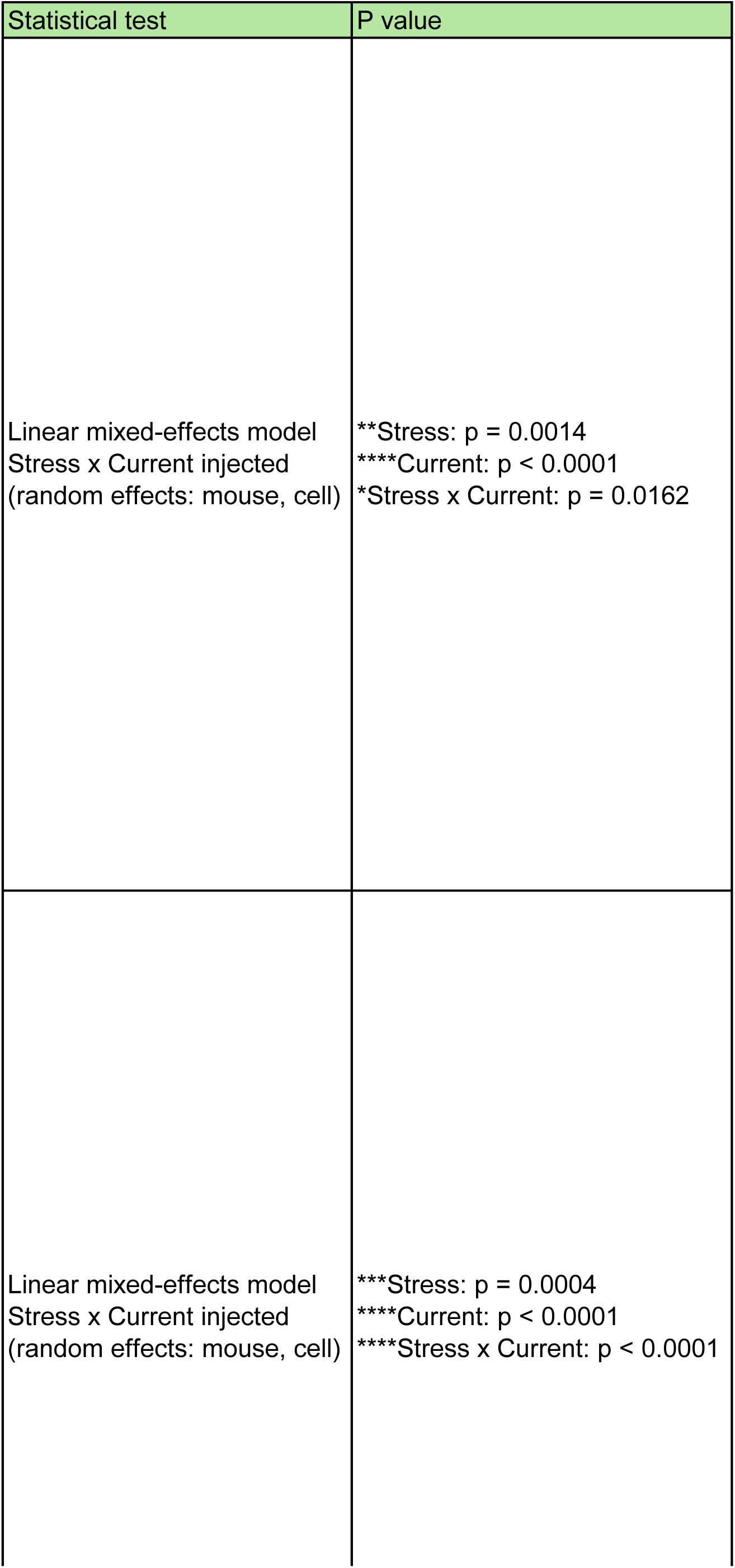
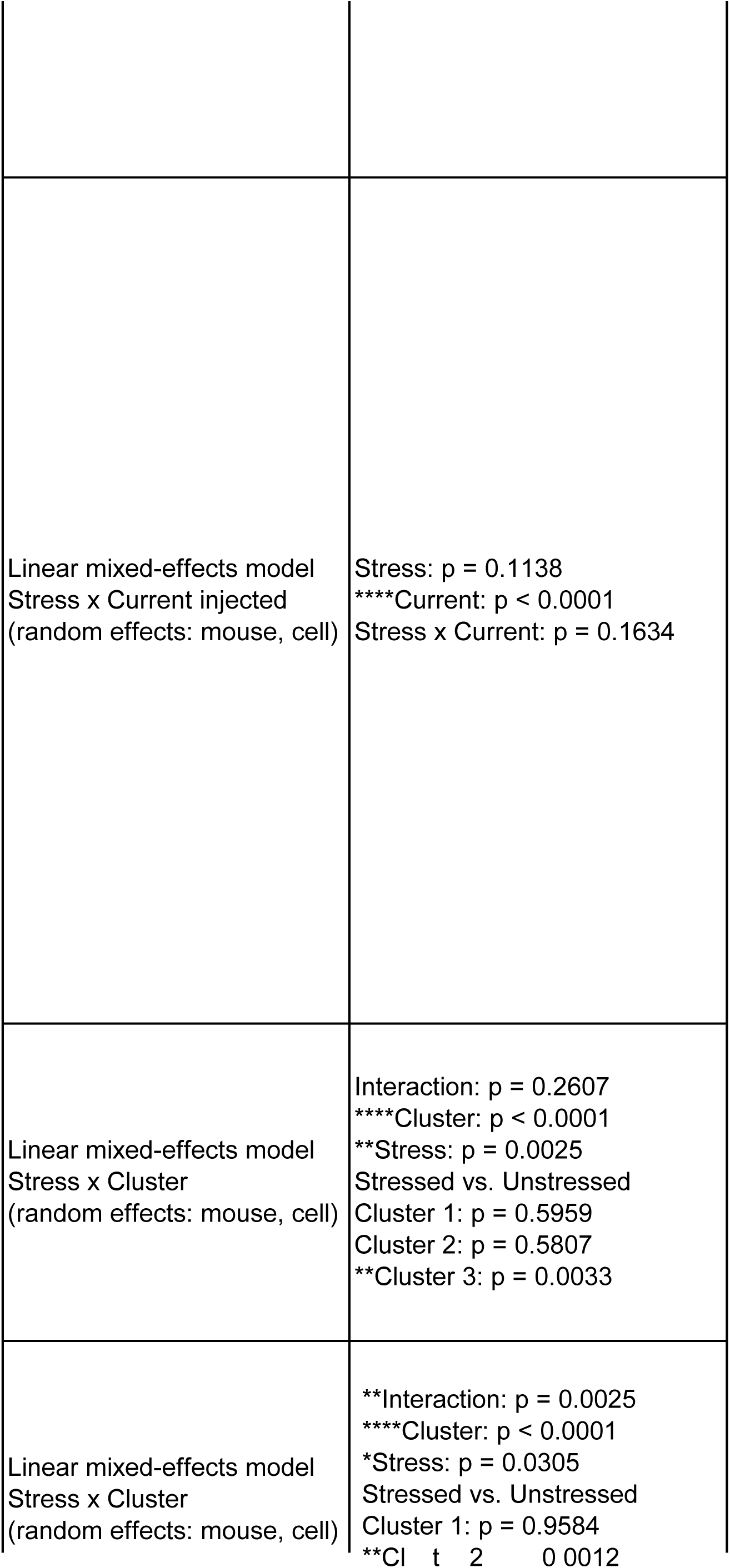
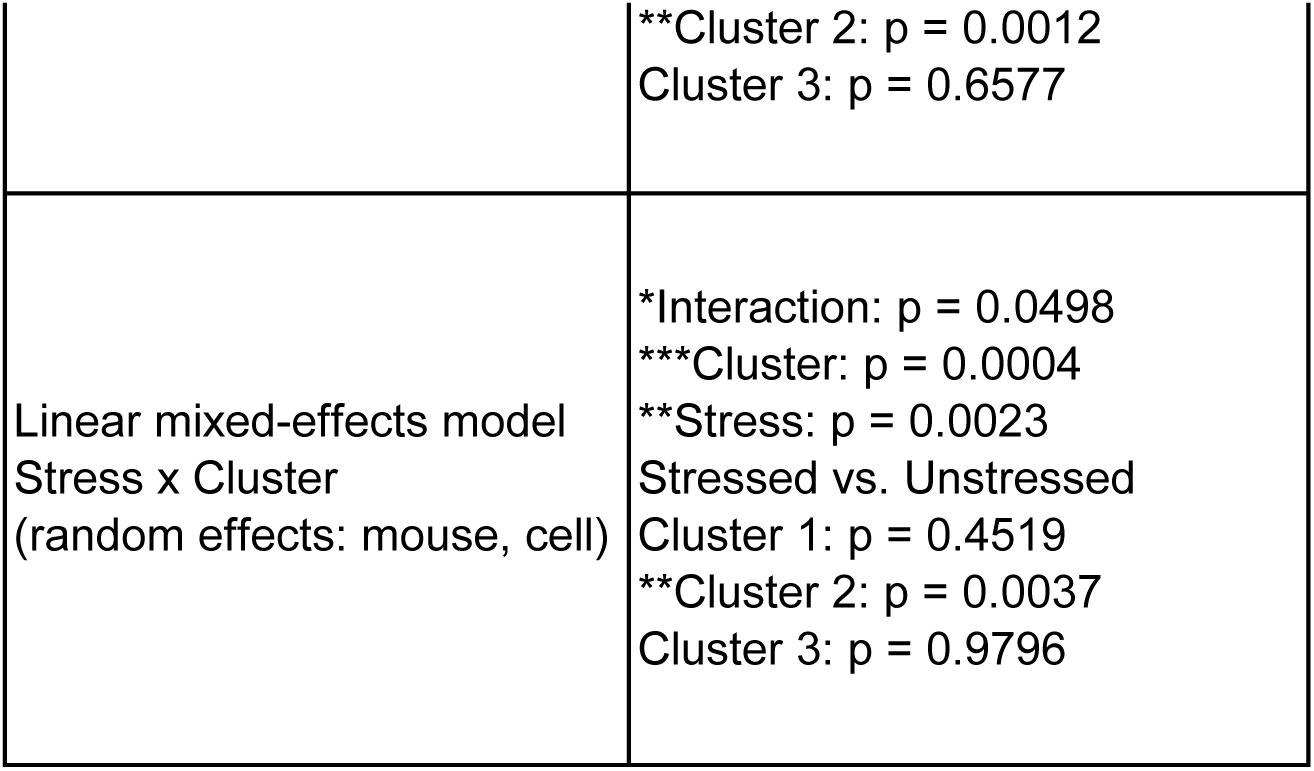
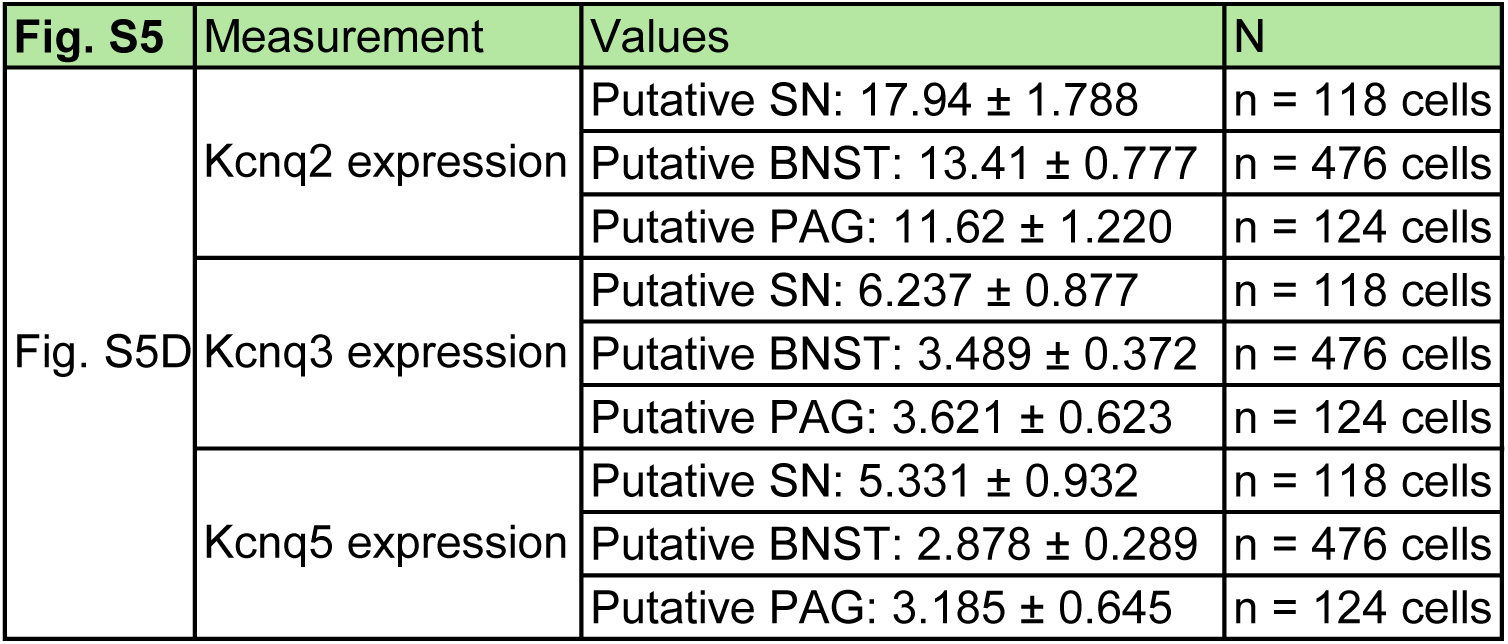
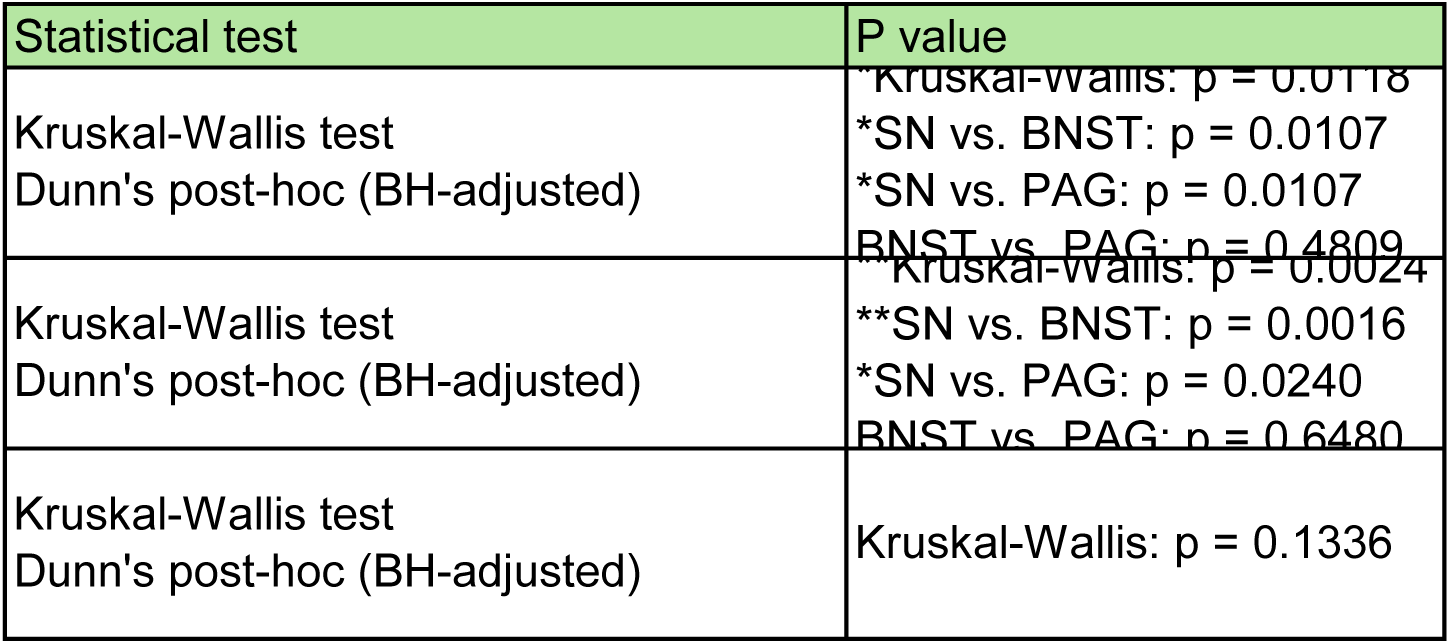
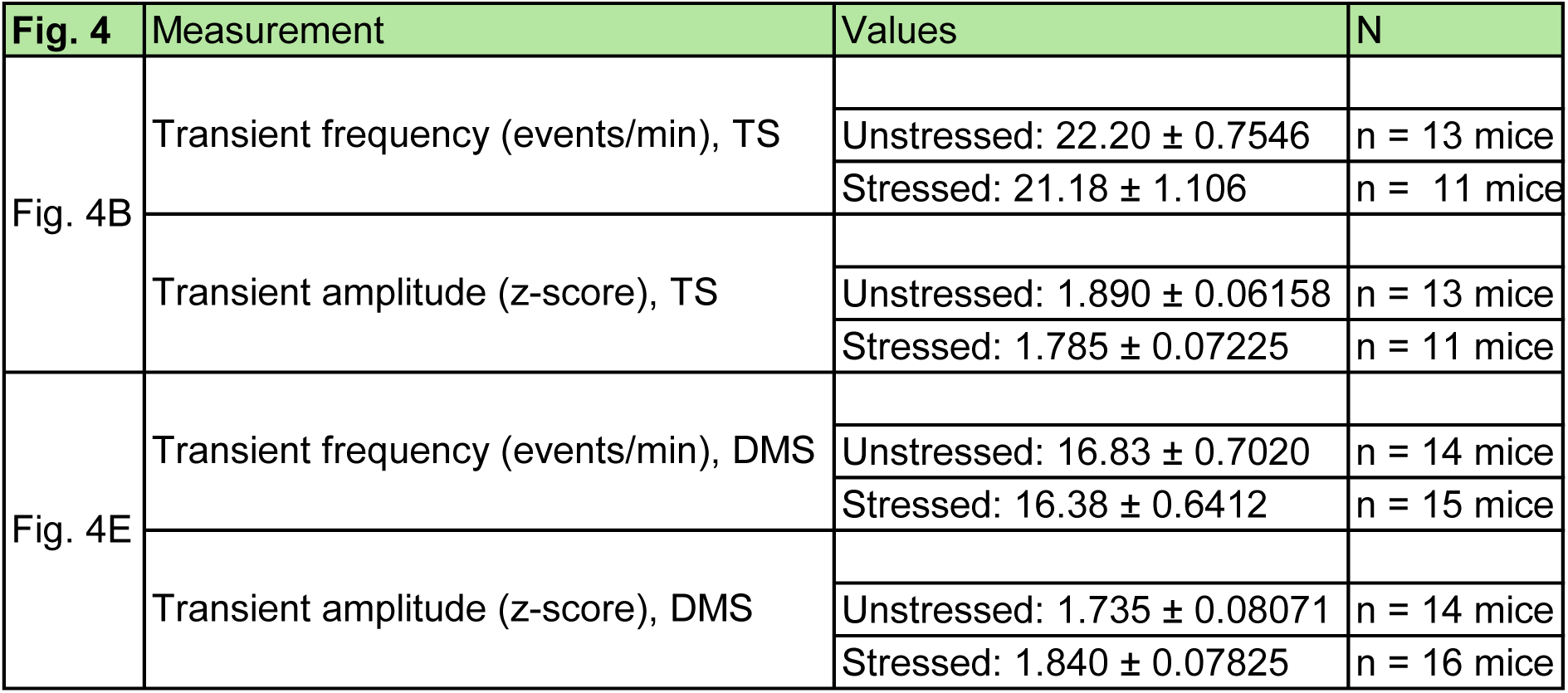
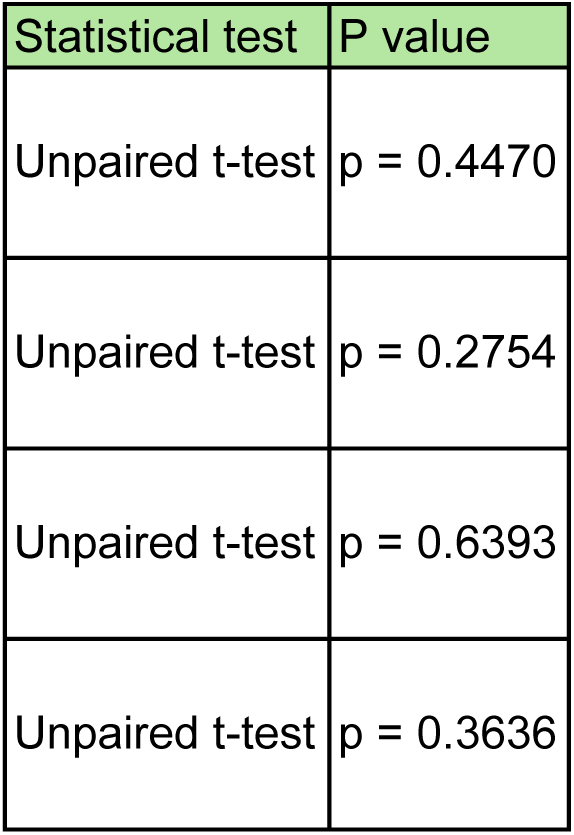
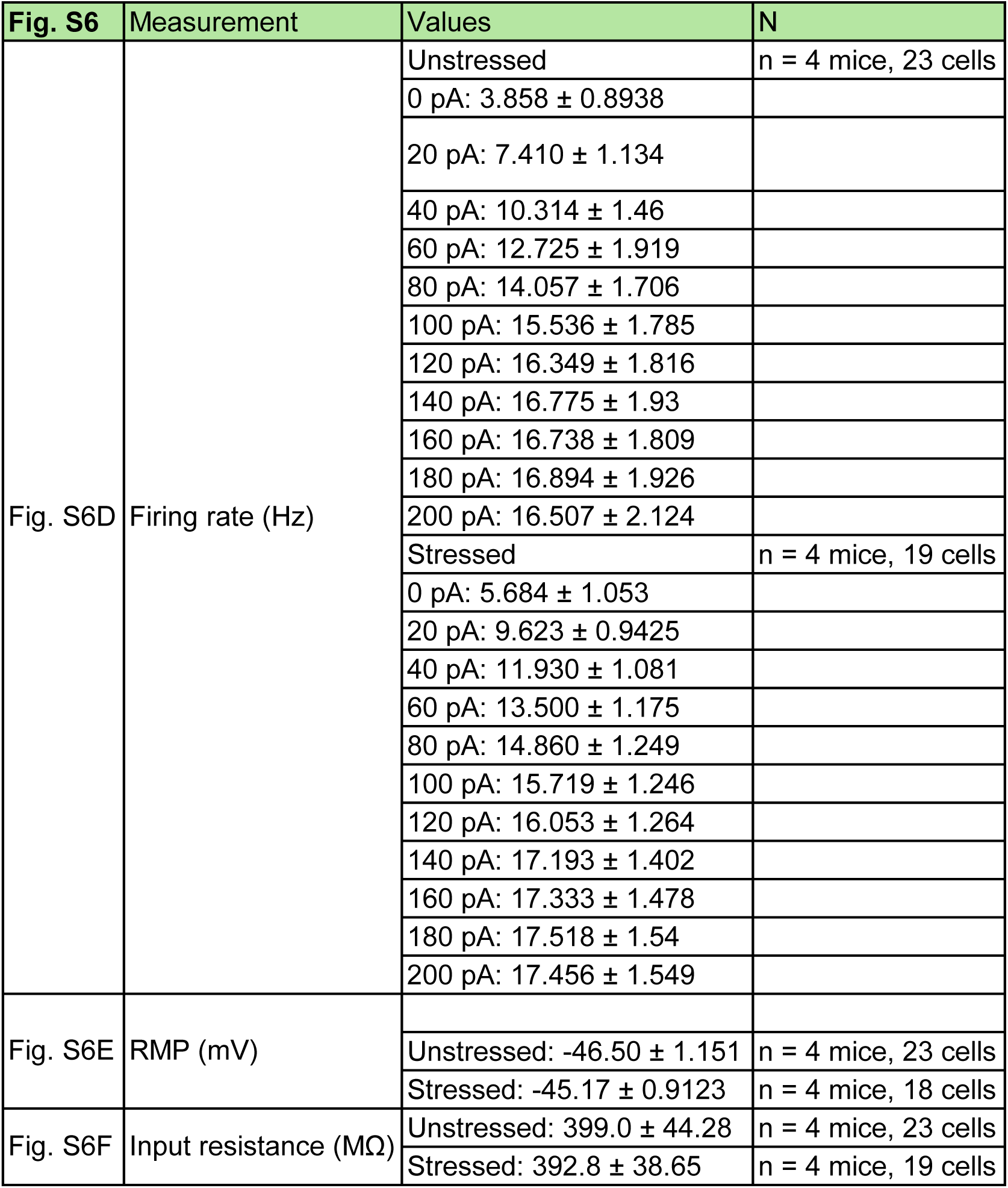
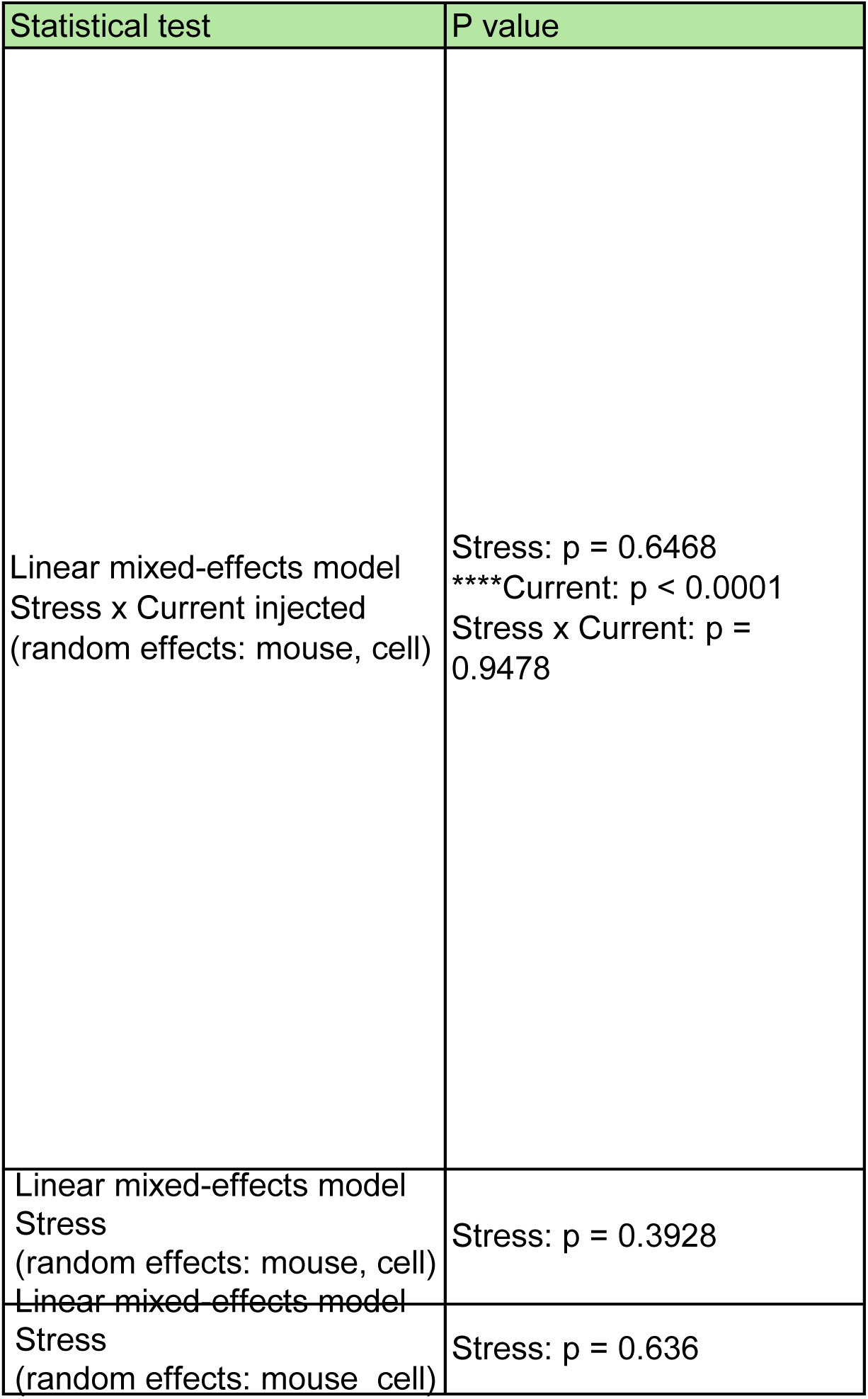
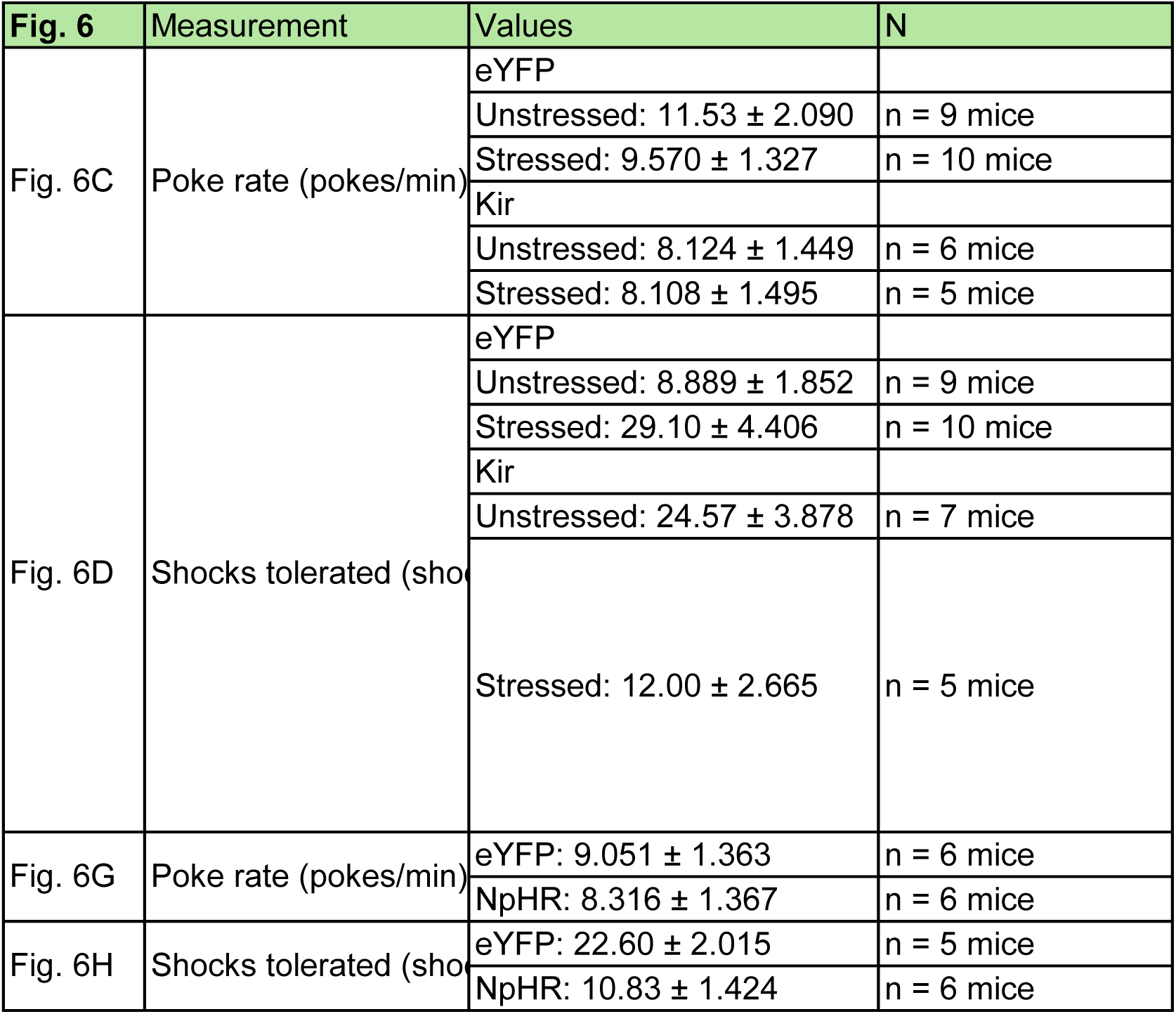
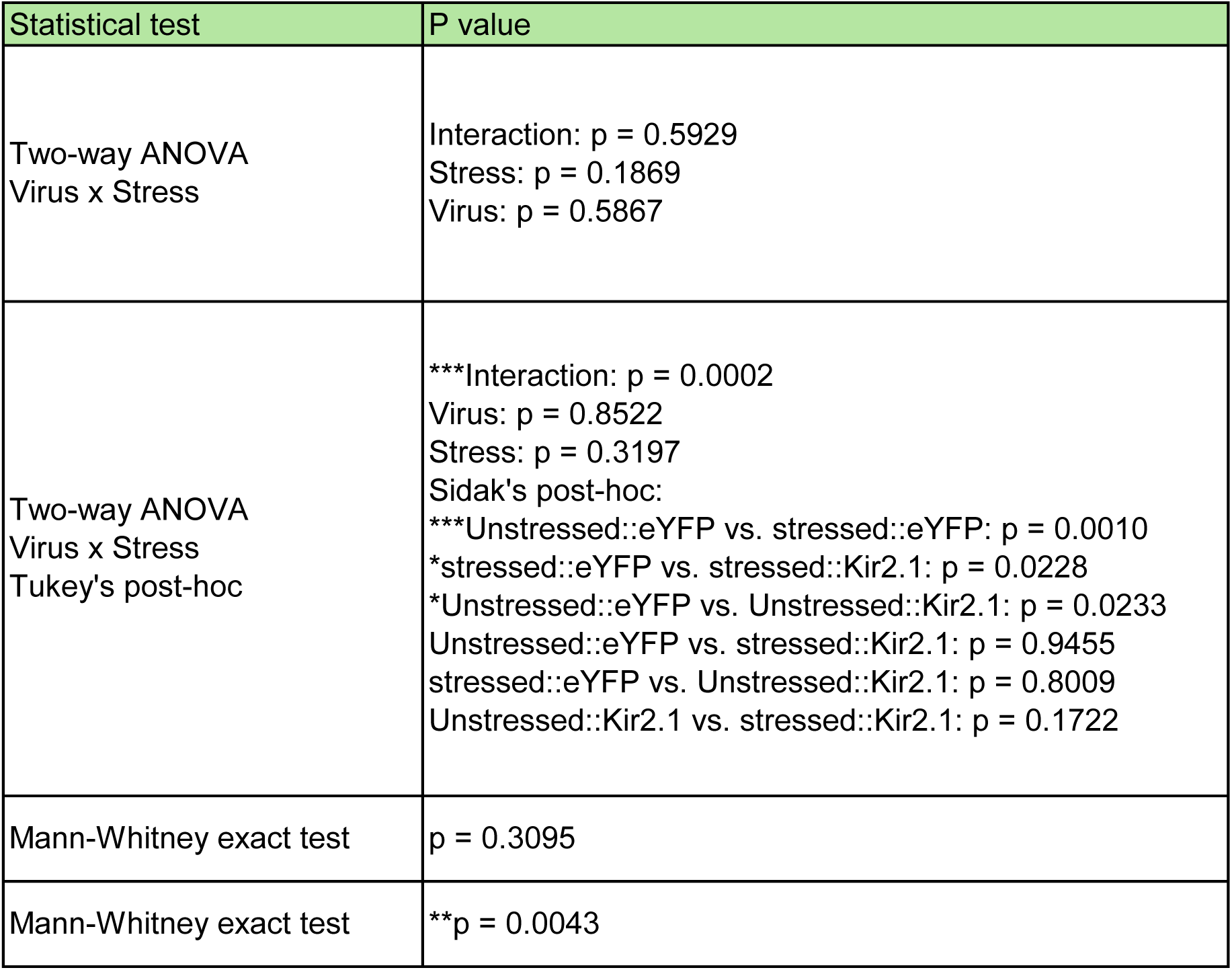
Summary of data and statistical analyses, related to all Figures.

## Notes

### Competing Interest Statement

The authors have declared no competing interest.

